# Social influences complement environmental cues to stimulate migrating juvenile salmon

**DOI:** 10.1101/2024.06.14.598894

**Authors:** Maria Kururvilla, Thomas P. Quinn, Joseph H. Anderson, Erika M. Miller, Andrew G. Berger, Connie Okasaki, John R. McMillan, George R. Pess, Peter A.H. Westley, Andrew M. Berdahl

## Abstract

**Background:** The large-scale seasonal migrations undertaken by many species require complex navigational and timing decisions. Animals migrating in groups might benefit from collective decision making, especially if the environment is noisy (i.e., has high degree of local variation rather than smooth gradients in, for example, salinity or temperature), unpredictable, or the migrants cannot rely on individually acquired information. We focus on juvenile salmon whose migration from fresh water to the ocean is timed to match suitable conditions for growth and survival. While the environmental and physiological factors that influence the timing of migration have been well studied, the influence of social interactions on migration timing is poorly understood.

**Method:** We compiled juvenile salmon data, collected at trap over 19 years, during their downstream seaward migration in three rivers in Washington state along with a suite of relevant environmental time series. We developed state space statistical models to estimate the influence of hatchery-produced salmon to stimulate the downstream migration of wild salmon, while also incorporating potential environmental stimuli.

**Results:** Our results are consistent with the “pied-piper” hypothesis that large numbers of migrating hatchery-origin salmon provide a social cue stimulating migration of co-occurring wild salmon. The number of hatchery salmon counted at the trap was a strong predictor of the number of wild sub-yearling Chinook salmon in the Dungeness and Puyallup rivers and on yearling coho salmon in the Puyallup and Skagit rivers. Migration timing was also influenced by a suite of physical factors related to temperature, river flow, photoperiod, and lunar phase.

**Conclusions:** Our findings highlight the potential for social cues to affect migration timing of downstream migrating salmon, in concert with environmental factors. Incorporating social information into timing decisions may allow animals to benefit from collective decision making strategies and better time their migrations. Moreover, understanding the effects of large-scale hatchery releases on wild salmon migration may provide valuable insights for planning the timing and duration of hatchery releases.

## Introduction

From monarch butterflies (*Danaus plexippus*) to blue whales (*Balaenoptera musculus*), many species undertake large-scale seasonal migrations to take advantage of predictable changes in abiotic and biotic conditions [1]. These migrations often involve challenging decisions regarding where and when to go. Such decisions must often be made based on noisy (i.e., locally variable) information that only weakly correlates with the conditions in distant locations the animals need to infer [2,3]. For example, long-distance migrants like the East Atlantic light-bellied brent goose (*Branta bernicla hrota*) need to infer the timing of the onset of spring in the Arctic based on conditions in the temperate winter staging area [4]. Some animals can learn from previous experience [5] but first-time migrants may not be able to determine conditions in the destination until they arrive, and in some species cannot follow their parents or experienced migrants. Collective decision-making – where individuals synchronize movements and/or migrations partly based on the collective behaviors of the larger group – may improve the accuracy of decisions based on noisy information and migrating animals in groups potentially benefit from collective decision making [6,7]. For example, the homing efficiency of homing pigeons (*Columba livia*) improved when flying in flocks [8,9]. Even migrations that appear to be asocial – such as the highly dispersed migrations [10] of monarch butterflies [11] or blue whales [12] – can be influenced by social information, through indirect cues such as visual or olfactory cues or long-range communication [13,14]. While most of the research on collective decision making during migration has been focused on deciding where to go (*i.e.*, collective navigation) interest is expanding to include another critical question migrants face: when to go [15,16].

Pacific salmon and trout (*Oncorhynchus* species) migrate as juveniles from freshwater to the ocean, and as adults return from the ocean back to their natal site to spawn [17]. Pink (*O. gorbuscha*) and chum (*O. keta*) salmon fry migrate immediately or almost immediately to sea after emerging from gravel nests but other species spend months or years feeding in freshwater habitats before migrating [17]. Chinook salmon (*O. tshawytscha*) display a range of juvenile life history and migration phenotypes. Ocean-type juveniles migrate to sea in their first year of life, shortly after they emerge from the gravel or after a few months of feeding in streams, whereas stream-type juveniles migrate in the spring after feeding for a full year in the stream [17,18]. Interestingly, species vary in social behavior as juveniles prior to seaward migration. Downstream migration is characteristically at night [19] and schooling is difficult to observe directly, but experiments [20,21] and field observations [22] indicate strong schooling by pink and chum salmon fry. In contrast, juvenile coho salmon (*O. kisutch*), anadromous rainbow trout, known as steelhead (*O. mykiss*) and coastal cutthroat trout (*O. clarkii clarkii*) typically establish and defend feeding territories in streams [23–26]. The ocean-type and stream-type variants of Chinook salmon also differ in their levels of aggression and territoriality; those feeding for longer periods in streams are more strongly territorial than those leaving after a shorter period of time [23,27]. However, the shift of this territorial behavior and upstream orientation to more aggregative behavior and downstream swimming is a key part of the transformation from stream-resident parr to seaward migrating, saltwater-tolerant smolts [28].

The smolt migration to sea is timed, on a broad level, to match arrival in salt water with optimal conditions for growth and survival in estuaries and at sea [3,29,30]. These conditions, including sea surface temperature and spring plankton bloom, vary from year to year [31,32]. Consequently, the timing of arrival of juvenile salmon in marine waters affects their survival. For example, earlier migration has been associated with higher survival in both hatchery and wild Chinook salmon, whereas later migration has been associated with higher survival in hatchery coho salmon [33,34]. Photoperiod, combined with internal circannual rhythms, provides the primary cue for the complex changes in physiology and endocrine systems needed to prepare the fish for the radical change in osmotic environments and habitat [35,36]. The behavioral changes leading to the transformation from upstream-oriented territorial parr to downstream-migrating, more aggregative or schooling smolts [21] are much less well known than the physiological processes [28].

The importance of salmon in commerce, culture, and ecosystems in the northern Pacific Ocean (and Atlantic Ocean as well for *Salmo salar*, the Atlantic salmon) [37] has led to many long-term monitoring programs to assess the abundance of seaward migrating smolts and returning adults [38], and efforts to determine the environmental stimuli that trigger them [39–41]. Daily counts of migrating juvenile salmon from traps typically show a series of peaks and troughs rather than the smooth, more or less bell-shaped curve seen when data from many years are pooled. Efforts to model these day-to-day patterns have tended to focus on physical features of the rivers and especially aspects of water temperature and river flow, and lunar phase, typically with only modest success [42]. Considerable variation is left unexplained after various combinations of physical data are included in models. However, this ruggedness in count data from the spawning migrations of adult salmon have been replicated by models that include social behavior [43]. Therefore, social interactions may also be important in the timing of migrations of salmon smolts.

Hatcheries, used for many decades to bolster salmon populations, may incidentally provide a way to test for the use of social information in the migration timing decisions of juvenile salmon. Hatcheries rear young salmon in mortality-reduced conditions, before releasing them into rivers where they migrate to the ocean. Due to a combination of release strategies, physiological readiness to migrate and the abundance of hatchery-origin juveniles, these releases often greatly increase the number of fish in the rivers over a brief period of time. Data from traps reveal that hatchery-produced salmon often migrate rapidly, and *en masse*, to the ocean (Figure 1). Trap operators and biologists have hypothesized that these large numbers of hatchery-origin fish may provide a social cue to migrate for the wild conspecifics (personal communication with Peter Topping), which typically have a much more protracted migratory window [44,45]. Efforts to test this so-called “pied-piper” hypothesis (referring to the folktale of the pied piper of Hamlin) have yielded supportive [42] or equivocal results [46].

With respect to migrating juvenile salmon, the pied piper hypothesis proposes that fish that are more or less physiologically ready, but have not yet begun to migrate may experience other conspecifics migrating downstream past them through visual, odor or lateral line cues. Due to social attraction, the former may be influenced to join the latter, increasing the social migration stimulus for downstream fish. Once this bolus of smolts has departed, there are fewer fish ready to migrate, and even ideal physical conditions do not result in large numbers of migrants, until the remaining fish increase in readiness and the process repeats itself until the population has all migrated. This conceptual model of migration in wild populations thus incorporates social with physical factors triggering migration, the result being a migration characterized by pulses of migrants even in the presence of smoothly varying potential environmental triggers for migration. The null hypothesis is that wild salmon only respond to environmental factors and do so independently of conspecifics.

In this paper, we leverage a multidecadal dataset to test for the use of social information in the timing decisions of migrating salmon smolts of two different species. Specifically, we use a state space statistical model to estimate the influence of hatchery-produced coho and Chinook salmon (*O. tshawytscha*) smolts on the movement decisions of wild-origin smolts, relative to established physical factors affecting movement (Table 1). A state space model can allow us to estimate the effect of covariates (both environmental and social) on the number of wild salmon migrating after accounting for process errors, observation errors, and autocorrelation. Chinook and coho salmon are widely propagated in the region’s hatcheries [47], and transition from stream-resident to migratory behavior after several months to a year [17], and thus are suitable for testing our hypothesis. While augmenting our view of social decision making to include temporal aspects of migration, this work also contributes to our knowledge of salmon life history and our understanding of the unintended ecological impacts of hatcheries on wild salmon populations [48,49].

**Figure 1.**
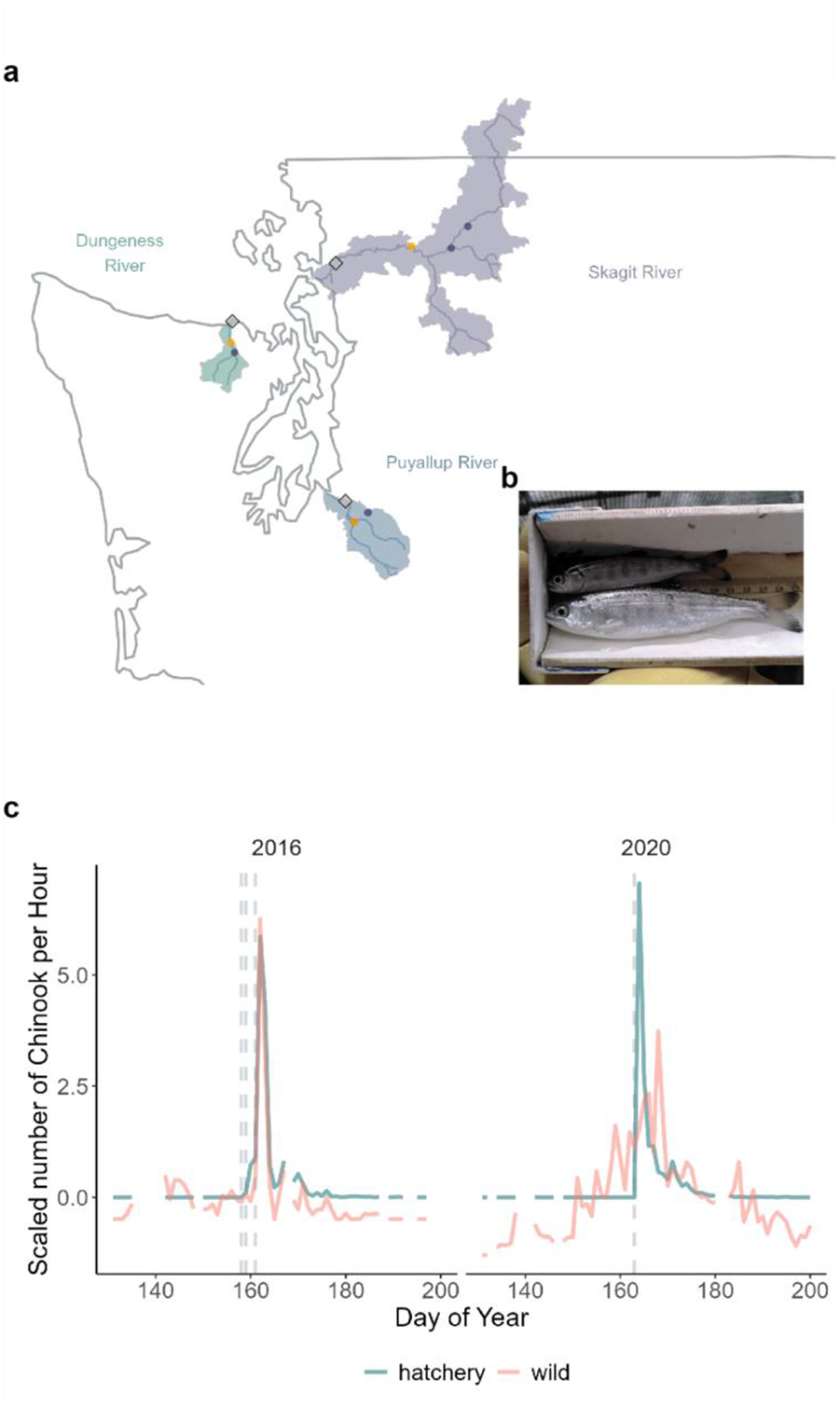
a) Map of the rivers (lines) included in this study, along with their watersheds (polygons) in Washington State, USA. Black diamonds indicate the location of the smolt traps. Purple and orange circles indicate common Chinook and coho salmon release sites respectively. b) Image of a wild (top), and a hatchery (bottom) yearling coho salmon caught in the Dungeness River trap. c) Scaled values of number of Chinook salmon per hour caught in the trap in the Dungeness River from 2016 and 2020. Gray vertical dashed lines indicate the date of hatchery release(s). In some years (*e.g*., 2016), many wild salmon migrate past the trap concurrent with the pulse of hatchery fish. In other years (*e.g.*, 2020), this correlation is not clear.

## Methods

### Trap Data

To estimate the influence of hatchery salmon on the migration of wild salmon, we focused on species in three rivers in Washington state with large hatchery programs propagating sub-yearling Chinook salmon (hereafter Chinook salmon) and yearling coho salmon (hereafter coho salmon) (<u>Table S1</u>). We refer to hatchery-origin salmon as hatchery salmon and natural-origin salmon as wild salmon, acknowledging that fish termed “wild” may have recent hatchery ancestry (*i.e*., one or two hatchery-origin parents spawning in the river). Other life history variants such as yearling Chinook salmon and sub-yearling coho salmon are much less numerous and were excluded from our analyses. We used data from salmon traps deployed in the Dungeness (years 2005-2020), Puyallup (years 2004-2021), and Skagit (years 2010-2022) rivers to quantify juvenile salmon populations migrating to sea. These three rivers are representative of those in which coho and Chinook salmon naturally occur in the region, but are not replicates of each other. The Skagit is the largest in terms of watershed area, followed by the Puyallup, and the Dungeness is the smallest (Figure 1), and the extent of human habitat modification is greatest in the Puyallup River. We chose them primarily because they have long-term records of migrating hatchery and natural origin salmon. Average number of releases per year and average number of salmon released per year in each river is shown in <u>Table S14</u>.

Migrating smolts are caught in a single floating screw trap (in the Dungeness and Puyallup rivers) or in both a floating scoop trap and a floating screw trap (in the Skagit River), counted and released [50]. The hatchery origin salmon were identified by trained staff operating the traps by the presence of an internal coded wire tag (detected magnetically), or a clipped adipose fin. While most hatcheries try to mark all the salmon that are released in the rivers, a small percentage (<5% of hatchery coho salmon in the Dungeness and Skagit rivers; <5% of hatchery Chinook salmon in the Skagit and Puyallup rivers; <12% of hatchery Chinook salmon in the Dungeness River; <5% of hatchery coho salmon in the Puyallup River except for two instances (2009 and 2021) with >50%) were unmarked (unclipped and untagged). The unmarked hatchery salmon are comparable in abundance to the wild salmon present in the Dungeness and Puyallup rivers. Because hatchery salmon are often bigger than wild salmon and have a different appearance (Figure 1), the trap operators sometimes distinguish between the hatchery salmon and wild salmon and such fish are recorded as ‘unmarked hatchery’; however, this categorization may not always be accurate. In the Puyallup River, the unmarked hatchery Chinook salmon were accounted for in the past seven years by subtracting the number of unmarked hatchery salmon estimated by Regional Mark Information System (RMIS) from the number of wild salmon caught. We addressed the issue of misidentification of unmarked hatchery salmon as wild salmon at the traps in all the rivers by using data from RMIS to get the proportion of unmarked hatchery salmon released by the hatcheries to look at the correlation between the number of unmarked hatchery salmon released and the number of wild salmon caught in the traps (<u>Figure S1</u>). A strong relationship would suggest wild catch totals were mostly driven by unmarked hatchery fish, potentially confounding our counts of wild salmon. We found no correlation between the number of unmarked hatchery salmon and the number of wild salmon for any river-species pair, except Skagit River coho salmon, which had a moderate correlation (0.36). Therefore, we use unmodified hatchery counts in our analysis. Because the proportion of unmarked hatchery coho salmon in the Puyallup River was unusually high in 2009 and 2021 (>50%), we reran our analysis without those years and found similar results (<u>Figure S8</u>).

In all three rivers, some Chinook salmon migrate shortly after emergence in January to March (fry migration) and others migrate later, from May to July, after feeding in the river for several months (parr migration), as is typical in the region [42,43]. This results in a strongly bimodal distribution of migration times. Since the hatchery smolt releases coincided with the later parr migration, we restricted our analysis to days of the year 130 - 200 in the Dungeness River and days of the year 130 - 218 in the Puyallup River (<u>Figure S1</u>). In the Skagit River, the proportions of fry migrants and parr migrants vary from year to year [44]. Most of the sub-yearlings migrated as fry in the years of data we had, leaving only a small proportion of the migrants remaining during hatchery releases. The Skagit River trap was only operated up to day 189 in some years. Therefore, we used only data from days 150 - 189 for Chinook salmon in the Skagit River. For coho salmon, we used data between days 120 and 160 in the Dungeness River, between days 90 and 160 in the Puyallup River, and between days 100 and 150 in the Skagit River.

In the Dungeness and Puyallup Rivers, salmon caught in the trap were counted roughly twice a day – at around 06:00 h and 18:00 h. ‘Night’ counts tended to be much higher than the ‘day’ counts (<u>Figure S2</u>) because the juvenile salmon tend to migrate at night. In the Skagit River, the trap operated only during the nights, except for every third day, when the trap operated all day, but was checked twice. Because two thirds of the ‘day’ counts were not enumerated, we only analyzed the ‘night’ counts in the Skagit River. The counts were separated by scoop trap counts and screw trap counts in the Skagit River. In all rivers, because the time at which the trap started and stopped operating varied slightly each deployment, we used the time deployed to calculate the catch per unit effort or catch rate for each day or night by dividing by the number of hours deployed.

We log-transformed the number of wild salmon (after adding 1 to each data point to avoid infinite values) because the data were log normally distributed. We scaled (divided by the standard deviation) and centered (subtracted the mean) the data for each year to account for large year-to-year variation in population size so that we could compare between the years. We scaled, but did not center, the catches of hatchery salmon counts so that the counts remained 0 when no hatchery fish were caught (for example, Figure 1c). Immediately after a hatchery release, the traps sometimes get inundated with hatchery salmon. To avoid this, trap operators sometimes suspend operations for a few days, resulting in missing data. We linearly interpolated the hatchery data for days when the trap was out since the analysis that we used does not allow missing values in the covariates. We interpolated around 16% (526 values) of the data from the Dungeness River, 37% (1148 values) of the data from the Puyallup River and 13% (267 values) of the data from the Skagit River. Most of the missing values in the Puyallup and Skagit rivers were towards the end of the season and during the Covid-19 lockdown in 2020. Because missing values are allowed in the response variable of our model, we did not interpolate missing values of the wild salmon counts.

### Environmental Data

We aimed to estimate the social effects on the migration of salmon smolts, while controlling for any environmental effects. Based on previous studies [39,51–53], we selected ten environmental covariates (Table 1) that have been reported to influence Chinook [39,51,53] and coho salmon smolt migration [52,53] in British Columbia [51,53] and Oregon [39,52]. Although this is not a comprehensive review of the literature on environmental effects on salmon smolt migration, and there may be differences in the results due to geographic variations [52], we hypothesized that these covariates would affect the downstream migration of the salmon smolt species included in our study. All the environmental data that we used were collected in the same river basin as the migrating salmon. While these data do not capture the full range of environmental conditions that the fish experience throughout the spatial extent of juvenile rearing habitats, we assumed that measured water/air temperature and discharge downstream were correlated to the water temperature and discharge experienced by the fish.

For the Dungeness River, we used flow data from the Washington Department of Ecology (Dungeness River mouth, Station ID 18A050). For the Skagit River, we used flow data from USGS (Newhalem, Site no. 1217800). For the Puyallup River, we used flow data from USGS (Alderton site, Site no. 12096500).

For the Skagit River, we obtained water temperature data from USGS (Site no. - 12178000). Water temperature data was not available for the Puyallup River so we supplemented this with air temperature data from USDA Natural Resources Conservation Service (Mowich, 941), a site within the Puyallup drainage. For the Dungeness River, we obtained water temperature data from the Washington Department of Ecology (2004-2014, Station ID 18A050) and from the Washington Department of Fish and Wildlife (2016-2020) and calculated the accumulated temperature units (ATU) as the cumulative sum of the daily mean water temperature from the earliest date that was sampled every year (April 1st). In the Skagit/Puyallup rivers, we calculated ATU as the cumulative sum of the daily mean water/air temperature from the Winter Solstice of the previous year (December 21st). Even though no catch data were used prior to May 10, temperatures prior to this date affect growth and hence migration decisions after May 10.

We scaled, and for some variables centered, the environmental data for each year to compare the effects of all the covariates (Table 1). Since our analysis did not allow missing variables in the covariates, we linearly interpolated the data for the days in which the data were missing. Before using the environmental data as covariates in the model, we calculated the correlation between every pair of covariates (<u>Figure S3, Figure S6</u>, and <u>Figure S10</u>). If the correlation of any pair was > 0.5, we did not use those covariates in the model at the same time. Instead, we determined the covariate in each pair that had more support from the data (with lower AICc value) and then used those covariates for subsequent analysis.

**Table 1.** List of environmental covariates, source of data, description, data transformation, predicted effect on number of wild salmon migrating (positive or negative) of the covariates used in MARSS analysis for each river. The reference column provides an example of a paper that has used the given covariate in a model of juvenile migration timing of Chinook, and coho salmon in various rivers. (–) indicates no reference but we hypothesize that the covariate would have an effect on migration timing of juvenile salmon.

| Covariate | Data Source | Description | Transformation | Predicted effect (+/-) | Reference |
| --- | --- | --- | --- | --- | --- |
| Temperature | WA Department of Ecology, WDFW, USGS, USDA Natural Resources Conservation Service | Daily mean of water temperature (Skagit River) or water temperature measured at a time point closest to the midpoint of the daily trapping duration (Dungeness River) or daily mean of air temperature (Puyallup River) | Scaled and centered | + | [39,52] |
| Temperature difference | Calculated | Change in water/air temperature between that day and previous day $x_t - x_{t-1}$ | Scaled | + | [52] |
| ATU (accumulated thermal units) | Calculated | Cumulative sum of water/air temperature starting on the winter solstice of the previous year (Skagit and Puyallup rivers) or April 1st (Dungeness River) | Scaled and centered | + | [51] |
| Temperature residuals | Calculated | Residual effect of temperature after controlling for photoperiod in a linear model | Scaled | + | — |
| Photoperiod | sunrise-sunset.org | Number of daylight hours for a latitude, longitude near the trap | Scaled and centered | + | [39,52] |
| Photoperiod difference | Calculated | Change in photoperiod between that day and previous day $x_t - x_{t-1}$ | Scaled | + | — |
| Flow | WA department of Ecology, USGS | Daily mean of water flow (Skagit River), or daily flow measured at a time point closest to the midpoint of the trapping duration (Dungeness River), or daily flow measured when the trapping started (Puyallup River) | Scaled and centered | - | [51] |
| Flow difference | Calculated | Change in water flow between that day and previous day $x_t - x_{t-1}$ | Scaled | + | [52] |
| Lunar phase | Ephem package, python | Percentage of the moon illuminated | Scaled and centered | - | [39,52] |
| Secchi depth | Puyallup Tribe of Indians | Water clarity measured with a Secchi disk when the trapping started (only available for Puyallup River) | Scaled and centered | - | [53] |

### Analysis

We used multivariate autoregressive state-space (MARSS) models to estimate the influence of the number of hatchery salmon migrating downstream as determined by the number of hatchery salmon caught in the trap, and of the different environmental variables, on the number of wild juvenile salmon out-migrating [54,55]. MARSS models allowed us to separate the underlying process from the observed data. In the MARSS framework, *y_t_* is the *n* × 1 vector of the log of the observations of wild salmon per hour on the *t*th day of the year for *n* years and *x*_*t*_ represents the *n* × 1 vector of the log of the true, but unknown number of wild salmon per hour on the *t*th day of the year.

They are related by the following equations:

Observation model -

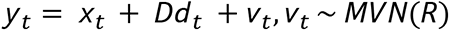

Process model -

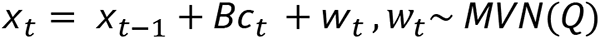

where *c_t_* are the *p* × 1 vector of covariates affecting the states or the average number of salmon per hour migrating past the trap, *p* is the number of covariates multiplied by the number of years, *B* is the *n* × *p* matrix of coefficients relating the effects of the covariates *c_t_* to the states *x_t_*, *d_t_* is the vector of covariates affecting the observations, *D* is the *n* × *p* matrix of the effects of the covariates on the observation. *w_t_* and *v_t_* are the *n* × 1 vectors of normally distributed errors. *R* is the *n* × *n* variance-covariance matrix of the observation errors and *Q* is the *n* × *n* variance-covariance matrix of the process errors. We assumed that the observations and states were independent between years and that the observation and process variances were the same for all years. Therefore, *R* and *Q* are diagonal matrices.

We separated the data into day and night for the analysis and then fit two models: one with equal process variances and the other with unequal process variances for day and night. The effect of the environmental covariates was assumed to be equal for all years and for day and night, whereas the effect of hatchery was equal for all years but allowed to be different for day and night. We did this to test the hypothesis that the effect of hatchery salmon would be higher during the day because the wild salmon might have stronger visual cues from the presence of hatchery salmon during the day [41].

To ensure that the errors met the model assumptions, we plotted the residuals of the predicted values against the fitted values, the autocorrelation function of the residuals, as well as quantile-quantile plots. We assumed that both the process errors and observation errors are identical, independent, and normally distributed within years.

The MARSS framework allowed us to address an alternate hypothesis to the pied piper: that hatchery and wild salmon are *independently* responding to the same environmental stimuli. For example, if both hatchery and wild smolts independently respond to the river flow, peaks in their respective migration timing distributions would coincide. However, in this scenario, a high number of wild smolts would be correlated with high flow when there are no hatchery smolts migrating downstream as well as when there are many hatchery smolts migrating. Therefore this model would estimate the coefficient of flow to be positive and the coefficient of the hatchery (catch rate of hatchery smolts) covariate to be zero and not falsely support the pied piper hypothesis.

#### Model Selection and Relative Importance of Variables

Akaike Information Criterion (AIC) is often used to select the best model among models with different combinations of covariates. However, with autocorrelated data, the sample size is much lower than the number of data points we have. Thus, we used AICc, which is AIC with a correction factor for small sample size, to compare models.

We first compared the model with equal process variance for day and night with the model with unequal process variance for day and night and used AICc to select the most appropriate variance structure in all six cases.

Among the models with each of the correlated covariates, the model with photoperiod difference had the lowest AICc for Chinook salmon in the Dungeness and Puyallup rivers (Table S2, S6). In the Skagit river, the model with temperature residuals had the lowest AICc for Chinook salmon (<u>Table S10</u>). For coho salmon in the Dungeness River, the model with photoperiod had the lowest AICc among all the correlated covariates (<u>Table S2</u>). On the other hand, the model with ATU had the lowest AICc among the correlated covariates for coho salmon in the Puyallup River and the Skagit River (<u>Table S10</u>). We then fit each model for each river-species combination with every combination of uncorrelated covariates (including the hatchery covariate) and chose the model with the lowest AICc to get the estimates of the effects (*B*) and model diagnostics.

In addition to estimating the best model from model selection, we calculated the relative variable importance as the probability that the given variable is in the best model. We estimated the probability by calculating the Akaike weight of each model and then summing the weights of all the models for which that variable was present. We consider a variable ‘important’ if it has at least 90% probability of being in the best model. While this method does not describe the significance of the variable or the size of the effect that the variable has on the response variable, it describes the relative importance of the covariate in the set of candidate models.

## Results

### Social Effects

To test the pied piper hypothesis, we looked at the following three criteria. One, whether the hatchery covariate was in the top model(s) (*i.e.*, the model with the lowest AICc contained the hatchery covariate). The hatchery covariate being included in this top model indicates that considering the number of hatchery fish out migrating allows for better prediction of smolt outmigration than considering only environmental variables (discussed in following section). Two, the effect size and significance of the hatchery covariate. A positive effect means that more wild fish migrated when the hatchery fish migrated, even accounting for environmental cues that could cause them both to migrate independently. The significance means that the estimated range of effect sizes did not include zero. Three, the relative importance of the hatchery covariate – described in the Methods section. The importance measures the role of a variable across all potential model fits and we used a threshold of ≥0.9 to indicate high relative variable importance.

### Chinook

#### Dungeness & Puyallup rivers

The hatchery covariate was included in the top AICc scoring model in the Dungeness and Puyallup rivers. The effect of hatchery salmon on wild salmon had a positive effect in both day and night in both rivers (Figure 2a<u>, 2b</u>). The estimated effect size was equal for both day and night, meaning we found no difference between the social effect during day and during night.

The hatchery covariate was an important variable in both rivers compared to all other variables we tested in model selection (Table S4, S9), as it is in all the top models.

#### Skagit River

While the best model for Chinook salmon in the Skagit River did not have the hatchery covariate, a model with equal support (*ΔAICc* < 2, <u>Table S12</u>) did. The effect was positive, but the confidence intervals overlapped with 0 (Figure 2c). The hatchery covariate was not an ‘important’ variable in the Skagit River compared to all other variables we evaluated in model selection (<u>Table S11</u>).

### Coho

#### Dungeness River

In the Dungeness River, while the hatchery covariate was not in the best model, the model with this covariate had equal support (*ΔAICc* < 2, <u>Table S5</u>). The effect of the hatchery covariate was positive during the day and negative during the night and the confidence intervals for both estimates overlapped with 0 (Figure 3a). The hatchery covariate was not an ‘important’ variable in the Dungeness River compared to all other variables we tested in model selection (<u>Table S4</u>).

#### Puyallup & Skagit rivers

In both the Puyallup and Skagit rivers the hatchery covariate was included in the top scoring model (Figure 3b,c). In both rivers, we found a strong positive effect of the hatchery covariate on the number of wild salmon migrating (Figure 3b,c). Further, the hatchery covariate was an important variable in the both rivers compared to all other variables we tested in model selection (<u>Table S11</u>).

#### Social Summary

For both species, we found that increases in the number of wild salmon migrating coincided with increases in the number of hatchery salmon migrating in two out three rivers. For Chinook salmon in the Skagit River, the estimate of the effect was also positive, but did overlap zero. This consistency is on par with the consistency of ‘known’ environmental effects explored in the following section. In summary, although our results are correlative, the positive effect of hatchery smolts on the migration of wild smolts is consistent with the results we would expect under the pied piper hypothesis – wild smolts being influenced to migrate when they observe many hatchery smolts outmigrating.

### Environmental Effects

The aim of this manuscript is to test the pied piper hypothesis, therefore we focus on the social variable. However, a shared response to the same environmental variable can be mistaken for social influence, so we also included environmental variables in our models to account for potential common cues shared by hatchery and wild salmon smolts. Below we report the environmental variables found in the models for our social analysis.

### Chinook salmon

#### Flow & Δ flow

Among all the covariates, flow difference or increase in the river’s flow had the strongest positive effect on the number of Chinook salmon migrating in the Dungeness and Puyallup rivers (Figure 2a<u>, 2b</u>). However, absolute flow had a small negative effect in the Puyallup River (Figure 2b). This suggests that when the flows are low, an increase in flow of the river from the previous day was correlated with more salmon migrating downstream. Flow was not a covariate in the top models in the Skagit River.

#### Photoperiod & Δ photoperiod

Photoperiod difference, or increase in day-length, also had a positive effect on the number of Chinook salmon migrating in the Dungeness and Puyallup rivers (Figure 2a,b). Photoperiod had a negative effect on Chinook salmon in the Skagit River (Figure 2c), indicating that as the season progressed and day length increased, the number of salmon migrating increased in two rivers but declined in the third.

#### Temperature, Δ temperature, & temperature residuals

Temperature difference, or change in water temperature from the previous day, had the strongest negative effect on the number of Chinook salmon migrating in the Dungeness and Skagit rivers (Figure 2a, c). The residuals of the model with temperature also had a negative effect on Chinook salmon in the Skagit River. This suggests that a drop in water temperature or unseasonably cooler water was correlated with more Chinook salmon migrating. Air temperature had a strong positive effect on Chinook salmon in the Puyallup river (Figure 2b), suggesting that more Chinook salmon migrated when the air was warmer, though warm air can lead to cool water through snow melt.

#### Lunar phase

Lunar phase, or the percentage of the moon that was illuminated, had a small positive effect (Figure 2b) on Chinook salmon in the Puyallup River, suggesting that Chinook salmon were more likely to migrate during the full moon.

**Figure 2.**
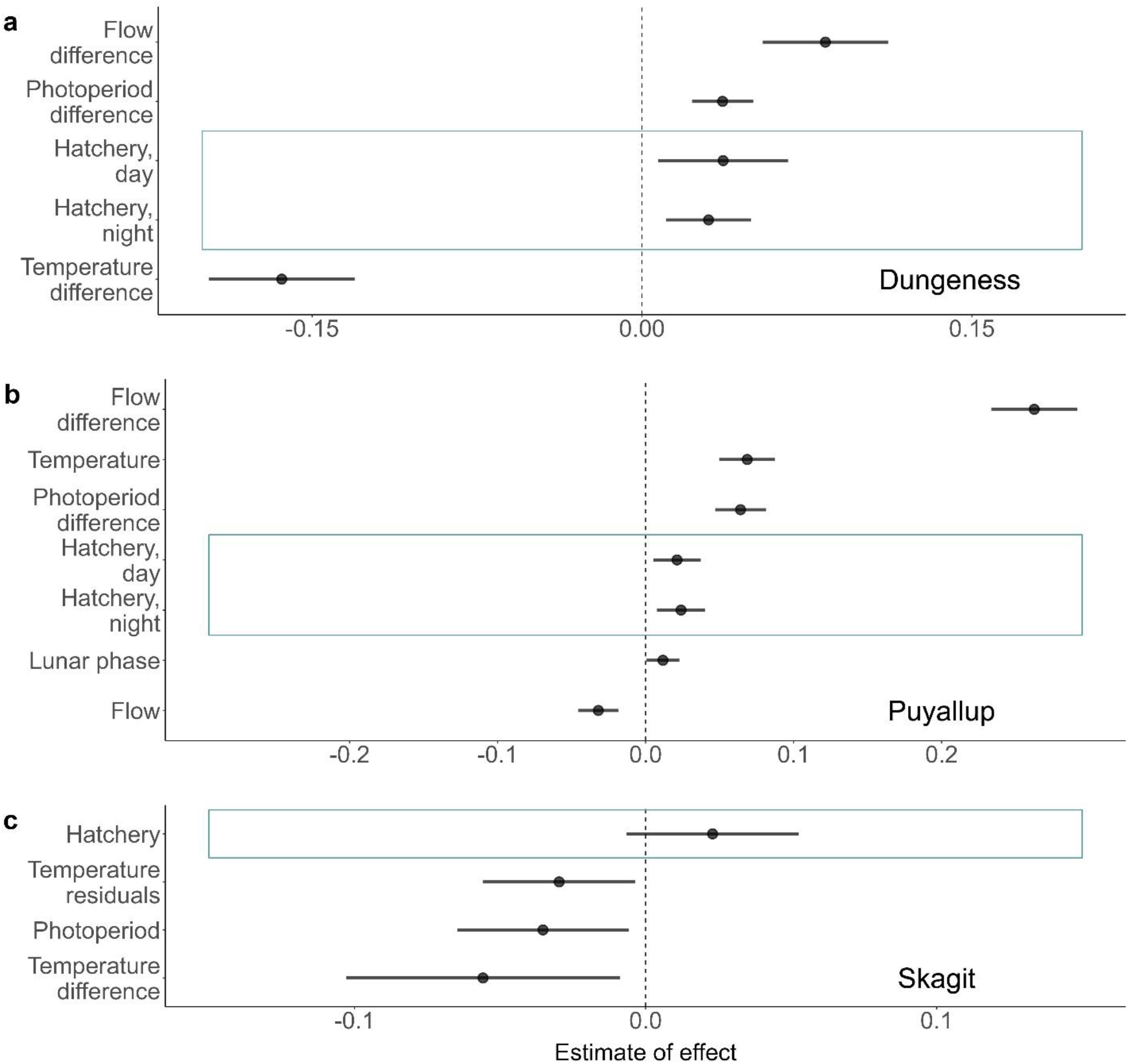
Estimates of the effect of different covariates in the best MARSS model or MARSS model with equal support for the number of wild Chinook salmon in the a) Dungeness, b) Puyallup, and c) Skagit rivers. Blue boxes indicate social variables associated with the pied piper hypothesis.

### Coho salmon

#### Flow and flow difference

Flow difference, or an increase in water flow, had a positive effect on the number of wild coho salmon migrating in the Dungeness and Puyallup rivers (Figure 3a, b), but absolute flow had a negative effect on the number of wild coho salmon caught in all rivers (Figure 3a, b, c). Thus when flows were low, an increase from the previous day was correlated with more wild salmon migrating.

#### Temperature difference

Temperature difference, or change in water temperature from the previous day, had a negative effect on the number of coho salmon in the Dungeness River (Figure 3a). This suggests that more salmon migrated after a drop in water temperature from the previous day.

#### Accumulated thermal units

ATU or accumulated temperature experience, had a negative effect on the number of coho salmon in the Skagit and Puyallup rivers (Figure 3b, c). This suggests that as the season progresses and the temperature experience increases, the number of salmon migrating reduces.

#### Photoperiod

Photoperiod, or the day-length, had a negative effect on the number of coho salmon in the Dungeness River (Figure 3a), but a positive effect in the Puyallup River (Figure 3b).

#### Lunar phase

Lunar phase, had a negative effect on the number of wild coho salmon in the Skagit River (Figure 3c), but it was not a covariate in any of the top models in the Dungeness or Puyallup rivers. This suggests that fewer fish migrated during the full moon in the Skagit River.

**Figure 3.**
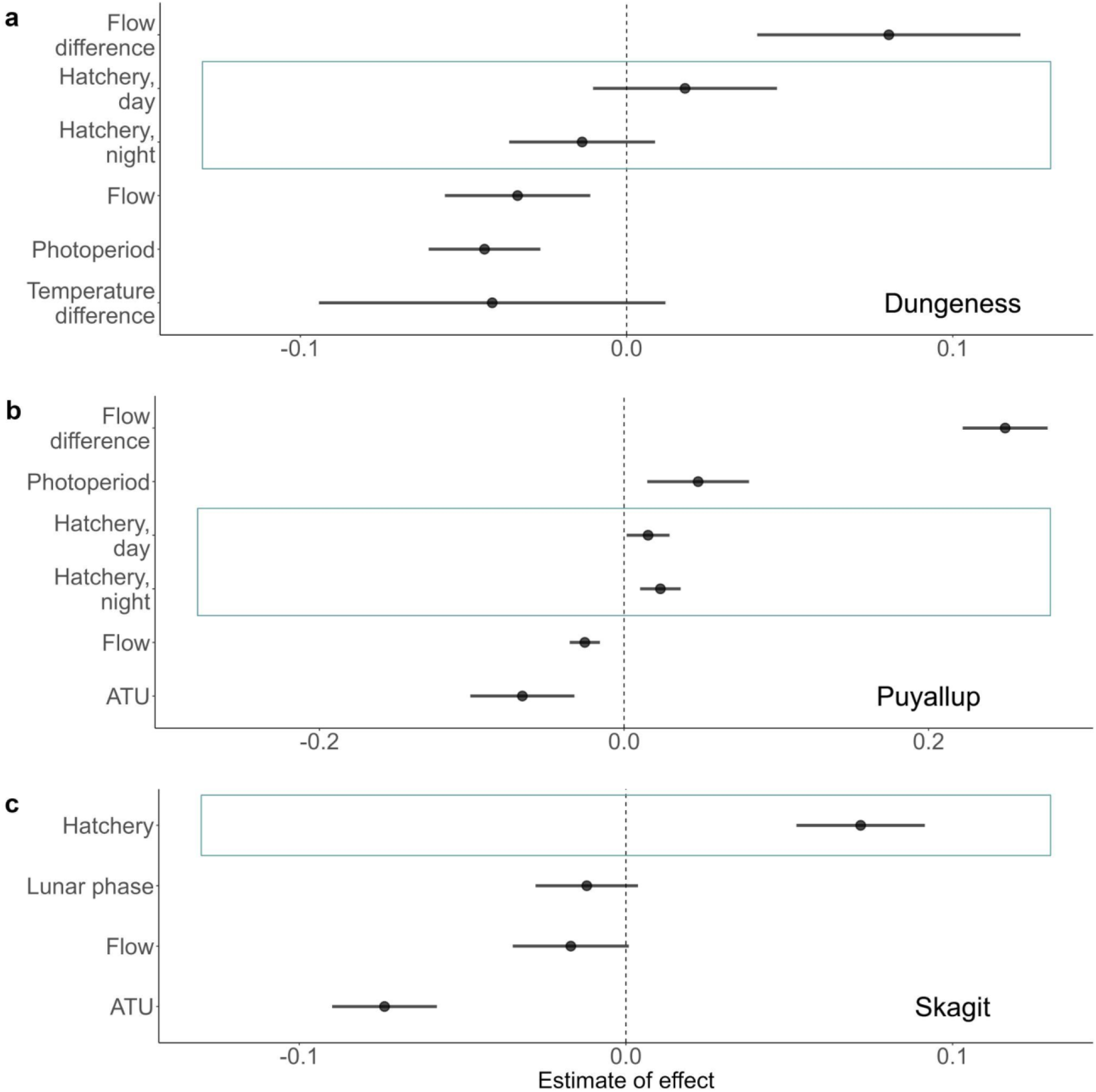
Estimates of the effect of different covariates in the best MARSS model or MARSS model with equal support for the number of wild coho salmon in the a) Dungeness, b) Puyallup, and c) Skagit rivers. Blue boxes indicate social variables associated with the pied piper hypothesis. Puyallup River coho results were similar when we excluded years with anomalously high proportions of unmarked hatchery smolts (<u>Figure S8</u>).

## Discussion

Our main objective was to determine if social information was important in the timing decisions of wild Chinook and coho salmon smolts during their seaward migration within the context of the many physical variables previously linked to migration of these species (Table 1). Wehypothesized migrating hatchery smolts would stimulate wild smolts to join, therefore increasing the number of wild smolts migrating above the level expected based on the contemporary environmental effects. In four of our six river-species combinations - Chinook salmon in the Dungeness and Puyallup rivers and coho salmon in the Puyallup and Skagit rivers - the number of hatchery smolts was an important variable with a positive statistical relationship with the number of wild salmon smolts, consistent with the pied piper hypothesis. Our results are supported by the visual observation that following hatchery releases of Chinook salmon in the Wenatchee River, wild Chinook smolts moved downstream along with the hatchery smolts unless they could not see the hatchery smolts [46]. Thus the weight of the evidence indicated an effect of social interactions with hatchery fish, though its magnitude varied among species and rivers, and the relative effects of different physical factors also varied among species and rivers.

For the Chinook salmon in the Dungeness and Puyallup rivers, and the coho salmon in the Puyallup and Skagit rivers, the results were consistent with the pied piper hypothesis that the numerous hatchery salmon smolts migrating downstream trigger the migration of wild salmon smolts. These social influences, however, are best understood in the context of the river’s environmental conditions (aspects of flow and temperature), time of year including photoperiod change, and lunar phase. In some cases (coho salmon in the Skagit River) the effect size of the hypothesized social influence was comparable to the environmental factors, whereas in other cases (Chinook salmon in the Dungeness and Puyallup rivers, and coho salmon in the Puyallup River) the effect of the social influence was smaller than the effect of the largest environmental factors, but as important. We did not find an effect of social influence in the coho salmon in the Dungeness River and the Chinook salmon in the Skagit rivers. Several factors may have contributed to the absence of detectable social cues in these cases. A large portion of the wild salmon smolts may have migrated before the hatchery fish were released. Alternatively, wild salmon may have relied more heavily on tributaries than hatchery salmon and were thus spatially isolated from the locations of hatchery releases, thereby diminishing the potential for social interaction. Furthermore, some of the wild salmon may have been upstream of the hatchery release sites, and thus not exposed to the hatchery salmon.

We saw a markedly greater correlation between hatchery and wild Chinook salmon movements in the Dungeness and Puyallup rivers than in the Skagit River. When both hatchery and wild Chinook were present, the hatchery fish were much more numerous than the wild fish in the Dungeness and Puyallup rivers (<u>Table S1</u>). In contrast, hatchery and wild Chinook salmon were similar in abundance in the Skagit River. In addition, the Skagit River is much larger than the other two rivers (Figure 1) (*e.g.*, Skagit River flows in May, when many smolts migrate, are ca. 4-5 times those in the Puyallup River and ca. 50 times those in the Dungeness River, based on USGS data), including many spawning and rearing areas that are not on the direct migration route from the hatchery to the trap. Furthermore, the Skagit River encompasses six independent populations of Chinook salmon monitored at the trap, whereas the Dungeness and Puyallup rivers each hold a single population [56]. We expect that all of these factors contribute to the greater influence of hatchery origin Chinook salmon in the Dungeness and Puyallup rivers than in the Skagit River.

We found a stronger statistical effect size of the number of hatchery coho salmon in the Skagit and Puyallup rivers than in the Dungeness River. The Dungeness and Puyallup rivers had similarly high ratios of hatchery to wild coho salmon so that attribute does not explain the result. Some aspect of the distribution of breeding and fry emergence (primarily in the tributaries of the Skagit River rather than the river itself [57]), and their subsequent rearing and movements, presumably contributes to these patterns.

Previous studies reported that temperature experience [51], stream temperatures [39], and temperature residuals [34] strongly influenced the migration of Chinook salmon. Temperature difference and temperature residuals had the strongest (negative) effect on the number of wild Chinook salmon in the Dungeness and Skagit rivers, respectively. Thus a drop in water temperature or unseasonably cooler water seemed to cause Chinook salmon parr to migrate, as can occur when rivers rise with melting snow after an increase in air temperatures or rain on snow events [58]. In line with this expectation, in the Puyallup River, air temperature had a strong positive effect on the number of Chinook salmon migrating. The correlation between a drop in water temperature and the increase in the number of salmon smolts migrating has previously been shown [42,52]. This was contrary to our expectation since warmer temperatures resulted in earlier migration in Chinook salmon [51]. Similarly, we found that ATU had a negative effect on the number of wild coho salmon migrating in the Puyallup and Skagit rivers. Similar patterns of negative effect of degree days were shown in coho salmon in Deer Creek, Oregon [52]. We found that flow difference had the strongest effect on both Chinook and coho salmon, in the Dungeness and Puyallup rivers. However, flow difference was not a covariate in the best model for Chinook or coho salmon in the Skagit River. Perhaps flow is more important in smaller rivers like the Dungeness and Puyallup where smaller changes in flow can have larger effects on the fish. Flow might also be less important in the Skagit River because it has several dams that affect flows, whereas the Dungeness River does not currently have any dams. The Puyallup River has one dam and its floodplain habitat is more degraded than the other rivers. Its flow varies widely, and so an increase in the flow can provide a cue for migration.

Although the data from salmon traps are highly appropriate for the purposes of our study, and collected “blind” with respect to our hypotheses, there remain some limitations to our data that warrant consideration. First, analyses assume that all unmarked individuals are ‘wild’, which we explored by looking at the correlation between the number of unmarked hatchery salmon released and the number of wild salmon caught in the traps (<u>Figure S1</u>). We found only a low-to-moderate correlation between the two, implying that the increase in the number of wild salmon was not exclusively because of the unmarked hatchery salmon. Second, the time scale at which data is collected by the traps might not be the time scale of the social influence we tried to study. If wild salmon respond to the hatchery salmon immediately or within a few hours, the effect of this social influence would be diluted by recording data only every 12 hours. Third, we could not distinguish between wild smolts being led by hatchery fish (pied piper hypothesis) or those wild smolts being displaced – wild smolts leaving their in-river holding areas due to an influx of hatchery fish entering those holding areas – downstream by hatchery smolts. Fourth, although we attempted to account for common environmental cues, we cannot exclude the possibility that both hatchery and wild fish responded to stimuli external to our model, rather than hatchery fish causing wild fish to migrate. Finally, the suspension of trap operations following a hatchery release resulted in the loss of valuable data – in particular data most likely to add support for the pied piper hypothesis – so our results may reflect a lower bound on the effect.

Despite the limitations of the dataset and the potential dilution of the effect caused by missing data and the time resolution of the data, our results suggest that along with environmental covariates, social information appears to provide additional cues for salmon seaward migration. This suggests that future studies, in addition to considering environmental cues, should consider social cues from both hatchery and wild conspecifics while studying migration timing of salmon smolts. If large hatchery releases motivate the wild salmon smolts to migrate with them, this can have fundamental ecological consequences. Wild salmon smolts that are being influenced by hatchery salmon smolts might migrate earlier than they normally would, encounter less than optimal foraging conditions and could encounter more predators, thereby experiencing higher mortality at sea [34]. As a result of early migration, juvenile wild salmon may enter the estuary with reduced saltwater readiness, which can extend residence time, and in turn, juvenile salmon with longer residence times can have higher predation [59]. On the other hand, wild salmon out migrating with the hatchery salmon might benefit from predator swamping [60].

Showing that salmon smolts use social information, especially that from hatchery conspecifics, can have implications for conservation and management of these species that are commercially and culturally important. Salmon hatcheries were set up in response to overfishing, the continued loss of habitat, and other factors decreasing salmon runs and they provide many sociocultural and economic benefits. While the goals of the hatcheries were to either increase abundance for fishing or, later, to promote conservation of imperiled populations, they can have unintended ecological consequences on the wild populations [47,61,62]. Therefore, reforming hatcheries to maximize the benefit and minimize the risk on wild populations is crucial [63]. Our study highlights another avenue by which hatcheries could indirectly impact wild salmon populations. To reduce the impact of hatchery releases on the migration timing of wild salmon, hatcheries could experiment with different release strategies. For example, hatcheries can consider timing their releases towards the end of the migration period of the wild salmon or consider releasing the hatchery salmon downstream of the wild salmon habitat. Some hatcheries are moving to volitional releases, which may spread the migration of hatchery smolts over a longer period of time [64]. This practice may reduce the pied piper effect, if the wild smolts respond to a large threshold number of hatchery smolts, or may increase the pied piper effect, by providing a longer social cue to influence the wild smolts. Our results suggest that managers should monitor changes in wild smolt migration timing patterns as they experiment with release practices.

Our findings, consistent with Pacific salmon smolts using social information when timing migrations, fit two nascent patterns in the field of collective behavior, with possible population-level implications. First, our results support an increasing recognition of the importance of sociality in salmon, from navigation [65,66] to trophic interactions [67]. Second, and more generally, our results expand a growing body of literature documenting the importance of social information on timing decisions in a wide range of taxa [15,43,68–71]. It remains to be seen if, and how, collective timing decisions may improve the accuracy of those decisions [16]. If salmon, and other species, do improve decision making by using social information, especially during fitness-critical events such as migration, reductions in population size are expected to decrease decision making ability and thereby feedback on further population decline, possibly contributing to population crashes [72–74]. We hope our results encourage more studies exploring social influence on timing decisions and the potential population-level implications.

## Declarations

### Ethics approval and consent to participate

Not applicable

### Consent for publication

Not applicable

### Availability of data and materials

The smolt trap data analyzed for this study are owned by the Puyallup Tribe of Indians (PTI) and the Washington Department of Fish & Wildlife (WDFW) and are not publicly available, however the data are available from PTI and WDFW on reasonable request. All the code used to process and analyze the data are available on github:

https://github.com/maria-kuruvilla/pied_piper

https://github.com/maria-kuruvilla/pied_piper_r https://github.com/maria-kuruvilla/pied_piper_MARSS.

### Competing interests

The authors declare that they have no competing interests.

### Funding

This project was financially supported in part by the SeaDoc Society, a program of the Karen C. Drayer Wildlife Health Center, School of Veterinary Medicine, University of California, Davis. MK was supported by the Pacific Northwest Salmon Conservation Fellowship through the H. Mason Keeler Endowment. AMB was supported by the H. Mason Keeler Endowed Professorship in Sports Fisheries Management.

### Authors’ contributions

AMB, TPK, PAHW, JHA, JRM and GRP conceived the study. MK, EMM, JHA, and AGB processed the data. MK, MS and CO designed the statistical analysis. MK performed the statistical analysis. MK, TPQ and AMB wrote the manuscript, with input from all authors. All authors read and approved the final manuscript.

## Acknowledgements

We would like to thank the following for their valuable contributions to this study:

○ Andrew Simmons, Peter Topping, and Josh Weinheimer and many seasonal technicians from WDFW, for operating the trap and collecting the data in the Dungeness River, and for assisting with the trap visit.
○ Jim Repoz, Dean Toba, Clayton Kinsel, Skyler Repoz, and many additional seasonal technicians for collecting the fish trap data from the Skagit River
○ Elizabeth Holmes from NOAA, for assistance with the initial MARSS analysis and model setup.
○ Eric Ward, for insightful feedback on the initial analysis.
○ Julia K. Parish from UW, for offering constructive suggestions to improve the manuscript.

## Notes

https://github.com/maria-kuruvilla/pied_piper_r

https://github.com/maria-kuruvilla/pied_piper_MARSS

https://github.com/maria-kuruvilla/pied_piper

### Competing Interest Statement

The authors have declared no competing interest.

### Summary of Updates

Author names revised; Figure 2 number corrected; Figure 3 number corrected.

## Citations

1. Dingle H. Migration: The Biology of Life on the Move. Oxford University Press; 2014.

2. Visser ME, Gienapp P. Evolutionary and demographic consequences of phenological mismatches. Nat Ecol Evol. 2019;3:879–85.

3. Wilson SM, Buehrens TW, Fisher JL, Wilson KL, Moore JW. Phenological mismatch, carryover effects, and marine survival in a wild steelhead trout *Oncorhynchus mykiss* population. Progress in Oceanography. 2021;193:102533.

4. Clausen KK, Clausen P. Earlier Arctic springs cause phenological mismatch in long-distance migrants. Oecologia. 2013;173:1101–12.

5. Baert JM, Stienen EWM, Verbruggen F, Van De Weghe N, Lens L, Müller W. Resource predictability drives interannual variation in migratory behavior in a long-lived bird. Simmons LW, editor. Behavioral Ecology. 2022;33:263–70.

6. Berdahl A, Kao AB, Flack A, Westley PAH, Codling EA, Couzin ID, et al. Collective animal navigation and migratory culture: from theoretical models to empirical evidence. Philosophical Transactions: Biological Sciences. 2018;373:1–16.

7. Larkin PA, Walton A. Fish school size and migration. J Fish Res Bd Can. 1969;26:1372–4.

8. Biro D, Sumpter DJT, Meade J, Guilford T. From compromise to leadership in pigeon homing. Current Biology. 2006;16:2123–8.

9. Kano F, Sasaki T, Biro D. Collective attention in navigating homing pigeons: group size effect and individual differences. Animal Behaviour. 2021;180:63–80.

10. Guttal V, Couzin ID. Social interactions, information use, and the evolution of collective migration. Proceedings of the National Academy of Sciences. 2010;107:16172–7.

11. Reppert SM, Guerra PA, Merlin C. Neurobiology of monarch butterfly migration. Annual Review of Entomology. 2016;61:25–42.

12. Oestreich WK, Fahlbusch JA, Cade DE, Calambokidis J, Margolina T, Joseph J, et al. Animal-borne metrics enable acoustic detection of blue whale migration. Current Biology. 2020;30:4773–4779.e3.

13. Couzin ID. Collective animal migration. Current Biology. 2018;28:R976–80.

14. Aikens EO, Bontekoe ID, Blumenstiel L, Schlicksupp A, Flack A. Viewing animal migration through a social lens. Trends in Ecology & Evolution. 2022;37:985–96.

15. Oestreich WK, Aiu KM, Crowder LB, McKenna MF, Berdahl A, Abrahms B. The influence of social cues on timing of animal migrations. Nat Ecol Evol. 2022;6:1617–25.

16. Kao AB, Banerjee S, Francisco F, Berdahl A. Timing decisions as the next frontier for collective intelligence [Internet]. Available from: http://arxiv.org/abs/2312.02187

17. Quinn TP. The Behavior and Ecology of Pacific Salmon and Trout. 2nd ed. Seattle: University of Washington Press; 2018.

18. Healey MC. Life History of Chinook Salmon (Oncorhynchus tshawytscha). In: Groot C, Margolis L, Clarke WC, editors. Pacific Salmon Life Histories. Vancouver: University of British Columbia Press; 1991. p. 311–294.

19. Roberts LJ, Taylor J, Gough PJ, Forman DW, De LEANIZ CG. Night stocking facilitates nocturnal migration of hatchery-reared Atlantic salmon, Salmo salar, smolts. Fisheries Management and Ecology. 2009;16:10–3.

20. Hoar WS. The evolution of migratory behaviour among juvenile salmon of the genus *Oncorhynchus*. J Fish Res Bd Can. 1958;15:391–428.

21. McDonald J. The behaviour of Pacific salmon fry during their downstream migration to freshwater and saltwater nursery areas. J Fish Res Bd Can. 1960;17:655–76.

22. Munsch S, Cordell J, Toft J. Fine-scale habitat use and behavior of a nearshore fish community: nursery functions, predation avoidance, and spatiotemporal habitat partitioning. Mar Ecol Prog Ser. 2016;557:1–15.

23. Taylor EB. Behavioural interaction and habitat use in juvenile chinook, *Oncorhynchus tshawytscha*, and coho, O. kisutch, salmon. Animal Behaviour. 1991;42:729–44.

24. Sabo JL, Pauley GB. Competition between stream-dwelling cutthroat trout (Oncorhynchus clarki) and coho salmon (Oncorhynchus kisutch): effects of relative size and population origin. 1997;54.

25. Keeley ER. Demographic responses to food and space competition by juvenile steelhead trout. Ecology. 2001;82:1247–59.

26. Young KA. Asymmetric competition, habitat selection, and niche overlap in juvenile salmonids. Ecology. 2004;85:134–49.

27. Taylor EB. Adaptive Variation in Rheotactic and Agonistic Behavior in Newly Emerged Fry of Chinook Salmon, Oncorhynchus tshawytscha, from Ocean- and Stream-Type Populations. Can J Fish Aquat Sci. 1988;45:237–43.

28. Hoar WS. Smolt transformation: evolution, behavior, and physiology. J Fish Res Bd Can. 1976;33:1233–52.

29. Cooney RT, Allen JR, Bishop MA, Eslinger DL, Kline T, Norcross BL, et al. Ecosystem controls of juvenile pink salmon (*Oncorhynchus gorbuscha*) and Pacific herring (*Clupea pallasi*) populations in Prince William Sound, Alaska. Fisheries Oceanography. 2001;10:1–13.

30. Spence BC, Hall JD. Spatiotemporal patterns in migration timing of coho salmon (*Oncorhynchus kisutch*) smolts in North America. Fleming I, editor. Can J Fish Aquat Sci. 2010;67:1316–34.

31. Trenberth KE, Hurrell JW. Decadal atmosphere-ocean variations in the Pacific. Climate Dynamics. 1994;9:303–19.

32. Downton MW, Miller KA. Relationships between Alaskan salmon catch and North Pacific climate on interannual and interdecadal time scales. Can J Fish Aquat Sci. 1998;55:2255–65.

33. James SE, Doherty B, Cox SP, Pearsall IA, Riddell B. Size and timing of hatchery releases influence juvenile-to-adult survival rates of British Columbia Chinook (*Oncorhynchus tshawytscha*) and coho (*Oncorhynchus kisutch*) salmon. Can J Fish Aquat Sci. 2023;80:700–18.

34. Scheuerell MD, Zabel RW, Sandford BP. Relating juvenile migration timing and survival to adulthood in two species of threatened Pacific salmon (*Oncorhynchus* spp.). Journal of Applied Ecology. 2009;46:983–90.

35. Björnsson BT, Stefansson SO, McCormick SD. Environmental endocrinology of salmon smoltification. General and Comparative Endocrinology. 2011;170:290–8.

36. McCormick SD. Ontogeny and evolution of salinity tolerance in anadromous salmonids: hormones and heterochrony. Estuaries. 1994;17:26.

37. Noakes DJ. Overview of salmon and their ecological and economic importance. In: Woo PTK, Noakes DJ, editors. Salmon Biology, Ecological Impacts and Economic Importance. Nova Science Publishers; 2014. p. 1–10.

38. Link MR, English KK. Long-term, sustainable monitoring of Pacific salmon populations using fishwheels to integrate harvesting, management, and research. In: Knudsen EE, McDonald D, editors. Sustainable Fisheries Management. CRC Press; 2000. p. 667–74.

39. Roper BB, Scarnecchia DL. Emigration of age-0 chinook salmon (*Oncorhynchus tshawytscha*) smolts from the upper South Umpqua River basin, Oregon, U.S.A. Can J Fish Aquat Sci. 1999;56:939–46.

40. Teichert N, Benitez J, Dierckx A, Tétard S, Oliveira E, Trancart T, et al. Development of an accurate model to predict the phenology of Atlantic salmon smolt spring migration. Aquatic Conserv: Mar Freshw Ecosyst. 2020;30:1552–65.

41. Simmons OM, Gregory SD, Gillingham PK, Riley WD, Scott LJ, Britton JR. Biological and environmental influences on the migration phenology of Atlantic salmon *Salmo salar* smolts in a chalk stream in southern England. Freshw Biol. 2021;66:1581–94.

42. Hvidsten NA, Jensen AJ, Vivas H, Oyvind B, Heggberget TG. Downstream migration of Atlantic salmon smolts in relation to water flow, water temperature, moon phase and social interaction. Nordic J Freshw Res. 1995;38–48.

43. Berdahl A, Westley PAH, Quinn TP. Social interactions shape the timing of spawning migrations in an anadromous fish. Animal Behaviour. 2017;126:221–9.

44. Hansen LP, Jonsson B. Downstream migration of hatchery-reared smolts of Atlantic salmon (*Salmo salar* L.) in the River Imsa, Norway. Aquaculture. 1985;45:237–48.

45. Kennedy GJA, Strange CD, Anderson RJD, Johnston PM. Experiments on the descent and feeding of hatchery-reared salmon smolts (*Salmo salar* L.) in the River Bush. Aquaculture Res. 1984;15:15–25.

46. Hillman TW, Mullan JW. Effect of hatchery releases on the abundance and behavior of wild juvenile salmonids. McCall, Idaho: Don Chapman Consultants; 1989 p. 265–85.

47. Naish KA, Taylor JE, Levin PS, Quinn TP, Winton JR, Huppert D, et al. An evaluation of the effects of conservation and fishery enhancement hatcheries on wild populations of salmon. Advances in Marine Biology [Internet]. Academic Press; 2007. p. 61–194. Available from: https://www.sciencedirect.com/science/article/pii/S0065288107530026

48. Claussen JE, Philipp DP. Assessing the role of supplementation stocking: A perspective. Fisheries Management and Ecology. 2023;30:583–91.

49. McMillan JR, Morrison B, Chambers N, Ruggerone G, Bernatchez L, Stanford J, et al. A global synthesis of peer-reviewed research on the effects of hatchery salmonids on wild salmonids. Fisheries Management and Ecology. 2023;30:446–63.

50. Volkhardt GC, Johnson SL, Miller BA, Nickelson TE, Seiler DE. Rotary Screw Traps and Inclined Plane Screen Traps. Salmonid field protocols handbook: techniques for assessing status and trends in salmon and trout populations. Bethesda, MD : Portland, Or: American Fisheries Society ; State of the Salmon; 2007. p. 235–66.

51. Sykes GE, Johnson CJ, Shrimpton JM. Temperature and flow effects on migration timing of Chinook Salmon smolts. Transactions of the American Fisheries Society. 2009;138:1252–65.

52. Spence BC, Dick EJ. Geographic variation in environmental factors regulating outmigration timing of coho salmon (*Oncorhynchus kisutch*) smolts. Can J Fish Aquat Sci. 2014;71:56–69.

53. Melnychuk MC, Welch DW. Habitat-mediated effects of diurnal and seasonal migration strategies on juvenile salmon survival. Behavioral Ecology. 2018;29:1340–50.

54. Hinrichsen R, Holmes E. Using multivariate state-space models to study spatial structure and dynamics. Spatial Ecology. 2009;145–66.

55. Holmes E E, Ward E J, Wills K. MARSS: Multivariate Autoregressive State-space Models for Analyzing Time-series Data. The R Journal. 2012;4:11.

56. Ruckelshaus M, Currens K, Graeber W, Fuerstenberg R, Rawson K, Sands N, et al. Independent Populations of Chinook Salmon in Puget Sound. NOAA Technical Memorandum NMFS-NWFSC-78. 2006;

57. Austin CS, Torgersen CE, Quinn TP. Who spawns where? Temperature, elevation, and discharge differentially affect the distribution of breeding by six Pacific salmonids within a large river basin. Can J Fish Aquat Sci. 2023;80:1365–84.

58. Leach JA, Moore RD. Winter stream temperature in the rain-on-snow zone of the Pacific Northwest: influences of hillslope runoff and transient snow cover. Hydrol Earth Syst Sci. 2014;18:819–38.

59. Schreck CB, Stahl TP, Davis LE, Roby DD, Clemens BJ. Mortality estimates of juvenile spring–summer Chinook salmon in the lower Columbia River and estuary, 1992–1998: evidence for delayed mortality? Transactions of the American Fisheries Society. 2006;135:457–75.

60. Furey NB, Martins EG, Hinch SG. Migratory salmon smolts exhibit consistent interannual depensatory predator swamping: Effects on telemetry-based survival estimates. Ecology of Freshwater Fish. 2021;30:18–30.

61. Maynard DJ, Trial JG. The use of hatchery technology for the conservation of Pacific and Atlantic salmon. Rev Fish Biol Fisheries. 2014;24:803–17.

62. Bottom DL. To till the water—a history of ideas in fisheries conservation. In: Stouder DJ, Bisson PA, Naiman RJ, editors. Pacific Salmon & their Ecosystems: Status and Future Options [Internet]. New York: Chapman and Hall; 1997 [cited 2023 Nov 27]. p. 569–97. Available from: 10.1007/978-1-4615-6375-4_31

63. Anderson JH, Warheit KI, Craig BE, Seamons TR, Haukenes AH. A review of hatchery reform science in Washington State. Washington Department of Fish and Wildlife. 2020;

64. Johnson MS, Murdoch AR, Moran CPH. Adult survival of hatchery spring Chinook salmon released volitionally or forcibly as juveniles. North American Journal of Aquaculture. 2015;77:547–50.

65. Berdahl A, Westley PAH, Levin SA, Couzin ID, Quinn TP. A collective navigation hypothesis for homeward migration in anadromous salmonids. Fish and Fisheries. 2016;17:525–42.

66. Okasaki C, Keefer ML, Westley PAH, Berdahl A. Collective navigation can facilitate passage through human-made barriers by homeward migrating Pacific salmon. Proceedings of the Royal Society B: Biological Sciences. 2020;287:20202137.

67. Polyakov AY, Quinn TP, Myers KW, Berdahl A. Group size affects predation risk and foraging success in Pacific salmon at sea. Sci Adv. 2022;8:eabm7548.

68. Dibnah AJ, Herbert-Read JE, Boogert NJ, McIvor GE, Jolles JW, Thornton A. Vocally mediated consensus decisions govern mass departures from jackdaw roosts. Current Biology. 2022;32:R455–6.

69. Gall GEC, Strandburg-Peshkin A, Clutton-Brock T, Manser MB. As dusk falls: collective decisions about the return to sleeping sites in meerkats. Animal Behaviour. 2017;132:91–9.

70. Helm B, Piersma T, van der Jeugd H. Sociable schedules: interplay between avian seasonal and social behaviour. Animal Behaviour. 2006;72:245–62.

71. Sandlund OT, Diserud OH, Poole R, Bergesen K, Dillane M, Rogan G, et al. Timing and pattern of annual silver eel migration in two European watersheds are determined by similar cues. Ecology and Evolution. 2017;7:5956–66.

72. Berdahl A, Van Leeuwen A, Levin SA, Torney CJ. Collective behavior as a driver of critical transitions in migratory populations. Mov Ecol. 2016;4:18.

73. Fagan WF, Cantrell RS, Cosner C, Mueller T, Noble AE. Leadership, social learning, and the maintenance (or collapse) of migratory populations. Theor Ecol. 2012;5:253–64.

74. Granger J, Johnsen S. Collective movement as a solution to noisy navigation and its vulnerability to population loss. Proc R Soc B. 2022;289:20221910.

