## Supplementary Information for "Social influences complement environmental cues to stimulate migrating juvenile salmon"

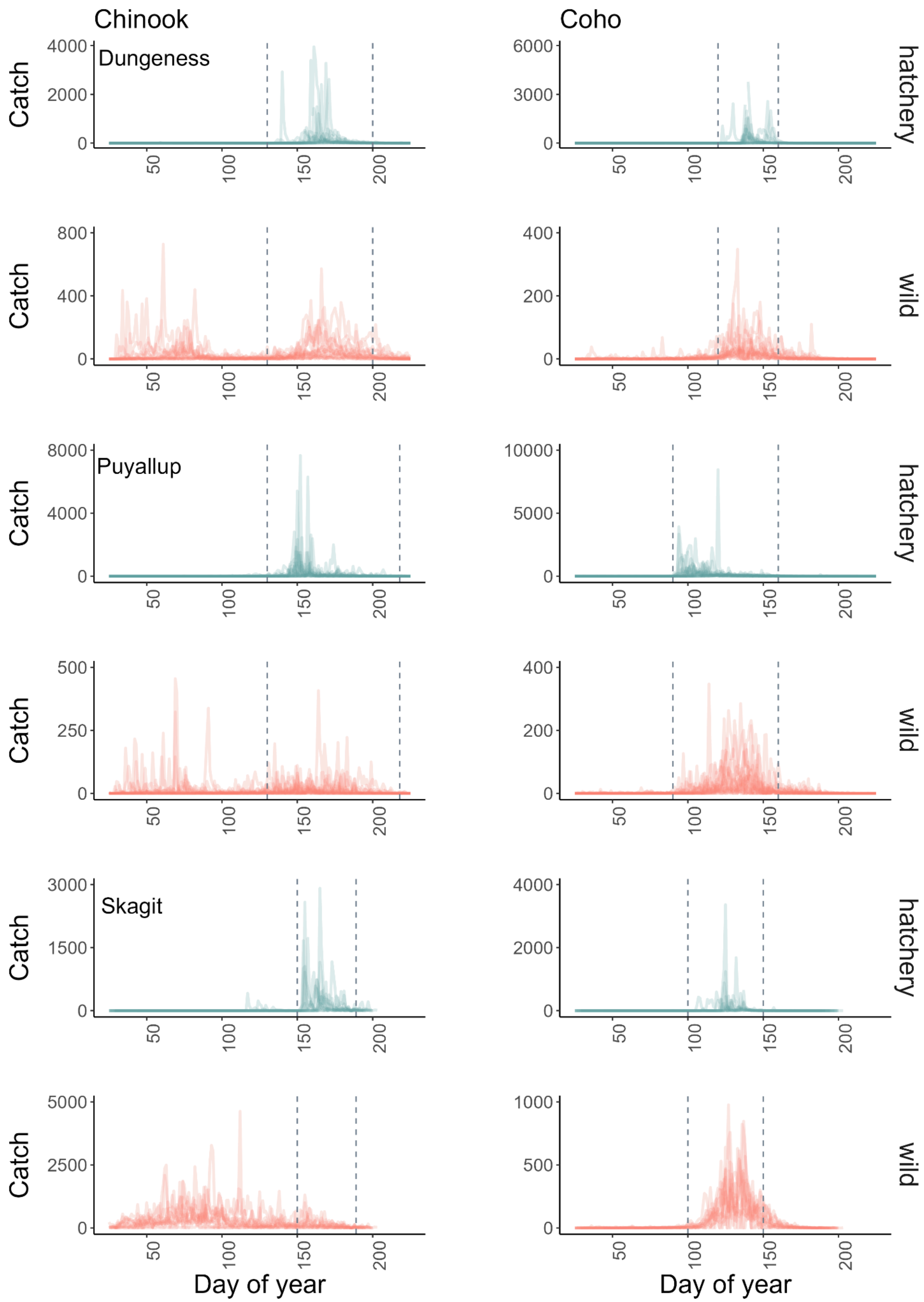

Chinook
Coho
Dungeness
Puyallup
Skagit
hatchery
wild
hatchery
wild
hatchery
wild
Catch
Day of year

Figure S1 - Daily catches for hatchery and wild sub-yearling Chinook salmon, hatchery and wild yearling coho salmon in the Dungeness (years 2005-2020), Puyallup (years 2004-2021), and Skagit rivers (years 2010-2022). Raw data without interpolation from every year available is overlaid. The vertical dashed lines bracket the period when both hatchery and wild salmon were caught in the trap, which was the portion of data we used for the analysis.

| River | Species | Origin | Total number caught | Max number caught |
|---|---|---|---|---|
| Dungeness | Chinook | hatchery | 62146 | 3274 |
| Dungeness | Chinook | wild | 42836 | 355 |
| Dungeness | Coho | hatchery | 58857 | 1983 |
| Dungeness | Coho | wild | 15414 | 348 |
| Puyallup | Chinook | hatchery | 153030 | 7429 |
| Puyallup | Chinook | wild | 15360 | 342 |
| Puyallup | Coho | hatchery | 155593 | 8436 |
| Puyallup | Coho | wild | 27759 | 291 |
| Skagit | Chinook | hatchery | 51332 | 2912 |
| Skagit | Chinook | wild | 47285 | 982 |
| Skagit | Coho | hatchery | 29207 | 3364 |
| Skagit | Coho | wild | 83347 | 976 |

Table S1 - Total and maximum number of fish caught in the trap during the days and years used in our analysis.

Unmarked hatchery salmon

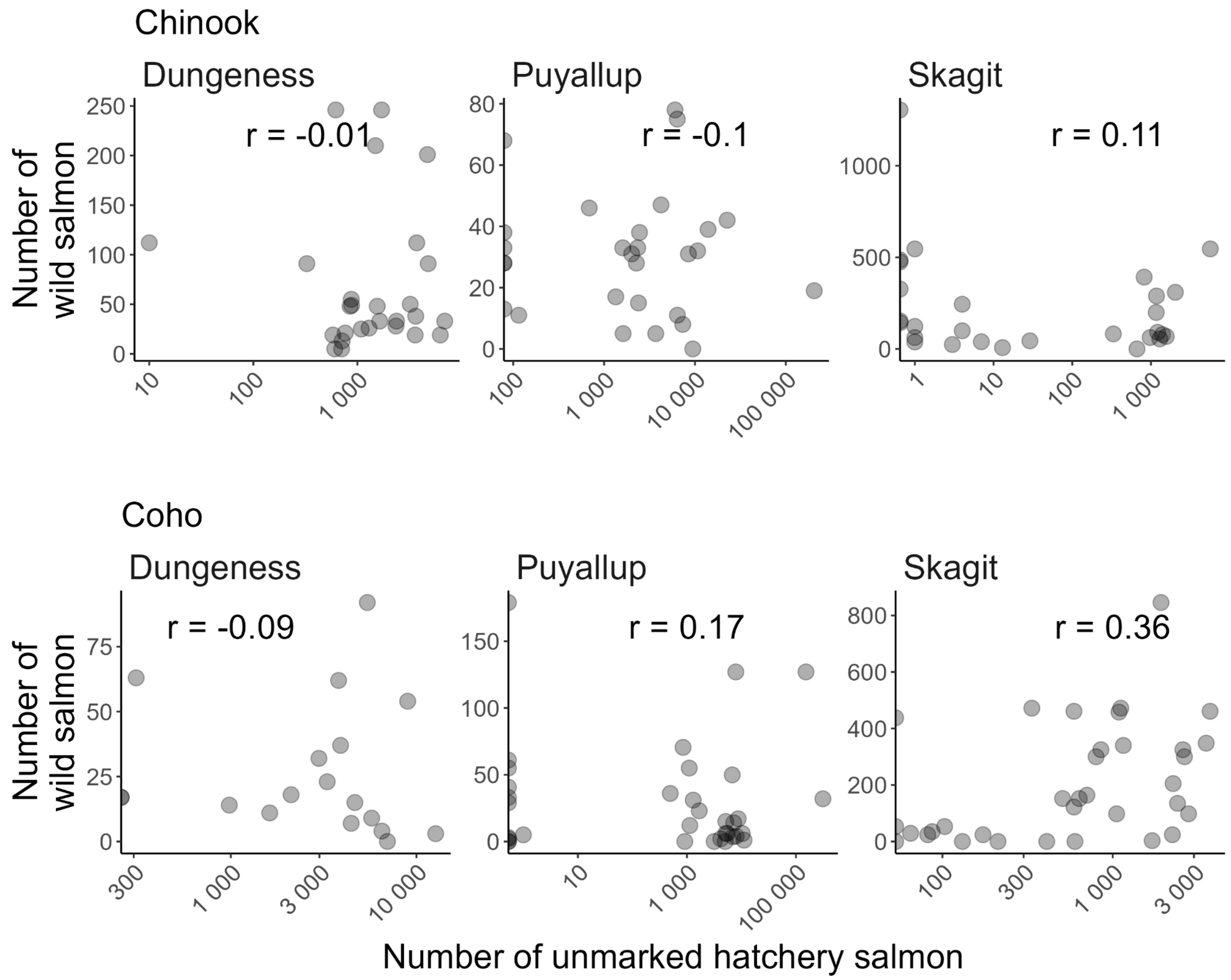


Figure S1 - Correlation between unmarked hatchery salmon and wild salmon for Chinook and coho salmon in the Dungeness, Puyallup, and Skagit rivers. Each data point represents a day with the maximum number of hatchery salmon caught in the trap within ten days of a hatchery release. Note log scale on the *x*-axis.

### Proportion of day vs. night migrants

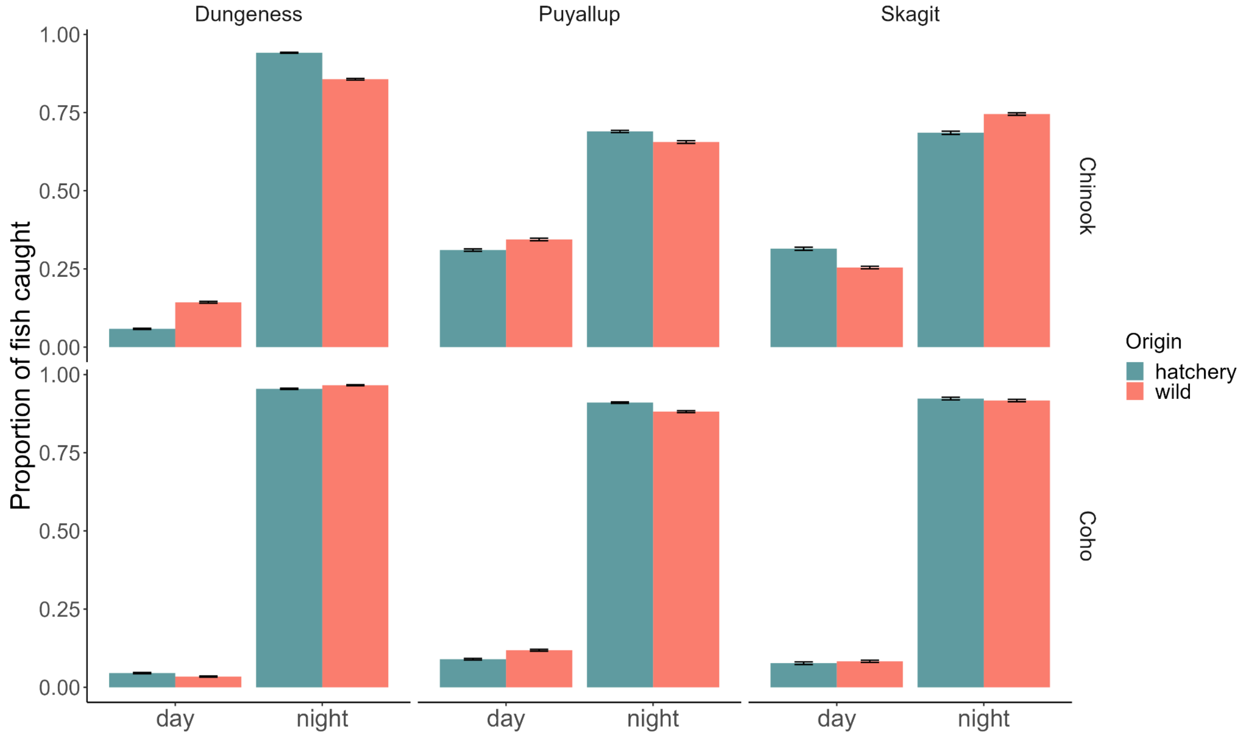


Figure S2 - Average proportion of Chinook salmon and coho salmon caught in the trap at night vs. day in all three rivers.

### Dungeness River

The Dungeness River is a 45-km river in the northwest part of Washington State. It originates in the Olympic Mountains and drains into the Strait of Juan de Fuca. The Dungeness River is fed through melting snowpack from the Olympic Mountains and by rain. The lowest flows are in August-September whereas the highest temperature is around 15° C in July-August. The Dungeness River supports seven species of salmonids - Chinook, coho, chum and pink salmon, steelhead, cutthroat and bull trout. There are multiple hatcheries that release Chinook and coho salmon and steelhead into the river at various stages. The Dungeness River hatchery, Gray Wolf, Hurd Creek, and Upper Dungeness Hatchery are the main hatcheries on the Dungeness River. A rotary screw trap is used to monitor juvenile salmon that migrate from their natal site to the ocean. The trap, operated by the Washington Department of Fish and Wildlife (WDFW), is located about 1.6 km upstream of the Strait of Juan De Fuca (48.1441, -123.1283).

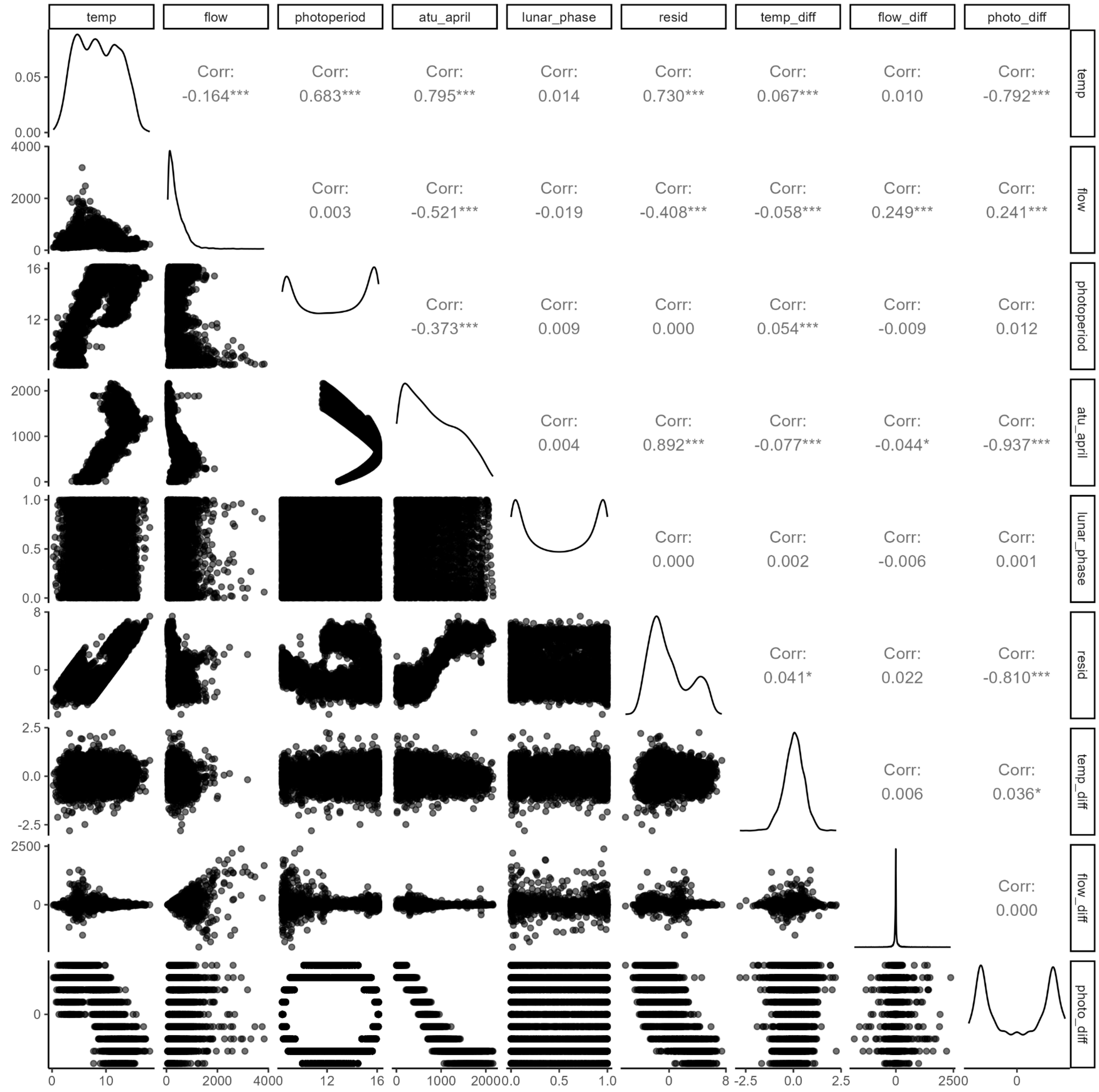


Figure S3 - Correlation between all the environmental covariates in the Dungeness River.

| **Chinook** | | **Coho** | |
|---|---|---|---|
| **Covariate** | $\Delta AICc$ | **Covariate** | $\Delta AICc$ |
| Photoperiod difference | 0 | Photoperiod | 0 |
| Flow | 3.73 | ATU | 3.03 |
| ATU | 5.50 | Flow | 27.9 |
| Temperature | 8.37 | Temperature | 30.01 |

| | | | |
|---|---|---|---|
| Residuals | 12.56 | Photoperiod difference | 43.12 |
| Photoperiod | 25.09 | Residuals | 44.16 |

Table S2 - Model selection for Chinook salmon and coho salmon in the Dungeness with each of the correlated covariates.

| Photoperiod | Photoperiod difference | Temperature difference | flow | Flow difference | Lunar phase | Hatchery | $\Delta AICc$ | rel.LL | weights | Cumulative weights |
|---|---|---|---|---|---|---|---|---|---|---|
| 0 | 1 | 1 | 0 | 1 | 0 | 1 | 0 | 1 | 0.24 | 0.24 |
| 0 | 1 | 1 | 0 | 1 | 1 | 1 | 0.42 | 0.81 | 0.2 | 0.44 |
| 1 | 1 | 1 | 0 | 1 | 0 | 1 | 0.82 | 0.66 | 0.16 | 0.6 |
| 1 | 1 | 1 | 0 | 1 | 1 | 1 | 1.21 | 0.55 | 0.13 | 0.73 |
| 0 | 1 | 1 | 1 | 1 | 0 | 1 | 2.09 | 0.35 | 0.09 | 0.82 |
| 0 | 1 | 1 | 1 | 1 | 1 | 1 | 2.51 | 0.28 | 0.07 | 0.89 |
| 1 | 1 | 1 | 1 | 1 | 0 | 1 | 2.9 | 0.23 | 0.06 | 0.94 |
| 1 | 1 | 1 | 1 | 1 | 1 | 1 | 3.29 | 0.19 | 0.05 | 0.99 |
| 1 | 0 | 1 | 1 | 1 | 0 | 1 | 9.98 | 0.01 | 0 | 0.99 |
| 1 | 0 | 1 | 1 | 1 | 1 | 1 | 10.27 | 0.01 | 0 | 0.99 |
| 1 | 0 | 1 | 0 | 1 | 0 | 1 | 10.72 | 0 | 0 | 1 |
| 0 | 1 | 1 | 0 | 1 | 0 | 0 | 11.06 | 0 | 0 | 1 |
| 1 | 0 | 1 | 0 | 1 | 1 | 1 | 11.09 | 0 | 0 | 1 |
| 1 | 1 | 1 | 0 | 1 | 0 | 0 | 12.15 | 0 | 0 | 1 |
| 0 | 1 | 1 | 0 | 1 | 1 | 0 | 12.44 | 0 | 0 | 1 |
| 0 | 1 | 1 | 1 | 1 | 0 | 0 | 12.81 | 0 | 0 | 1 |
| 1 | 1 | 1 | 0 | 1 | 1 | 0 | 13.44 | 0 | 0 | 1 |

| Photoperiod | Photoperiod difference | Temperature difference | flow | Flow difference | Lunar phase | Hatchery | $\Delta AICc$ | rel.LL | weights | Cumulative weights |
|---|---|---|---|---|---|---|---|---|---|---|
| 1 | 1 | 1 | 1 | 1 | 0 | 0 | 14.08 | 0 | 0 | 1 |
| 0 | 1 | 1 | 1 | 1 | 1 | 0 | 14.15 | 0 | 0 | 1 |
| 1 | 1 | 1 | 1 | 1 | 1 | 0 | 15.36 | 0 | 0 | 1 |
| 0 | 0 | 1 | 1 | 1 | 0 | 1 | 18.57 | 0 | 0 | 1 |
| 0 | 0 | 1 | 1 | 1 | 1 | 1 | 18.91 | 0 | 0 | 1 |
| 0 | 0 | 1 | 0 | 1 | 0 | 1 | 21.12 | 0 | 0 | 1 |
| 0 | 0 | 1 | 0 | 1 | 1 | 1 | 21.56 | 0 | 0 | 1 |
| 1 | 1 | 1 | 0 | 0 | 0 | 1 | 28.41 | 0 | 0 | 1 |
| 0 | 1 | 1 | 0 | 0 | 0 | 1 | 28.46 | 0 | 0 | 1 |
| 1 | 1 | 1 | 0 | 0 | 1 | 1 | 29.34 | 0 | 0 | 1 |
| 0 | 1 | 1 | 0 | 0 | 1 | 1 | 29.43 | 0 | 0 | 1 |
| 1 | 1 | 1 | 1 | 0 | 0 | 1 | 29.62 | 0 | 0 | 1 |
| 0 | 1 | 1 | 1 | 0 | 0 | 1 | 30 | 0 | 0 | 1 |
| 0 | 0 | 1 | 1 | 1 | 0 | 0 | 30.21 | 0 | 0 | 1 |
| 1 | 0 | 1 | 1 | 1 | 0 | 0 | 30.47 | 0 | 0 | 1 |
| 1 | 1 | 1 | 1 | 0 | 1 | 1 | 30.5 | 0 | 0 | 1 |
| 0 | 1 | 1 | 1 | 0 | 1 | 1 | 30.95 | 0 | 0 | 1 |
| 0 | 0 | 1 | 1 | 1 | 1 | 0 | 31.51 | 0 | 0 | 1 |

| Photoperiod | Photoperiod difference | Temperature difference | flow | Flow difference | Lunar phase | Hatchery | $\Delta AICc$ | rel.LL | weights | Cumulative weights |
|---|---|---|---|---|---|---|---|---|---|---|
| 1 | 0 | 1 | 1 | 1 | 1 | 0 | 31.87 | 0 | 0 | 1 |
| 1 | 0 | 1 | 1 | 0 | 0 | 1 | 36.38 | 0 | 0 | 1 |
| 0 | 1 | 1 | 0 | 0 | 0 | 0 | 36.56 | 0 | 0 | 1 |
| 1 | 0 | 1 | 0 | 1 | 0 | 0 | 36.78 | 0 | 0 | 1 |
| 0 | 0 | 1 | 0 | 1 | 0 | 0 | 37.01 | 0 | 0 | 1 |
| 0 | 1 | 1 | 1 | 0 | 0 | 0 | 37.1 | 0 | 0 | 1 |
| 1 | 0 | 1 | 1 | 0 | 1 | 1 | 37.16 | 0 | 0 | 1 |
| 0 | 1 | 1 | 0 | 0 | 1 | 0 | 38.19 | 0 | 0 | 1 |
| 1 | 0 | 1 | 0 | 1 | 1 | 0 | 38.36 | 0 | 0 | 1 |
| 1 | 1 | 1 | 0 | 0 | 0 | 0 | 38.41 | 0 | 0 | 1 |
| 0 | 0 | 1 | 0 | 1 | 1 | 0 | 38.5 | 0 | 0 | 1 |
| 0 | 1 | 1 | 1 | 0 | 1 | 0 | 38.67 | 0 | 0 | 1 |
| 1 | 1 | 1 | 1 | 0 | 0 | 0 | 39.12 | 0 | 0 | 1 |
| 1 | 1 | 1 | 0 | 0 | 1 | 0 | 40.01 | 0 | 0 | 1 |
| 1 | 1 | 1 | 1 | 0 | 1 | 0 | 40.67 | 0 | 0 | 1 |
| 1 | 0 | 1 | 0 | 0 | 0 | 1 | 41.06 | 0 | 0 | 1 |
| 1 | 0 | 1 | 0 | 0 | 1 | 1 | 41.95 | 0 | 0 | 1 |
| 0 | 0 | 1 | 1 | 0 | 0 | 1 | 48.28 | 0 | 0 | 1 |

| Photoperiod | Photoperiod difference | Temperature difference | flow | Flow difference | Lunar phase | Hatchery | $\Delta AICc$ | rel.LL | weights | Cumulative weights |
|---|---|---|---|---|---|---|---|---|---|---|
| 0 | 0 | 1 | 1 | 0 | 1 | 1 | 49.14 | 0 | 0 | 1 |
| 1 | 0 | 1 | 1 | 0 | 0 | 0 | 54.45 | 0 | 0 | 1 |
| 1 | 0 | 1 | 1 | 0 | 1 | 0 | 56.08 | 0 | 0 | 1 |
| 0 | 0 | 1 | 0 | 0 | 0 | 1 | 56.22 | 0 | 0 | 1 |
| 0 | 0 | 1 | 1 | 0 | 0 | 0 | 56.3 | 0 | 0 | 1 |
| 0 | 0 | 1 | 0 | 0 | 1 | 1 | 57.22 | 0 | 0 | 1 |
| 0 | 0 | 1 | 1 | 0 | 1 | 0 | 57.83 | 0 | 0 | 1 |
| 1 | 0 | 1 | 0 | 0 | 0 | 0 | 66.16 | 0 | 0 | 1 |
| 1 | 0 | 1 | 0 | 0 | 1 | 0 | 67.98 | 0 | 0 | 1 |
| 0 | 0 | 1 | 0 | 0 | 0 | 0 | 69.13 | 0 | 0 | 1 |
| 0 | 0 | 1 | 0 | 0 | 1 | 0 | 70.86 | 0 | 0 | 1 |
| 1 | 1 | 0 | 1 | 1 | 0 | 1 | 84.74 | 0 | 0 | 1 |
| 1 | 1 | 0 | 1 | 1 | 1 | 1 | 84.84 | 0 | 0 | 1 |
| 1 | 0 | 0 | 1 | 1 | 0 | 1 | 85.38 | 0 | 0 | 1 |
| 1 | 0 | 0 | 1 | 1 | 1 | 1 | 85.44 | 0 | 0 | 1 |
| 0 | 1 | 0 | 1 | 1 | 0 | 1 | 86.63 | 0 | 0 | 1 |
| 0 | 1 | 0 | 1 | 1 | 1 | 1 | 86.81 | 0 | 0 | 1 |
| 1 | 1 | 0 | 0 | 1 | 0 | 1 | 88.31 | 0 | 0 | 1 |

| Photoperiod | Photoperiod difference | Temperature difference | flow | Flow difference | Lunar phase | Hatchery | $\Delta AICc$ | rel.LL | weights | Cumulative weights |
|---|---|---|---|---|---|---|---|---|---|---|
| 1 | 1 | 0 | 0 | 1 | 1 | 1 | 88.51 | 0 | 0 | 1 |
| 0 | 1 | 0 | 0 | 1 | 0 | 1 | 89.17 | 0 | 0 | 1 |
| 0 | 1 | 0 | 0 | 1 | 1 | 1 | 89.43 | 0 | 0 | 1 |
| 0 | 1 | 0 | 1 | 1 | 0 | 0 | 89.87 | 0 | 0 | 1 |
| 0 | 1 | 0 | 1 | 1 | 1 | 0 | 90.77 | 0 | 0 | 1 |
| 1 | 1 | 0 | 1 | 1 | 0 | 0 | 91.8 | 0 | 0 | 1 |
| 1 | 1 | 0 | 1 | 1 | 1 | 0 | 92.74 | 0 | 0 | 1 |
| 0 | 1 | 0 | 0 | 1 | 0 | 0 | 94.37 | 0 | 0 | 1 |
| 0 | 0 | 0 | 1 | 1 | 0 | 1 | 94.75 | 0 | 0 | 1 |
| 0 | 0 | 0 | 1 | 1 | 1 | 1 | 94.85 | 0 | 0 | 1 |
| 1 | 0 | 0 | 0 | 1 | 0 | 1 | 95.25 | 0 | 0 | 1 |
| 0 | 1 | 0 | 0 | 1 | 1 | 0 | 95.42 | 0 | 0 | 1 |
| 1 | 0 | 0 | 0 | 1 | 1 | 1 | 95.42 | 0 | 0 | 1 |
| 1 | 1 | 0 | 0 | 1 | 0 | 0 | 96.45 | 0 | 0 | 1 |
| 1 | 1 | 0 | 0 | 1 | 1 | 0 | 97.49 | 0 | 0 | 1 |
| 1 | 0 | 0 | 1 | 1 | 0 | 0 | 97.63 | 0 | 0 | 1 |
| 1 | 0 | 0 | 1 | 1 | 1 | 0 | 98.69 | 0 | 0 | 1 |
| 0 | 0 | 0 | 1 | 1 | 0 | 0 | 99.15 | 0 | 0 | 1 |

| Photoperiod | Photoperiod difference | Temperature difference | flow | Flow difference | Lunar phase | Hatchery | $\Delta AICc$ | rel.LL | weights | Cumulative weights |
|---|---|---|---|---|---|---|---|---|---|---|
| 0 | 0 | 0 | 1 | 1 | 1 | 0 | 100.04 | 0 | 0 | 1 |
| 1 | 1 | 0 | 1 | 0 | 0 | 1 | 102.95 | 0 | 0 | 1 |
| 1 | 1 | 0 | 1 | 0 | 1 | 1 | 103.51 | 0 | 0 | 1 |
| 1 | 0 | 0 | 1 | 0 | 0 | 1 | 103.92 | 0 | 0 | 1 |
| 1 | 0 | 0 | 1 | 0 | 1 | 1 | 104.42 | 0 | 0 | 1 |
| 0 | 1 | 0 | 1 | 0 | 0 | 1 | 106.24 | 0 | 0 | 1 |
| 0 | 1 | 0 | 1 | 0 | 1 | 1 | 106.9 | 0 | 0 | 1 |
| 0 | 1 | 0 | 1 | 0 | 0 | 0 | 107.79 | 0 | 0 | 1 |
| 0 | 0 | 0 | 0 | 1 | 0 | 1 | 108.41 | 0 | 0 | 1 |
| 0 | 0 | 0 | 0 | 1 | 1 | 1 | 108.67 | 0 | 0 | 1 |
| 0 | 1 | 0 | 1 | 0 | 1 | 0 | 108.97 | 0 | 0 | 1 |
| 1 | 1 | 0 | 1 | 0 | 0 | 0 | 109.1 | 0 | 0 | 1 |
| 1 | 1 | 0 | 0 | 0 | 0 | 1 | 109.97 | 0 | 0 | 1 |
| 1 | 1 | 0 | 1 | 0 | 1 | 0 | 110.35 | 0 | 0 | 1 |
| 1 | 1 | 0 | 0 | 0 | 1 | 1 | 110.67 | 0 | 0 | 1 |
| 0 | 1 | 0 | 0 | 0 | 0 | 1 | 111.78 | 0 | 0 | 1 |
| 0 | 1 | 0 | 0 | 0 | 1 | 1 | 112.56 | 0 | 0 | 1 |
| 1 | 0 | 0 | 1 | 0 | 0 | 0 | 115.15 | 0 | 0 | 1 |

| Photoperiod | Photoperiod difference | Temperature difference | flow | Flow difference | Lunar phase | Hatchery | $\Delta AICc$ | rel.LL | weights | Cumulative weights |
|---|---|---|---|---|---|---|---|---|---|---|
| 0 | 1 | 0 | 0 | 0 | 0 | 0 | 115.34 | 0 | 0 | 1 |
| 1 | 0 | 0 | 0 | 1 | 0 | 0 | 115.67 | 0 | 0 | 1 |
| 0 | 0 | 0 | 1 | 0 | 0 | 1 | 116.42 | 0 | 0 | 1 |
| 1 | 0 | 0 | 1 | 0 | 1 | 0 | 116.48 | 0 | 0 | 1 |
| 0 | 1 | 0 | 0 | 0 | 1 | 0 | 116.68 | 0 | 0 | 1 |
| 0 | 0 | 0 | 1 | 0 | 1 | 1 | 117.01 | 0 | 0 | 1 |
| 1 | 0 | 0 | 0 | 1 | 1 | 0 | 117.03 | 0 | 0 | 1 |
| 1 | 1 | 0 | 0 | 0 | 0 | 0 | 117.28 | 0 | 0 | 1 |
| 0 | 0 | 0 | 0 | 1 | 0 | 0 | 118.35 | 0 | 0 | 1 |
| 1 | 1 | 0 | 0 | 0 | 1 | 0 | 118.65 | 0 | 0 | 1 |
| 0 | 0 | 0 | 1 | 0 | 0 | 0 | 119.06 | 0 | 0 | 1 |
| 0 | 0 | 0 | 0 | 1 | 1 | 0 | 119.57 | 0 | 0 | 1 |
| 1 | 0 | 0 | 0 | 0 | 0 | 1 | 119.96 | 0 | 0 | 1 |
| 0 | 0 | 0 | 1 | 0 | 1 | 0 | 120.22 | 0 | 0 | 1 |
| 1 | 0 | 0 | 0 | 0 | 1 | 1 | 120.63 | 0 | 0 | 1 |
| 0 | 0 | 0 | 0 | 0 | 0 | 1 | 138.12 | 0 | 0 | 1 |
| 0 | 0 | 0 | 0 | 0 | 1 | 1 | 138.93 | 0 | 0 | 1 |
| 1 | 0 | 0 | 0 | 0 | 0 | 0 | 140.43 | 0 | 0 | 1 |

| Photoperiod | Photoperiod difference | Temperature difference | flow | Flow difference | Lunar phase | Hatchery | $\Delta AICc$ | rel.LL | weights | Cumulative weights |
|---|---|---|---|---|---|---|---|---|---|---|
| 1 | 0 | 0 | 0 | 0 | 1 | 0 | 142.07 | 0 | 0 | 1 |
| 0 | 0 | 0 | 0 | 0 | 0 | 0 | 146.49 | 0 | 0 | 1 |
| 0 | 0 | 0 | 0 | 0 | 1 | 0 | 147.99 | 0 | 0 | 1 |

Table S3 - Model selection with all combinations of uncorrelated covariates for Chinook yearlings in the Dungeness River.

| **Chinook** | | **Coho** | |
|---|---|---|---|
| **Variable** | **Relative Importance** | **Variable** | **Relative Importance** |
| Temperature difference | 1 | Photoperiod | 1 |
| Flow difference | 1 | Flow difference | 1 |
| Photoperiod difference | 1 | Flow | 0.9 |
| Hatchery | 1 | Temperature difference | 0.5 |
| Lunar Phase | 0.4 | Hatchery | 0.4 |
| Photoperiod | 0.4 | Photoperiod difference | 0.3 |
| Flow | 0.3 | Lunar phase | 0.3 |

Table S4 - Relative variable importance for all variables used in the model selection process for sub-yearling Chinook salmon and yearling coho salmon in the Dungeness River.

| Photoperiod | Photo difference | Temperature difference | Flow | Flow difference | Lunar phase | Hatchery | $\Delta AICc$ | rel.LL | weights | Cumulative weights |
|---|---|---|---|---|---|---|---|---|---|---|
| 1 | 0 | 1 | 1 | 1 | 0 | 0 | 0 | 1 | 0.17 | 0.17 |
| 1 | 0 | 0 | 1 | 1 | 0 | 0 | 0.52 | 0.77 | 0.13 | 0.31 |
| 1 | 0 | 1 | 1 | 1 | 0 | 1 | 1.01 | 0.6 | 0.11 | 0.41 |
| 1 | 0 | 0 | 1 | 1 | 0 | 1 | 1.16 | 0.56 | 0.1 | 0.51 |
| 1 | 0 | 1 | 1 | 1 | 1 | 0 | 2 | 0.37 | 0.06 | 0.58 |
| 1 | 1 | 1 | 1 | 1 | 0 | 0 | 2.14 | 0.34 | 0.06 | 0.64 |
| 1 | 1 | 0 | 1 | 1 | 0 | 0 | 2.5 | 0.29 | 0.05 | 0.69 |
| 1 | 0 | 0 | 1 | 1 | 1 | 0 | 2.53 | 0.28 | 0.05 | 0.74 |
| 1 | 0 | 1 | 1 | 1 | 1 | 1 | 3.02 | 0.22 | 0.04 | 0.77 |
| 1 | 1 | 1 | 1 | 1 | 0 | 1 | 3.15 | 0.21 | 0.04 | 0.81 |
| 1 | 0 | 0 | 1 | 1 | 1 | 1 | 3.17 | 0.21 | 0.04 | 0.85 |
| 1 | 1 | 0 | 1 | 1 | 0 | 1 | 3.27 | 0.19 | 0.03 | 0.88 |
| 1 | 1 | 1 | 1 | 1 | 1 | 0 | 4.14 | 0.13 | 0.02 | 0.9 |
| 1 | 1 | 0 | 1 | 1 | 1 | 0 | 4.53 | 0.1 | 0.02 | 0.92 |
| 1 | 1 | 1 | 1 | 1 | 1 | 1 | 5.14 | 0.08 | 0.01 | 0.93 |
| 1 | 1 | 0 | 1 | 1 | 1 | 1 | 5.3 | 0.07 | 0.01 | 0.95 |
| 1 | 0 | 0 | 0 | 1 | 0 | 1 | 5.68 | 0.06 | 0.01 | 0.96 |
| 1 | 0 | 0 | 0 | 1 | 0 | 0 | 5.99 | 0.05 | 0.01 | 0.97 |

| Photoperiod | Photo difference | Temperature difference | Flow | Flow difference | Lunar phase | Hatchery | $\Delta AICc$ | rel.LL | weights | Cumulative weights |
|---|---|---|---|---|---|---|---|---|---|---|
| 1 | 0 | 1 | 0 | 1 | 0 | 1 | 7.47 | 0.02 | 0 | 0.97 |
| 1 | 0 | 1 | 0 | 1 | 0 | 0 | 7.51 | 0.02 | 0 | 0.97 |
| 1 | 0 | 0 | 0 | 1 | 1 | 1 | 7.76 | 0.02 | 0 | 0.98 |
| 1 | 1 | 0 | 0 | 1 | 0 | 0 | 7.79 | 0.02 | 0 | 0.98 |
| 1 | 1 | 0 | 0 | 1 | 0 | 1 | 7.82 | 0.02 | 0 | 0.98 |
| 1 | 0 | 0 | 0 | 1 | 1 | 0 | 8.08 | 0.02 | 0 | 0.99 |
| 1 | 1 | 1 | 0 | 1 | 0 | 0 | 9.51 | 0.01 | 0 | 0.99 |
| 1 | 1 | 1 | 0 | 1 | 0 | 1 | 9.55 | 0.01 | 0 | 0.99 |
| 1 | 0 | 1 | 0 | 1 | 1 | 1 | 9.56 | 0.01 | 0 | 0.99 |
| 1 | 0 | 1 | 0 | 1 | 1 | 0 | 9.6 | 0.01 | 0 | 0.99 |
| 1 | 1 | 0 | 0 | 1 | 1 | 0 | 9.89 | 0.01 | 0 | 0.99 |
| 1 | 1 | 0 | 0 | 1 | 1 | 1 | 9.91 | 0.01 | 0 | 1 |
| 1 | 1 | 1 | 0 | 1 | 1 | 0 | 11.61 | 0 | 0 | 1 |
| 1 | 1 | 1 | 0 | 1 | 1 | 1 | 11.64 | 0 | 0 | 1 |
| 1 | 0 | 0 | 1 | 0 | 0 | 0 | 12.18 | 0 | 0 | 1 |
| 1 | 0 | 0 | 1 | 0 | 0 | 1 | 12.66 | 0 | 0 | 1 |
| 1 | 0 | 1 | 1 | 0 | 0 | 0 | 13.3 | 0 | 0 | 1 |
| 1 | 0 | 0 | 0 | 0 | 0 | 0 | 13.85 | 0 | 0 | 1 |

| Photoperiod | Photo difference | Temperature difference | Flow | Flow difference | Lunar phase | Hatchery | $\Delta AICc$ | rel.LL | weights | Cumulative weights |
|---|---|---|---|---|---|---|---|---|---|---|
| 1 | 0 | 0 | 1 | 0 | 1 | 0 | 13.88 | 0 | 0 | 1 |
| 1 | 0 | 1 | 1 | 0 | 0 | 1 | 13.9 | 0 | 0 | 1 |
| 1 | 0 | 0 | 0 | 0 | 0 | 1 | 14.03 | 0 | 0 | 1 |
| 1 | 1 | 0 | 1 | 0 | 0 | 0 | 14.05 | 0 | 0 | 1 |
| 1 | 1 | 0 | 1 | 0 | 0 | 1 | 14.25 | 0 | 0 | 1 |
| 1 | 0 | 0 | 1 | 0 | 1 | 1 | 14.36 | 0 | 0 | 1 |
| 1 | 1 | 1 | 1 | 0 | 0 | 0 | 14.59 | 0 | 0 | 1 |
| 1 | 1 | 1 | 1 | 0 | 0 | 1 | 14.98 | 0 | 0 | 1 |
| 1 | 0 | 1 | 1 | 0 | 1 | 0 | 14.99 | 0 | 0 | 1 |
| 1 | 1 | 0 | 0 | 0 | 0 | 1 | 15.42 | 0 | 0 | 1 |
| 1 | 0 | 1 | 1 | 0 | 1 | 1 | 15.59 | 0 | 0 | 1 |
| 1 | 0 | 0 | 0 | 0 | 1 | 0 | 15.7 | 0 | 0 | 1 |
| 1 | 1 | 0 | 1 | 0 | 1 | 0 | 15.73 | 0 | 0 | 1 |
| 1 | 0 | 1 | 0 | 0 | 0 | 0 | 15.73 | 0 | 0 | 1 |
| 1 | 0 | 0 | 0 | 0 | 1 | 1 | 15.88 | 0 | 0 | 1 |
| 1 | 1 | 0 | 1 | 0 | 1 | 1 | 15.9 | 0 | 0 | 1 |
| 1 | 1 | 0 | 0 | 0 | 0 | 0 | 15.91 | 0 | 0 | 1 |
| 1 | 0 | 1 | 0 | 0 | 0 | 1 | 16.06 | 0 | 0 | 1 |

| Photoperiod | Photo difference | Temperature difference | Flow | Flow difference | Lunar phase | Hatchery | $\Delta AICc$ | rel.LL | weights | Cumulative weights |
|---|---|---|---|---|---|---|---|---|---|---|
| 1 | 1 | 1 | 1 | 0 | 1 | 0 | 16.21 | 0 | 0 | 1 |
| 1 | 1 | 1 | 1 | 0 | 1 | 1 | 16.58 | 0 | 0 | 1 |
| 1 | 1 | 1 | 0 | 0 | 0 | 1 | 17.21 | 0 | 0 | 1 |
| 1 | 1 | 0 | 0 | 0 | 1 | 1 | 17.21 | 0 | 0 | 1 |
| 1 | 0 | 1 | 0 | 0 | 1 | 0 | 17.59 | 0 | 0 | 1 |
| 1 | 1 | 1 | 0 | 0 | 0 | 0 | 17.69 | 0 | 0 | 1 |
| 1 | 1 | 0 | 0 | 0 | 1 | 0 | 17.76 | 0 | 0 | 1 |
| 1 | 0 | 1 | 0 | 0 | 1 | 1 | 17.91 | 0 | 0 | 1 |
| 0 | 0 | 1 | 1 | 1 | 0 | 0 | 18.28 | 0 | 0 | 1 |
| 1 | 1 | 1 | 0 | 0 | 1 | 1 | 18.99 | 0 | 0 | 1 |
| 1 | 1 | 1 | 0 | 0 | 1 | 0 | 19.53 | 0 | 0 | 1 |
| 0 | 0 | 1 | 1 | 1 | 0 | 1 | 19.76 | 0 | 0 | 1 |
| 0 | 1 | 1 | 1 | 1 | 0 | 0 | 20.05 | 0 | 0 | 1 |
| 0 | 0 | 0 | 1 | 1 | 0 | 0 | 20.36 | 0 | 0 | 1 |
| 0 | 0 | 1 | 1 | 1 | 1 | 0 | 20.39 | 0 | 0 | 1 |
| 0 | 1 | 1 | 1 | 1 | 0 | 1 | 21.07 | 0 | 0 | 1 |
| 0 | 0 | 0 | 1 | 1 | 0 | 1 | 21.37 | 0 | 0 | 1 |
| 0 | 0 | 1 | 1 | 1 | 1 | 1 | 21.88 | 0 | 0 | 1 |

| Photoperiod | Photo difference | Temperature difference | Flow | Flow difference | Lunar phase | Hatchery | $\Delta AICc$ | rel.LL | weights | Cumulative weights |
|---|---|---|---|---|---|---|---|---|---|---|
| 0 | 1 | 1 | 1 | 1 | 1 | 0 | 22.16 | 0 | 0 | 1 |
| 0 | 0 | 0 | 1 | 1 | 1 | 0 | 22.48 | 0 | 0 | 1 |
| 0 | 1 | 0 | 1 | 1 | 0 | 0 | 22.5 | 0 | 0 | 1 |
| 0 | 1 | 1 | 1 | 1 | 1 | 1 | 23.17 | 0 | 0 | 1 |
| 0 | 1 | 0 | 1 | 1 | 0 | 1 | 23.28 | 0 | 0 | 1 |
| 0 | 0 | 0 | 1 | 1 | 1 | 1 | 23.5 | 0 | 0 | 1 |
| 0 | 1 | 0 | 1 | 1 | 1 | 0 | 24.63 | 0 | 0 | 1 |
| 0 | 1 | 0 | 1 | 1 | 1 | 1 | 25.4 | 0 | 0 | 1 |
| 0 | 0 | 0 | 0 | 1 | 0 | 1 | 39.86 | 0 | 0 | 1 |
| 0 | 1 | 1 | 1 | 0 | 0 | 1 | 40.2 | 0 | 0 | 1 |
| 0 | 1 | 0 | 0 | 1 | 0 | 1 | 40.26 | 0 | 0 | 1 |
| 0 | 1 | 1 | 1 | 0 | 0 | 0 | 40.79 | 0 | 0 | 1 |
| 0 | 0 | 0 | 0 | 1 | 0 | 0 | 41.33 | 0 | 0 | 1 |
| 0 | 0 | 1 | 1 | 0 | 0 | 0 | 41.44 | 0 | 0 | 1 |
| 0 | 1 | 0 | 1 | 0 | 0 | 1 | 41.57 | 0 | 0 | 1 |
| 0 | 0 | 0 | 1 | 0 | 0 | 0 | 41.75 | 0 | 0 | 1 |
| 0 | 0 | 1 | 0 | 1 | 0 | 1 | 41.83 | 0 | 0 | 1 |
| 0 | 1 | 1 | 0 | 1 | 0 | 1 | 41.83 | 0 | 0 | 1 |

| Photoperiod | Photo difference | Temperature difference | Flow | Flow difference | Lunar phase | Hatchery | $\Delta AICc$ | rel.LL | weights | Cumulative weights |
|---|---|---|---|---|---|---|---|---|---|---|
| 0 | 1 | 1 | 1 | 0 | 1 | 1 | 41.96 | 0 | 0 | 1 |
| 0 | 0 | 0 | 0 | 1 | 1 | 1 | 42 | 0 | 0 | 1 |
| 0 | 1 | 0 | 0 | 1 | 1 | 1 | 42.42 | 0 | 0 | 1 |
| 0 | 1 | 0 | 1 | 0 | 0 | 0 | 42.46 | 0 | 0 | 1 |
| 0 | 0 | 1 | 1 | 0 | 0 | 1 | 42.53 | 0 | 0 | 1 |
| 0 | 0 | 0 | 1 | 0 | 0 | 1 | 42.55 | 0 | 0 | 1 |
| 0 | 1 | 1 | 1 | 0 | 1 | 0 | 42.61 | 0 | 0 | 1 |
| 0 | 0 | 1 | 0 | 1 | 0 | 0 | 43.13 | 0 | 0 | 1 |
| 0 | 0 | 1 | 1 | 0 | 1 | 0 | 43.33 | 0 | 0 | 1 |
| 0 | 1 | 0 | 1 | 0 | 1 | 1 | 43.4 | 0 | 0 | 1 |
| 0 | 0 | 0 | 0 | 1 | 1 | 0 | 43.46 | 0 | 0 | 1 |
| 0 | 1 | 0 | 0 | 1 | 0 | 0 | 43.46 | 0 | 0 | 1 |
| 0 | 0 | 0 | 1 | 0 | 1 | 0 | 43.67 | 0 | 0 | 1 |
| 0 | 0 | 1 | 0 | 1 | 1 | 1 | 43.98 | 0 | 0 | 1 |
| 0 | 1 | 1 | 0 | 1 | 1 | 1 | 43.99 | 0 | 0 | 1 |
| 0 | 1 | 0 | 1 | 0 | 1 | 0 | 44.35 | 0 | 0 | 1 |
| 0 | 0 | 1 | 1 | 0 | 1 | 1 | 44.43 | 0 | 0 | 1 |
| 0 | 0 | 0 | 1 | 0 | 1 | 1 | 44.47 | 0 | 0 | 1 |

| Photoperiod | Photo difference | Temperature difference | Flow | Flow difference | Lunar phase | Hatchery | $\Delta AICc$ | rel.LL | weights | Cumulative weights |
|---|---|---|---|---|---|---|---|---|---|---|
| 0 | 1 | 1 | 0 | 1 | 0 | 0 | 45.22 | 0 | 0 | 1 |
| 0 | 0 | 1 | 0 | 1 | 1 | 0 | 45.27 | 0 | 0 | 1 |
| 0 | 1 | 0 | 0 | 1 | 1 | 0 | 45.6 | 0 | 0 | 1 |
| 0 | 1 | 1 | 0 | 1 | 1 | 0 | 47.36 | 0 | 0 | 1 |
| 0 | 1 | 0 | 0 | 0 | 0 | 1 | 52.19 | 0 | 0 | 1 |
| 0 | 1 | 1 | 0 | 0 | 0 | 1 | 53.68 | 0 | 0 | 1 |
| 0 | 1 | 0 | 0 | 0 | 1 | 1 | 54.21 | 0 | 0 | 1 |
| 0 | 0 | 0 | 0 | 0 | 0 | 1 | 55.07 | 0 | 0 | 1 |
| 0 | 1 | 1 | 0 | 0 | 1 | 1 | 55.69 | 0 | 0 | 1 |
| 0 | 0 | 0 | 0 | 0 | 0 | 0 | 55.91 | 0 | 0 | 1 |
| 0 | 1 | 0 | 0 | 0 | 0 | 0 | 56.96 | 0 | 0 | 1 |
| 0 | 0 | 1 | 0 | 0 | 0 | 1 | 57.14 | 0 | 0 | 1 |
| 0 | 0 | 0 | 0 | 0 | 1 | 1 | 57.16 | 0 | 0 | 1 |
| 0 | 0 | 1 | 0 | 0 | 0 | 0 | 57.84 | 0 | 0 | 1 |
| 0 | 0 | 0 | 0 | 0 | 1 | 0 | 58.01 | 0 | 0 | 1 |
| 0 | 1 | 1 | 0 | 0 | 0 | 0 | 58.61 | 0 | 0 | 1 |
| 0 | 1 | 0 | 0 | 0 | 1 | 0 | 59.06 | 0 | 0 | 1 |
| 0 | 0 | 1 | 0 | 0 | 1 | 1 | 59.23 | 0 | 0 | 1 |

| Photoperiod | Photo difference | Temperature difference | Flow | Flow difference | Lunar phase | Hatchery | $\Delta AICc$ | rel.LL | weights | Cumulative weights |
|---|---|---|---|---|---|---|---|---|---|---|
| 0 | 0 | 1 | 0 | 0 | 1 | 0 | 59.95 | 0 | 0 | 1 |
| 0 | 1 | 1 | 0 | 0 | 1 | 0 | 60.7 | 0 | 0 | 1 |

Table S5 - Model selection with all combinations of uncorrelated covariates for coho salmon in the Dungeness River.

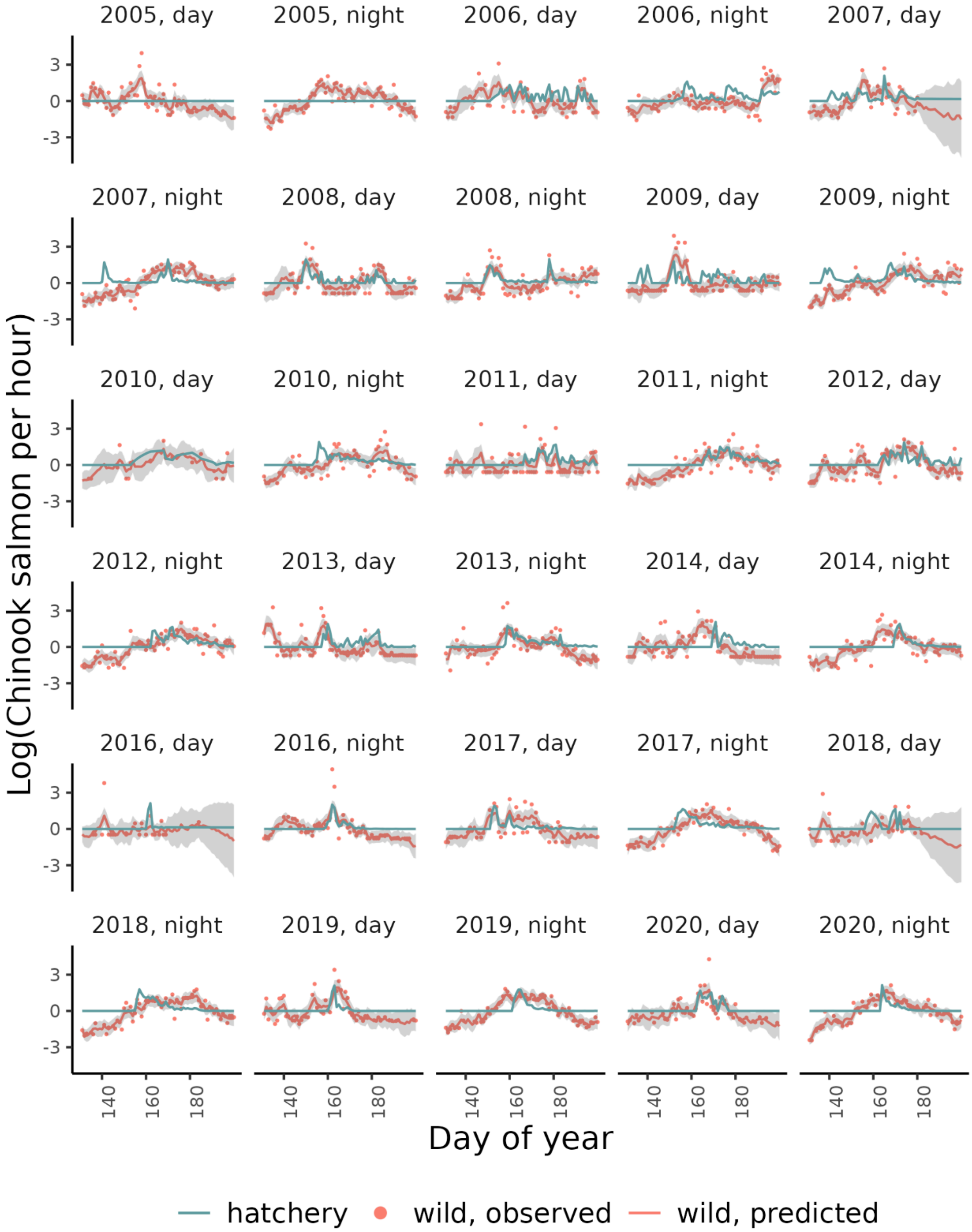

2005, day
2005, night
2006, day
2006, night
2007, day
2007, night
2008, day
2008, night
2009, day
2009, night
2010, day
2010, night
2011, day
2011, night
2012, day
2012, night
2013, day
2013, night
2014, day
2014, night
2016, day
2016, night
2017, day
2017, night
2018, day
2018, night
2019, day
2019, night
2020, day
2020, night
Log(Chinook salmon per hour)
3
0
-3
140
160
180
Day of year
hatchery
wild, observed
wild, predicted

Figure S4 - Model estimates and observations of wild Chinook salmon and observations of hatchery Chinook salmon in the Dungeness River.

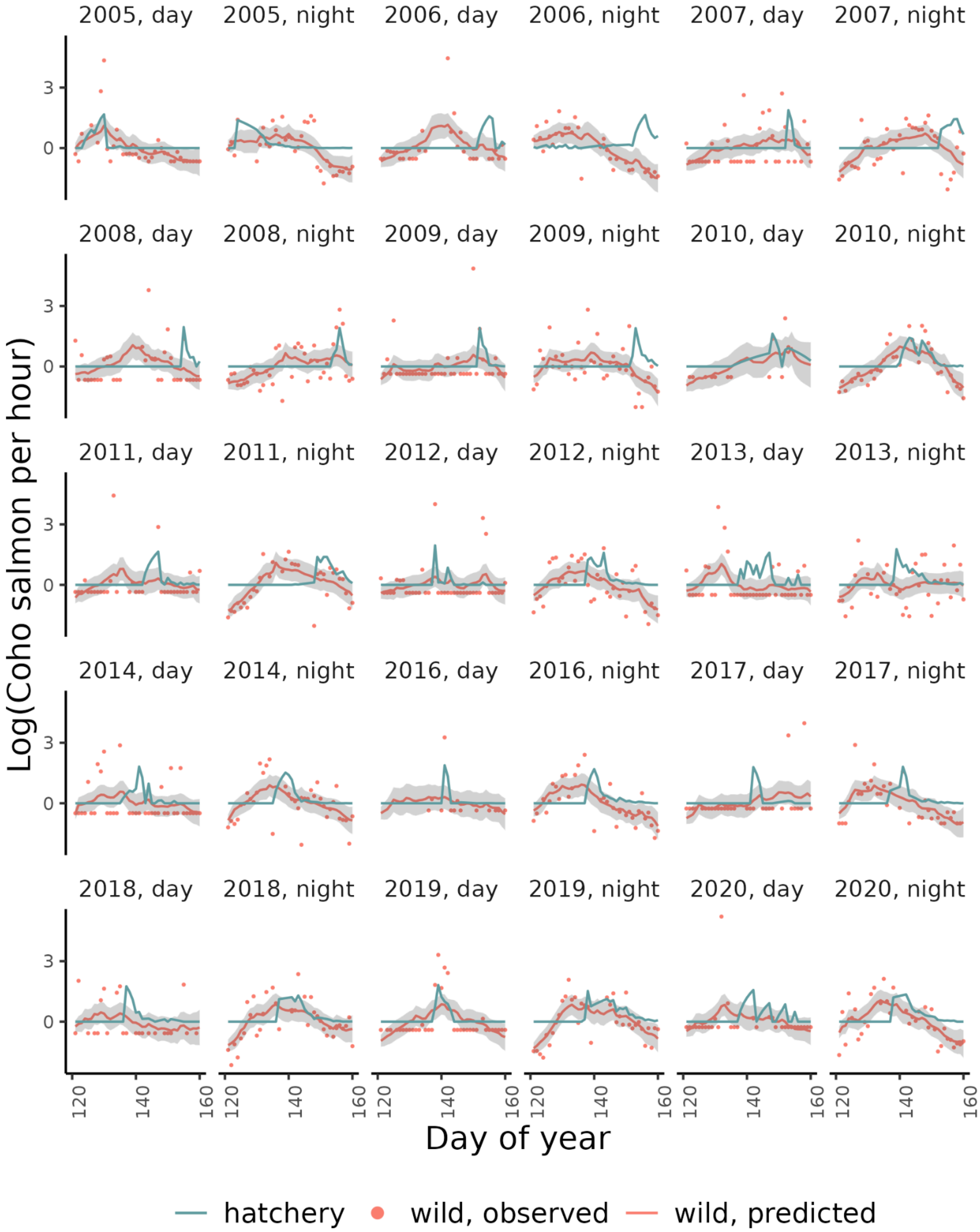
2005, day
2005, night
2006, day
2006, night
2007, day
2007, night
2008, day
2008, night
2009, day
2009, night
2010, day
2010, night
2011, day
2011, night
2012, day
2012, night
2013, day
2013, night
2014, day
2014, night
2016, day
2016, night
2017, day
2017, night
2018, day
2018, night
2019, day
2019, night
2020, day
2020, night
Log(Coho salmon per hour)
3
0
120
140
160
Day of year
hatchery
wild, observed
wild, predicted

Figure S5 - Model estimates and observations of wild coho salmon and observations of hatchery coho salmon in the Dungeness River

### Puyallup River

The Puyallup River is a 72-km river that originates in the glaciers on Mount Rainier and drains into Puget Sound. In the summer, the river is fed by melting glaciers which increases the turbidity. River flow and sediment budget is largely natural, except a diversion dam located in the upper Puyallup which diverts and returns a portion of the river for hydropower. The highest water temperatures are around July-August and the lowest flows are in September-October. The Puyallup River supports Chinook, coho, pink, and chum salmon, and steelhead, cutthroat and bull trout. The Puyallup River trap, operated by the Puyallup Tribe of Indians, is located just upstream of the confluence of the White River and Puyallup River (47.1971, -122.2523). The trap catches hatchery salmon from the Voights Creek hatchery, and Rushingwater, Cowskull, and Wilkeson Creek acclimation ponds and Lake Kapowsin net pen.

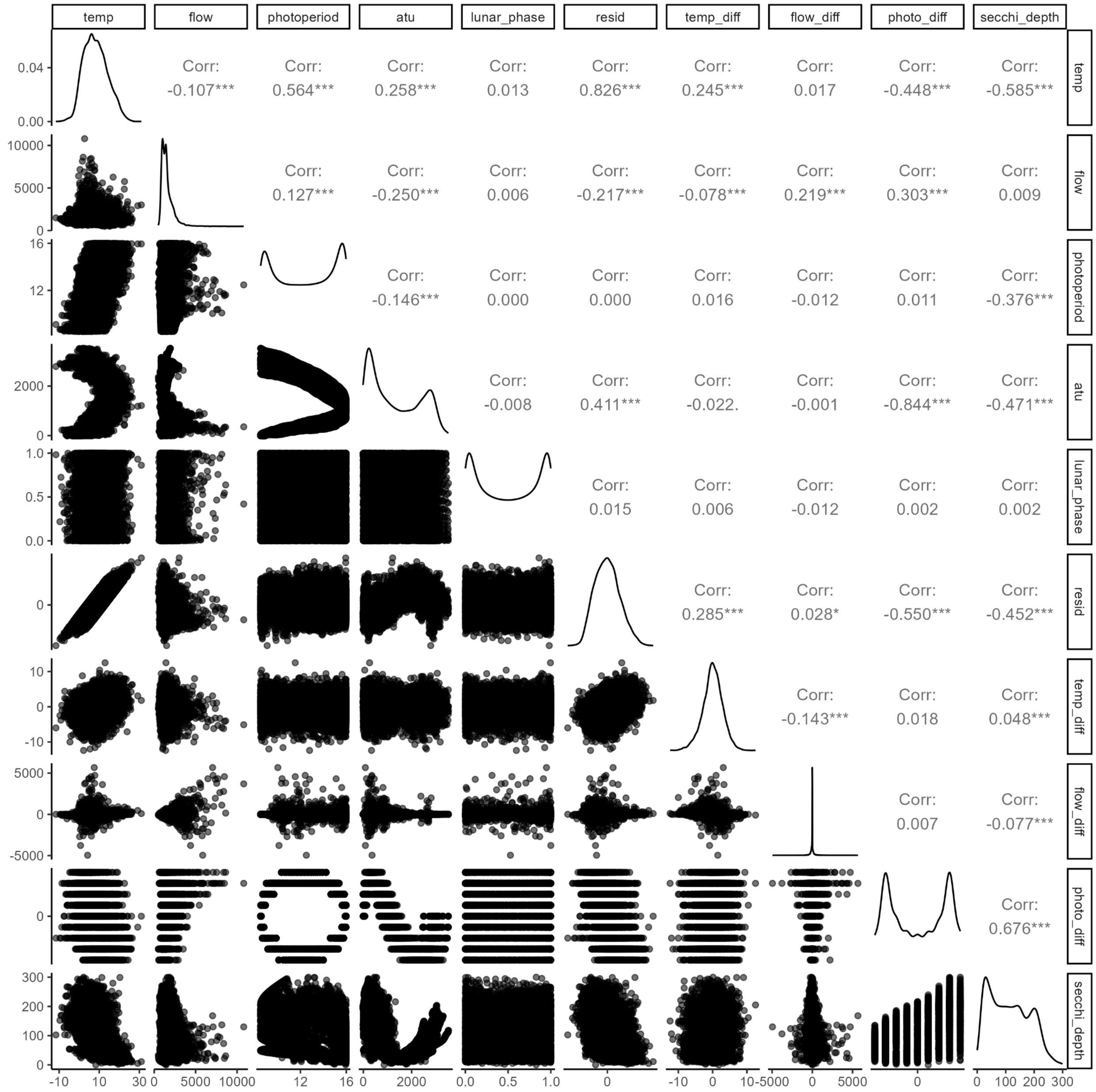


Figure S6 - Correlation between all the environmental variables in the Puyallup River.

| Chinook | | Coho | |
|---|---|---|---|
| **Covariate** | $\Delta AICc$ | **Covariate** | $\Delta AICc$ |
| Photoperiod difference | 0 | ATU | 0 |
| ATU | 1.26 | Photoperiod | 12.51 |
| Residuals | 6.76 | Photoperiod difference | 25.58 |

| | | | |
|---|---|---|---|
| Temperature | 8.33 | Secchi depth | 31.18 |
| Secchi depth | 10.36 | Temperature | 36.51 |
| Photoperiod | 12.22 | Residuals | 36.55 |

Table S6 - Model selection for Chinook and coho salmon in the Puyallup River with each of the correlated covariate.

| Photoperiod difference | Temperature | Temperature difference | Flow | Lunar phase | Flow difference | Hatchery | $\Delta AICc$ | Relative Likelihood | Weights | Cumulative weights |
|---|---|---|---|---|---|---|---|---|---|---|
| 1 | 1 | 0 | 1 | 1 | 1 | 1 | 0 | 1 | 0.5 | 0.5 |
| 1 | 1 | 1 | 1 | 1 | 1 | 1 | 1.43 | 0.49 | 0.24 | 0.74 |
| 1 | 1 | 0 | 1 | 0 | 1 | 1 | 2.17 | 0.34 | 0.17 | 0.91 |
| 1 | 1 | 1 | 1 | 0 | 1 | 1 | 3.68 | 0.16 | 0.08 | 0.99 |
| 1 | 1 | 0 | 1 | 1 | 1 | 0 | 9.66 | 0.01 | 0 | 1 |
| 1 | 1 | 0 | 1 | 0 | 1 | 0 | 11.22 | 0 | 0 | 1 |
| 1 | 1 | 1 | 1 | 1 | 1 | 0 | 11.3 | 0 | 0 | 1 |
| 1 | 1 | 1 | 1 | 0 | 1 | 0 | 12.92 | 0 | 0 | 1 |
| 1 | 1 | 0 | 0 | 1 | 1 | 1 | 18.82 | 0 | 0 | 1 |
| 1 | 1 | 0 | 0 | 0 | 1 | 1 | 19.31 | 0 | 0 | 1 |
| 1 | 1 | 1 | 0 | 1 | 1 | 1 | 20.9 | 0 | 0 | 1 |
| 1 | 1 | 1 | 0 | 0 | 1 | 1 | 21.39 | 0 | 0 | 1 |
| 1 | 1 | 0 | 0 | 1 | 1 | 0 | 23.1 | 0 | 0 | 1 |
| 1 | 1 | 0 | 0 | 0 | 1 | 0 | 23.38 | 0 | 0 | 1 |
| 1 | 1 | 1 | 0 | 1 | 1 | 0 | 25.18 | 0 | 0 | 1 |
| 1 | 1 | 1 | 0 | 0 | 1 | 0 | 25.47 | 0 | 0 | 1 |
| 1 | 0 | 0 | 1 | 1 | 1 | 1 | 48.35 | 0 | 0 | 1 |
| 1 | 0 | 0 | 1 | 0 | 1 | 1 | 48.4 | 0 | 0 | 1 |

| | | | | | | | | | | |
|---|---|---|---|---|---|---|---|---|---|---|
| 0 | 1 | 0 | 0 | 0 | 1 | 1 | 48.51 | 0 | 0 | 1 |
| 0 | 1 | 0 | 0 | 1 | 1 | 1 | 48.99 | 0 | 0 | 1 |
| 0 | 1 | 0 | 1 | 0 | 1 | 1 | 49.18 | 0 | 0 | 1 |
| 0 | 1 | 0 | 1 | 1 | 1 | 1 | 49.46 | 0 | 0 | 1 |
| 1 | 0 | 1 | 1 | 1 | 1 | 1 | 50.26 | 0 | 0 | 1 |
| 1 | 0 | 1 | 1 | 0 | 1 | 1 | 50.28 | 0 | 0 | 1 |
| 0 | 1 | 1 | 0 | 0 | 1 | 1 | 50.51 | 0 | 0 | 1 |
| 0 | 1 | 1 | 0 | 1 | 1 | 1 | 51.01 | 0 | 0 | 1 |
| 0 | 1 | 1 | 1 | 0 | 1 | 1 | 51.23 | 0 | 0 | 1 |
| 0 | 1 | 1 | 1 | 1 | 1 | 1 | 51.53 | 0 | 0 | 1 |
| 1 | 0 | 0 | 1 | 0 | 1 | 0 | 52.22 | 0 | 0 | 1 |
| 1 | 0 | 0 | 1 | 1 | 1 | 0 | 52.35 | 0 | 0 | 1 |
| 1 | 0 | 1 | 1 | 0 | 1 | 0 | 54.04 | 0 | 0 | 1 |
| 1 | 0 | 1 | 1 | 1 | 1 | 0 | 54.2 | 0 | 0 | 1 |
| 0 | 0 | 0 | 1 | 0 | 1 | 1 | 55.08 | 0 | 0 | 1 |
| 1 | 0 | 0 | 0 | 0 | 1 | 1 | 55.25 | 0 | 0 | 1 |
| 0 | 0 | 0 | 1 | 1 | 1 | 1 | 55.47 | 0 | 0 | 1 |
| 1 | 0 | 0 | 0 | 1 | 1 | 1 | 55.89 | 0 | 0 | 1 |
| 0 | 0 | 0 | 0 | 0 | 1 | 1 | 56.15 | 0 | 0 | 1 |
| 1 | 0 | 0 | 0 | 0 | 1 | 0 | 56.5 | 0 | 0 | 1 |

| | | | | | | | | | | |
|---|---|---|---|---|---|---|---|---|---|---|
| 1 | 0 | 1 | 0 | 0 | 1 | 1 | 56.73 | 0 | 0 | 1 |
| 0 | 0 | 0 | 0 | 1 | 1 | 1 | 56.85 | 0 | 0 | 1 |
| 0 | 0 | 1 | 1 | 0 | 1 | 1 | 56.88 | 0 | 0 | 1 |
| 1 | 0 | 0 | 0 | 1 | 1 | 0 | 57.18 | 0 | 0 | 1 |
| 0 | 0 | 1 | 1 | 1 | 1 | 1 | 57.31 | 0 | 0 | 1 |
| 1 | 0 | 1 | 0 | 1 | 1 | 1 | 57.4 | 0 | 0 | 1 |
| 0 | 0 | 1 | 0 | 0 | 1 | 1 | 57.72 | 0 | 0 | 1 |
| 1 | 0 | 1 | 0 | 0 | 1 | 0 | 57.97 | 0 | 0 | 1 |
| 0 | 0 | 1 | 0 | 1 | 1 | 1 | 58.44 | 0 | 0 | 1 |
| 1 | 0 | 1 | 0 | 1 | 1 | 0 | 58.66 | 0 | 0 | 1 |
| 0 | 1 | 0 | 0 | 0 | 1 | 0 | 60.82 | 0 | 0 | 1 |
| 0 | 1 | 0 | 0 | 1 | 1 | 0 | 61.57 | 0 | 0 | 1 |
| 0 | 0 | 0 | 0 | 0 | 1 | 0 | 62.42 | 0 | 0 | 1 |
| 0 | 1 | 1 | 0 | 0 | 1 | 0 | 62.77 | 0 | 0 | 1 |
| 0 | 1 | 0 | 1 | 0 | 1 | 0 | 62.9 | 0 | 0 | 1 |
| 0 | 0 | 0 | 0 | 1 | 1 | 0 | 63.24 | 0 | 0 | 1 |
| 0 | 1 | 1 | 0 | 1 | 1 | 0 | 63.54 | 0 | 0 | 1 |
| 0 | 1 | 0 | 1 | 1 | 1 | 0 | 63.64 | 0 | 0 | 1 |
| 0 | 0 | 1 | 0 | 0 | 1 | 0 | 64.07 | 0 | 0 | 1 |
| 0 | 0 | 0 | 1 | 0 | 1 | 0 | 64.17 | 0 | 0 | 1 |

| | | | | | | | | | | |
|---|---|---|---|---|---|---|---|---|---|---|
| 0 | 1 | 1 | 1 | 0 | 1 | 0 | 64.86 | 0 | 0 | 1 |
| 0 | 0 | 0 | 1 | 1 | 1 | 0 | 64.91 | 0 | 0 | 1 |
| 0 | 0 | 1 | 0 | 1 | 1 | 0 | 64.91 | 0 | 0 | 1 |
| 0 | 1 | 1 | 1 | 1 | 1 | 0 | 65.62 | 0 | 0 | 1 |
| 0 | 0 | 1 | 1 | 0 | 1 | 0 | 65.91 | 0 | 0 | 1 |
| 0 | 0 | 1 | 1 | 1 | 1 | 0 | 66.68 | 0 | 0 | 1 |
| 1 | 1 | 0 | 1 | 1 | 0 | 1 | 287.11 | 0 | 0 | 1 |
| 1 | 1 | 1 | 1 | 1 | 0 | 1 | 288.19 | 0 | 0 | 1 |
| 1 | 1 | 0 | 1 | 1 | 0 | 0 | 290.91 | 0 | 0 | 1 |
| 1 | 1 | 1 | 1 | 1 | 0 | 0 | 292.59 | 0 | 0 | 1 |
| 1 | 1 | 0 | 1 | 0 | 0 | 1 | 293.06 | 0 | 0 | 1 |
| 1 | 1 | 1 | 1 | 0 | 0 | 1 | 294.14 | 0 | 0 | 1 |
| 1 | 1 | 0 | 1 | 0 | 0 | 0 | 296.39 | 0 | 0 | 1 |
| 1 | 1 | 1 | 1 | 0 | 0 | 0 | 298.06 | 0 | 0 | 1 |
| 1 | 1 | 0 | 0 | 1 | 0 | 1 | 298.26 | 0 | 0 | 1 |
| 1 | 1 | 0 | 0 | 1 | 0 | 0 | 298.61 | 0 | 0 | 1 |
| 1 | 1 | 1 | 0 | 1 | 0 | 1 | 300.13 | 0 | 0 | 1 |
| 1 | 1 | 1 | 0 | 1 | 0 | 0 | 300.65 | 0 | 0 | 1 |
| 1 | 1 | 0 | 0 | 0 | 0 | 1 | 302.13 | 0 | 0 | 1 |
| 1 | 1 | 0 | 0 | 0 | 0 | 0 | 302.46 | 0 | 0 | 1 |

| | | | | | | | | | | |
|---|---|---|---|---|---|---|---|---|---|---|
| 1 | 1 | 1 | 0 | 0 | 0 | 1 | 303.97 | 0 | 0 | 1 |
| 1 | 1 | 1 | 0 | 0 | 0 | 0 | 304.47 | 0 | 0 | 1 |
| 0 | 1 | 0 | 0 | 1 | 0 | 1 | 357.78 | 0 | 0 | 1 |
| 0 | 1 | 1 | 0 | 1 | 0 | 1 | 359.25 | 0 | 0 | 1 |
| 0 | 1 | 0 | 0 | 0 | 0 | 1 | 359.29 | 0 | 0 | 1 |
| 0 | 1 | 0 | 1 | 1 | 0 | 1 | 359.41 | 0 | 0 | 1 |
| 0 | 1 | 0 | 1 | 0 | 0 | 1 | 360.73 | 0 | 0 | 1 |
| 0 | 1 | 1 | 0 | 0 | 0 | 1 | 360.74 | 0 | 0 | 1 |
| 0 | 1 | 1 | 1 | 1 | 0 | 1 | 361.02 | 0 | 0 | 1 |
| 0 | 1 | 1 | 1 | 0 | 0 | 1 | 362.34 | 0 | 0 | 1 |
| 1 | 0 | 0 | 0 | 1 | 0 | 0 | 362.92 | 0 | 0 | 1 |
| 1 | 0 | 0 | 1 | 1 | 0 | 0 | 363.41 | 0 | 0 | 1 |
| 1 | 0 | 1 | 0 | 1 | 0 | 0 | 364.39 | 0 | 0 | 1 |
| 1 | 0 | 0 | 0 | 0 | 0 | 0 | 364.84 | 0 | 0 | 1 |
| 1 | 0 | 1 | 1 | 1 | 0 | 0 | 365.04 | 0 | 0 | 1 |
| 1 | 0 | 0 | 1 | 1 | 0 | 1 | 365.2 | 0 | 0 | 1 |
| 1 | 0 | 0 | 0 | 1 | 0 | 1 | 365.42 | 0 | 0 | 1 |
| 1 | 0 | 0 | 1 | 0 | 0 | 0 | 365.79 | 0 | 0 | 1 |
| 1 | 0 | 1 | 0 | 0 | 0 | 0 | 366.36 | 0 | 0 | 1 |
| 1 | 0 | 1 | 1 | 1 | 0 | 1 | 367.12 | 0 | 0 | 1 |

| | | | | | | | | | | |
|---|---|---|---|---|---|---|---|---|---|---|
| 1 | 0 | 1 | 0 | 1 | 0 | 1 | 367.16 | 0 | 0 | 1 |
| 1 | 0 | 0 | 0 | 0 | 0 | 1 | 367.2 | 0 | 0 | 1 |
| 1 | 0 | 1 | 1 | 0 | 0 | 0 | 367.44 | 0 | 0 | 1 |
| 1 | 0 | 0 | 1 | 0 | 0 | 1 | 367.54 | 0 | 0 | 1 |
| 1 | 0 | 1 | 0 | 0 | 0 | 1 | 368.98 | 0 | 0 | 1 |
| 1 | 0 | 1 | 1 | 0 | 0 | 1 | 369.47 | 0 | 0 | 1 |
| 0 | 1 | 0 | 1 | 1 | 0 | 0 | 370.25 | 0 | 0 | 1 |
| 0 | 1 | 0 | 1 | 0 | 0 | 0 | 371.16 | 0 | 0 | 1 |
| 0 | 1 | 0 | 0 | 1 | 0 | 0 | 371.8 | 0 | 0 | 1 |
| 0 | 0 | 0 | 0 | 1 | 0 | 1 | 372.01 | 0 | 0 | 1 |
| 0 | 1 | 1 | 1 | 1 | 0 | 0 | 372.27 | 0 | 0 | 1 |
| 0 | 1 | 0 | 0 | 0 | 0 | 0 | 373.16 | 0 | 0 | 1 |
| 0 | 1 | 1 | 1 | 0 | 0 | 0 | 373.18 | 0 | 0 | 1 |
| 0 | 0 | 0 | 0 | 0 | 0 | 1 | 373.4 | 0 | 0 | 1 |
| 0 | 1 | 1 | 0 | 1 | 0 | 0 | 373.62 | 0 | 0 | 1 |
| 0 | 0 | 1 | 0 | 1 | 0 | 1 | 374.08 | 0 | 0 | 1 |
| 0 | 0 | 0 | 1 | 1 | 0 | 1 | 374.09 | 0 | 0 | 1 |
| 0 | 1 | 1 | 0 | 0 | 0 | 0 | 374.96 | 0 | 0 | 1 |
| 0 | 0 | 0 | 1 | 0 | 0 | 1 | 375.45 | 0 | 0 | 1 |
| 0 | 0 | 1 | 0 | 0 | 0 | 1 | 375.48 | 0 | 0 | 1 |

| | | | | | | | | | | |
|---|---|---|---|---|---|---|---|---|---|---|
| 0 | 0 | 1 | 1 | 1 | 0 | 1 | 376.17 | 0 | 0 | 1 |
| 0 | 0 | 0 | 0 | 1 | 0 | 0 | 376.96 | 0 | 0 | 1 |
| 0 | 0 | 1 | 1 | 0 | 0 | 1 | 377.53 | 0 | 0 | 1 |
| 0 | 0 | 0 | 1 | 1 | 0 | 0 | 377.65 | 0 | 0 | 1 |
| 0 | 0 | 0 | 0 | 0 | 0 | 0 | 378.35 | 0 | 0 | 1 |
| 0 | 0 | 0 | 0 | 0 | 0 | 0 | 378.35 | 0 | 0 | 1 |
| 0 | 0 | 0 | 1 | 0 | 0 | 0 | 378.75 | 0 | 0 | 1 |
| 0 | 0 | 1 | 0 | 1 | 0 | 0 | 379.02 | 0 | 0 | 1 |
| 0 | 0 | 1 | 1 | 1 | 0 | 0 | 379.59 | 0 | 0 | 1 |
| 0 | 0 | 1 | 0 | 0 | 0 | 0 | 380.41 | 0 | 0 | 1 |
| 0 | 0 | 1 | 1 | 0 | 0 | 0 | 380.7 | 0 | 0 | 1 |

Table S7 - Model selection with all combinations of uncorrelated covariates for Chinook salmon in the Puyallup River.

| Photoperiod difference | Temperature | Temperature difference | Flow | Lunar phase | Flow difference | Hatchery | $\Delta AICc$ | Relative Likelihood | Weights | Cumulative weights |
|---|---|---|---|---|---|---|---|---|---|---|
| 1 | 1 | 0 | 1 | 1 | 1 | 1 | 0 | 1 | 0.5 | 0.5 |
| 1 | 1 | 1 | 1 | 1 | 1 | 1 | 1.43 | 0.49 | 0.24 | 0.74 |
| 1 | 1 | 0 | 1 | 0 | 1 | 1 | 2.17 | 0.34 | 0.17 | 0.91 |
| 1 | 1 | 1 | 1 | 0 | 1 | 1 | 3.68 | 0.16 | 0.08 | 0.99 |
| 1 | 1 | 0 | 1 | 1 | 1 | 0 | 9.66 | 0.01 | 0 | 1 |

| | | | | | | | | | | |
|---|---|---|---|---|---|---|---|---|---|---|
| 1 | 1 | 0 | 1 | 0 | 1 | 0 | 11.22 | 0 | 0 | 1 |
| 1 | 1 | 1 | 1 | 1 | 1 | 0 | 11.3 | 0 | 0 | 1 |
| 1 | 1 | 1 | 1 | 0 | 1 | 0 | 12.92 | 0 | 0 | 1 |
| 1 | 1 | 0 | 0 | 1 | 1 | 1 | 18.82 | 0 | 0 | 1 |
| 1 | 1 | 0 | 0 | 0 | 1 | 1 | 19.31 | 0 | 0 | 1 |
| 1 | 1 | 1 | 0 | 1 | 1 | 1 | 20.9 | 0 | 0 | 1 |
| 1 | 1 | 1 | 0 | 0 | 1 | 1 | 21.39 | 0 | 0 | 1 |
| 1 | 1 | 0 | 0 | 1 | 1 | 0 | 23.1 | 0 | 0 | 1 |
| 1 | 1 | 0 | 0 | 0 | 1 | 0 | 23.38 | 0 | 0 | 1 |
| 1 | 1 | 1 | 0 | 1 | 1 | 0 | 25.18 | 0 | 0 | 1 |
| 1 | 1 | 1 | 0 | 0 | 1 | 0 | 25.47 | 0 | 0 | 1 |
| 1 | 0 | 0 | 1 | 1 | 1 | 1 | 48.35 | 0 | 0 | 1 |
| 1 | 0 | 0 | 1 | 0 | 1 | 1 | 48.4 | 0 | 0 | 1 |
| 0 | 1 | 0 | 0 | 0 | 1 | 1 | 48.51 | 0 | 0 | 1 |
| 0 | 1 | 0 | 0 | 1 | 1 | 1 | 48.99 | 0 | 0 | 1 |
| 0 | 1 | 0 | 1 | 0 | 1 | 1 | 49.18 | 0 | 0 | 1 |
| 0 | 1 | 0 | 1 | 1 | 1 | 1 | 49.46 | 0 | 0 | 1 |
| 1 | 0 | 1 | 1 | 1 | 1 | 1 | 50.26 | 0 | 0 | 1 |
| 1 | 0 | 1 | 1 | 0 | 1 | 1 | 50.28 | 0 | 0 | 1 |
| 0 | 1 | 1 | 0 | 0 | 1 | 1 | 50.51 | 0 | 0 | 1 |

| | | | | | | | | | | |
|---|---|---|---|---|---|---|---|---|---|---|
| 0 | 1 | 1 | 0 | 1 | 1 | 1 | 51.01 | 0 | 0 | 1 |
| 0 | 1 | 1 | 1 | 0 | 1 | 1 | 51.23 | 0 | 0 | 1 |
| 0 | 1 | 1 | 1 | 1 | 1 | 1 | 51.53 | 0 | 0 | 1 |
| 1 | 0 | 0 | 1 | 0 | 1 | 0 | 52.22 | 0 | 0 | 1 |
| 1 | 0 | 0 | 1 | 1 | 1 | 0 | 52.35 | 0 | 0 | 1 |
| 1 | 0 | 1 | 1 | 0 | 1 | 0 | 54.04 | 0 | 0 | 1 |
| 1 | 0 | 1 | 1 | 1 | 1 | 0 | 54.2 | 0 | 0 | 1 |
| 0 | 0 | 0 | 1 | 0 | 1 | 1 | 55.08 | 0 | 0 | 1 |
| 1 | 0 | 0 | 0 | 0 | 1 | 1 | 55.25 | 0 | 0 | 1 |
| 0 | 0 | 0 | 1 | 1 | 1 | 1 | 55.47 | 0 | 0 | 1 |
| 1 | 0 | 0 | 0 | 1 | 1 | 1 | 55.89 | 0 | 0 | 1 |
| 0 | 0 | 0 | 0 | 0 | 1 | 1 | 56.15 | 0 | 0 | 1 |
| 1 | 0 | 0 | 0 | 0 | 1 | 0 | 56.5 | 0 | 0 | 1 |
| 1 | 0 | 1 | 0 | 0 | 1 | 1 | 56.73 | 0 | 0 | 1 |
| 0 | 0 | 0 | 0 | 1 | 1 | 1 | 56.85 | 0 | 0 | 1 |
| 0 | 0 | 1 | 1 | 0 | 1 | 1 | 56.88 | 0 | 0 | 1 |
| 1 | 0 | 0 | 0 | 1 | 1 | 0 | 57.18 | 0 | 0 | 1 |
| 0 | 0 | 1 | 1 | 1 | 1 | 1 | 57.31 | 0 | 0 | 1 |
| 1 | 0 | 1 | 0 | 1 | 1 | 1 | 57.4 | 0 | 0 | 1 |
| 0 | 0 | 1 | 0 | 0 | 1 | 1 | 57.72 | 0 | 0 | 1 |

| | | | | | | | | | | |
|---|---|---|---|---|---|---|---|---|---|---|
| 1 | 0 | 1 | 0 | 0 | 1 | 0 | 57.97 | 0 | 0 | 1 |
| 0 | 0 | 1 | 0 | 1 | 1 | 1 | 58.44 | 0 | 0 | 1 |
| 1 | 0 | 1 | 0 | 1 | 1 | 0 | 58.66 | 0 | 0 | 1 |
| 0 | 1 | 0 | 0 | 0 | 1 | 0 | 60.82 | 0 | 0 | 1 |
| 0 | 1 | 0 | 0 | 1 | 1 | 0 | 61.57 | 0 | 0 | 1 |
| 0 | 0 | 0 | 0 | 0 | 1 | 0 | 62.42 | 0 | 0 | 1 |
| 0 | 1 | 1 | 0 | 0 | 1 | 0 | 62.77 | 0 | 0 | 1 |
| 0 | 1 | 0 | 1 | 0 | 1 | 0 | 62.9 | 0 | 0 | 1 |
| 0 | 0 | 0 | 0 | 1 | 1 | 0 | 63.24 | 0 | 0 | 1 |
| 0 | 1 | 1 | 0 | 1 | 1 | 0 | 63.54 | 0 | 0 | 1 |
| 0 | 1 | 0 | 1 | 1 | 1 | 0 | 63.64 | 0 | 0 | 1 |
| 0 | 0 | 1 | 0 | 0 | 1 | 0 | 64.07 | 0 | 0 | 1 |
| 0 | 0 | 0 | 1 | 0 | 1 | 0 | 64.17 | 0 | 0 | 1 |
| 0 | 1 | 1 | 1 | 0 | 1 | 0 | 64.86 | 0 | 0 | 1 |
| 0 | 0 | 0 | 1 | 1 | 1 | 0 | 64.91 | 0 | 0 | 1 |
| 0 | 0 | 1 | 0 | 1 | 1 | 0 | 64.91 | 0 | 0 | 1 |
| 0 | 1 | 1 | 1 | 1 | 1 | 0 | 65.62 | 0 | 0 | 1 |
| 0 | 0 | 1 | 1 | 0 | 1 | 0 | 65.91 | 0 | 0 | 1 |
| 0 | 0 | 1 | 1 | 1 | 1 | 0 | 66.68 | 0 | 0 | 1 |
| 1 | 1 | 0 | 1 | 1 | 0 | 1 | 287.11 | 0 | 0 | 1 |

| | | | | | | | | | | |
|---|---|---|---|---|---|---|---|---|---|---|
| 1 | 1 | 1 | 1 | 1 | 0 | 1 | 288.19 | 0 | 0 | 1 |
| 1 | 1 | 0 | 1 | 1 | 0 | 0 | 290.91 | 0 | 0 | 1 |
| 1 | 1 | 1 | 1 | 1 | 0 | 0 | 292.59 | 0 | 0 | 1 |
| 1 | 1 | 0 | 1 | 0 | 0 | 1 | 293.06 | 0 | 0 | 1 |
| 1 | 1 | 1 | 1 | 0 | 0 | 1 | 294.14 | 0 | 0 | 1 |
| 1 | 1 | 0 | 1 | 0 | 0 | 0 | 296.39 | 0 | 0 | 1 |
| 1 | 1 | 1 | 1 | 0 | 0 | 0 | 298.06 | 0 | 0 | 1 |
| 1 | 1 | 0 | 0 | 1 | 0 | 1 | 298.26 | 0 | 0 | 1 |
| 1 | 1 | 0 | 0 | 1 | 0 | 0 | 298.61 | 0 | 0 | 1 |
| 1 | 1 | 1 | 0 | 1 | 0 | 1 | 300.13 | 0 | 0 | 1 |
| 1 | 1 | 1 | 0 | 1 | 0 | 0 | 300.65 | 0 | 0 | 1 |
| 1 | 1 | 0 | 0 | 0 | 0 | 1 | 302.13 | 0 | 0 | 1 |
| 1 | 1 | 0 | 0 | 0 | 0 | 0 | 302.46 | 0 | 0 | 1 |
| 1 | 1 | 1 | 0 | 0 | 0 | 1 | 303.97 | 0 | 0 | 1 |
| 1 | 1 | 1 | 0 | 0 | 0 | 0 | 304.47 | 0 | 0 | 1 |
| 0 | 1 | 0 | 0 | 1 | 0 | 1 | 357.78 | 0 | 0 | 1 |
| 0 | 1 | 1 | 0 | 1 | 0 | 1 | 359.25 | 0 | 0 | 1 |
| 0 | 1 | 0 | 0 | 0 | 0 | 1 | 359.29 | 0 | 0 | 1 |
| 0 | 1 | 0 | 1 | 1 | 0 | 1 | 359.41 | 0 | 0 | 1 |
| 0 | 1 | 0 | 1 | 0 | 0 | 1 | 360.73 | 0 | 0 | 1 |

| | | | | | | | | | | |
|---|---|---|---|---|---|---|---|---|---|---|
| 0 | 1 | 1 | 0 | 0 | 0 | 1 | 360.74 | 0 | 0 | 1 |
| 0 | 1 | 1 | 1 | 1 | 0 | 1 | 361.02 | 0 | 0 | 1 |
| 0 | 1 | 1 | 1 | 0 | 0 | 1 | 362.34 | 0 | 0 | 1 |
| 1 | 0 | 0 | 0 | 1 | 0 | 0 | 362.92 | 0 | 0 | 1 |
| 1 | 0 | 0 | 1 | 1 | 0 | 0 | 363.41 | 0 | 0 | 1 |
| 1 | 0 | 1 | 0 | 1 | 0 | 0 | 364.39 | 0 | 0 | 1 |
| 1 | 0 | 0 | 0 | 0 | 0 | 0 | 364.84 | 0 | 0 | 1 |
| 1 | 0 | 1 | 1 | 1 | 0 | 0 | 365.04 | 0 | 0 | 1 |
| 1 | 0 | 0 | 1 | 1 | 0 | 1 | 365.2 | 0 | 0 | 1 |
| 1 | 0 | 0 | 0 | 1 | 0 | 1 | 365.42 | 0 | 0 | 1 |
| 1 | 0 | 0 | 1 | 0 | 0 | 0 | 365.79 | 0 | 0 | 1 |
| 1 | 0 | 1 | 0 | 0 | 0 | 0 | 366.36 | 0 | 0 | 1 |
| 1 | 0 | 1 | 1 | 1 | 0 | 1 | 367.12 | 0 | 0 | 1 |
| 1 | 0 | 1 | 0 | 1 | 0 | 1 | 367.16 | 0 | 0 | 1 |
| 1 | 0 | 0 | 0 | 0 | 0 | 1 | 367.2 | 0 | 0 | 1 |
| 1 | 0 | 1 | 1 | 0 | 0 | 0 | 367.44 | 0 | 0 | 1 |
| 1 | 0 | 0 | 1 | 0 | 0 | 1 | 367.54 | 0 | 0 | 1 |
| 1 | 0 | 1 | 0 | 0 | 0 | 1 | 368.98 | 0 | 0 | 1 |
| 1 | 0 | 1 | 1 | 0 | 0 | 1 | 369.47 | 0 | 0 | 1 |
| 0 | 1 | 0 | 1 | 1 | 0 | 0 | 370.25 | 0 | 0 | 1 |

| | | | | | | | | | | |
|---|---|---|---|---|---|---|---|---|---|---|
| 0 | 1 | 0 | 1 | 0 | 0 | 0 | 371.16 | 0 | 0 | 1 |
| 0 | 1 | 0 | 0 | 1 | 0 | 0 | 371.8 | 0 | 0 | 1 |
| 0 | 0 | 0 | 0 | 1 | 0 | 1 | 372.01 | 0 | 0 | 1 |
| 0 | 1 | 1 | 1 | 1 | 0 | 0 | 372.27 | 0 | 0 | 1 |
| 0 | 1 | 0 | 0 | 0 | 0 | 0 | 373.16 | 0 | 0 | 1 |
| 0 | 1 | 1 | 1 | 0 | 0 | 0 | 373.18 | 0 | 0 | 1 |
| 0 | 0 | 0 | 0 | 0 | 0 | 1 | 373.4 | 0 | 0 | 1 |
| 0 | 1 | 1 | 0 | 1 | 0 | 0 | 373.62 | 0 | 0 | 1 |
| 0 | 0 | 1 | 0 | 1 | 0 | 1 | 374.08 | 0 | 0 | 1 |
| 0 | 0 | 0 | 1 | 1 | 0 | 1 | 374.09 | 0 | 0 | 1 |
| 0 | 1 | 1 | 0 | 0 | 0 | 0 | 374.96 | 0 | 0 | 1 |
| 0 | 0 | 0 | 1 | 0 | 0 | 1 | 375.45 | 0 | 0 | 1 |
| 0 | 0 | 1 | 0 | 0 | 0 | 1 | 375.48 | 0 | 0 | 1 |
| 0 | 0 | 1 | 1 | 1 | 0 | 1 | 376.17 | 0 | 0 | 1 |
| 0 | 0 | 0 | 0 | 1 | 0 | 0 | 376.96 | 0 | 0 | 1 |
| 0 | 0 | 1 | 1 | 0 | 0 | 1 | 377.53 | 0 | 0 | 1 |
| 0 | 0 | 0 | 1 | 1 | 0 | 0 | 377.65 | 0 | 0 | 1 |
| 0 | 0 | 0 | 0 | 0 | 0 | 0 | 378.35 | 0 | 0 | 1 |
| 0 | 0 | 0 | 0 | 0 | 0 | 0 | 378.35 | 0 | 0 | 1 |
| 0 | 0 | 0 | 1 | 0 | 0 | 0 | 378.75 | 0 | 0 | 1 |

| | | | | | | | | | | |
|---|---|---|---|---|---|---|---|---|---|---|
| 0 | 0 | 1 | 0 | 1 | 0 | 0 | 379.02 | 0 | 0 | 1 |
| 0 | 0 | 1 | 1 | 1 | 0 | 0 | 379.59 | 0 | 0 | 1 |
| 0 | 0 | 1 | 0 | 0 | 0 | 0 | 380.41 | 0 | 0 | 1 |
| 0 | 0 | 1 | 1 | 0 | 0 | 0 | 380.7 | 0 | 0 | 1 |

Table S8 - Model selection with all combinations of uncorrelated covariates for coho salmon in the Puyallup River.

| **Chinook** | | **Coho** | |
|---|---|---|---|
| **Variable** | **Relative Importance** | **Variable** | **Relative Importance** |
| Flow difference | 1 | Flow difference | 1 |
| Temperature | 1 | Flow | 1 |
| Flow | 1 | Hatchery | 1 |
| Hatchery | 1 | ATU | 1 |
| Photoperiod difference | 1 | Photoperiod | 0.9 |
| Lunar Phase | 0.4 | Lunar phase | 0.5 |
| Temperature difference | 0.3 | Temperature difference | 0.3 |

Table S9 - Relative variable importance for all variables used in the model selection process for Chinook and coho salmon in the Puyallup River.

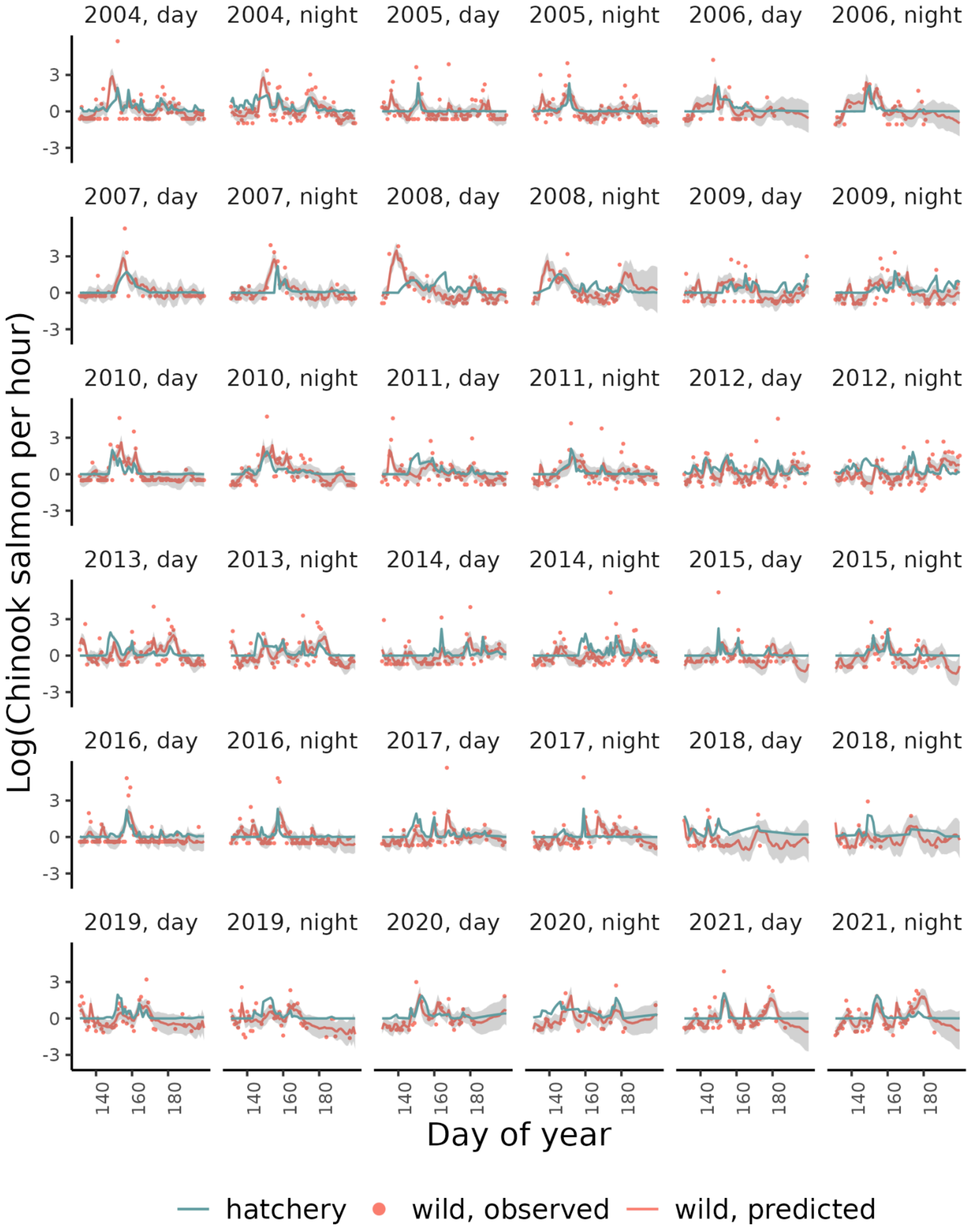
2004, day
2004, night
2005, day
2005, night
2006, day
2006, night
2007, day
2007, night
2008, day
2008, night
2009, day
2009, night
2010, day
2010, night
2011, day
2011, night
2012, day
2012, night
2013, day
2013, night
2014, day
2014, night
2015, day
2015, night
2016, day
2016, night
2017, day
2017, night
2018, day
2018, night
2019, day
2019, night
2020, day
2020, night
2021, day
2021, night
Log(Chinook salmon per hour)
3
0
-3
140
160
180
Day of year
hatchery
wild, observed
wild, predicted

Figure S7 - Model estimates and observations of wild Chinook salmon and observations of hatchery Chinook salmon in the Puyallup River.

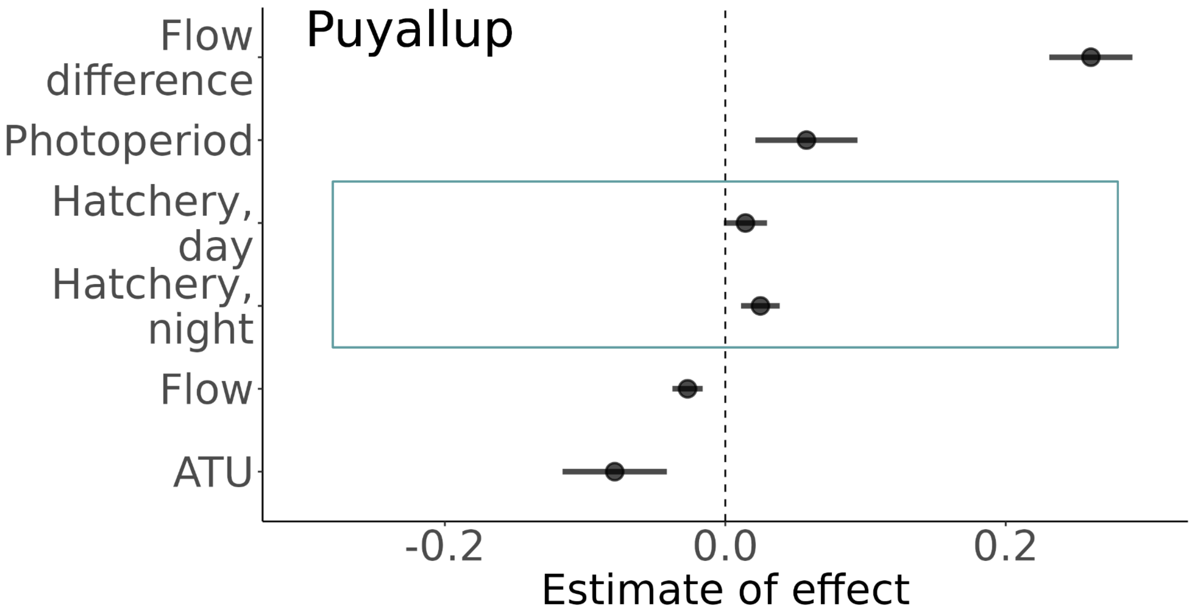


Figure S8 - Estimates of the effect of different covariates in the best MARSS model for coho salmon in the Puyallup River without 2009 and 2021 data (years with high proportions of unmarked hatchery fish). Blue boxes indicate social variables associated with the pied piper hypothesis.

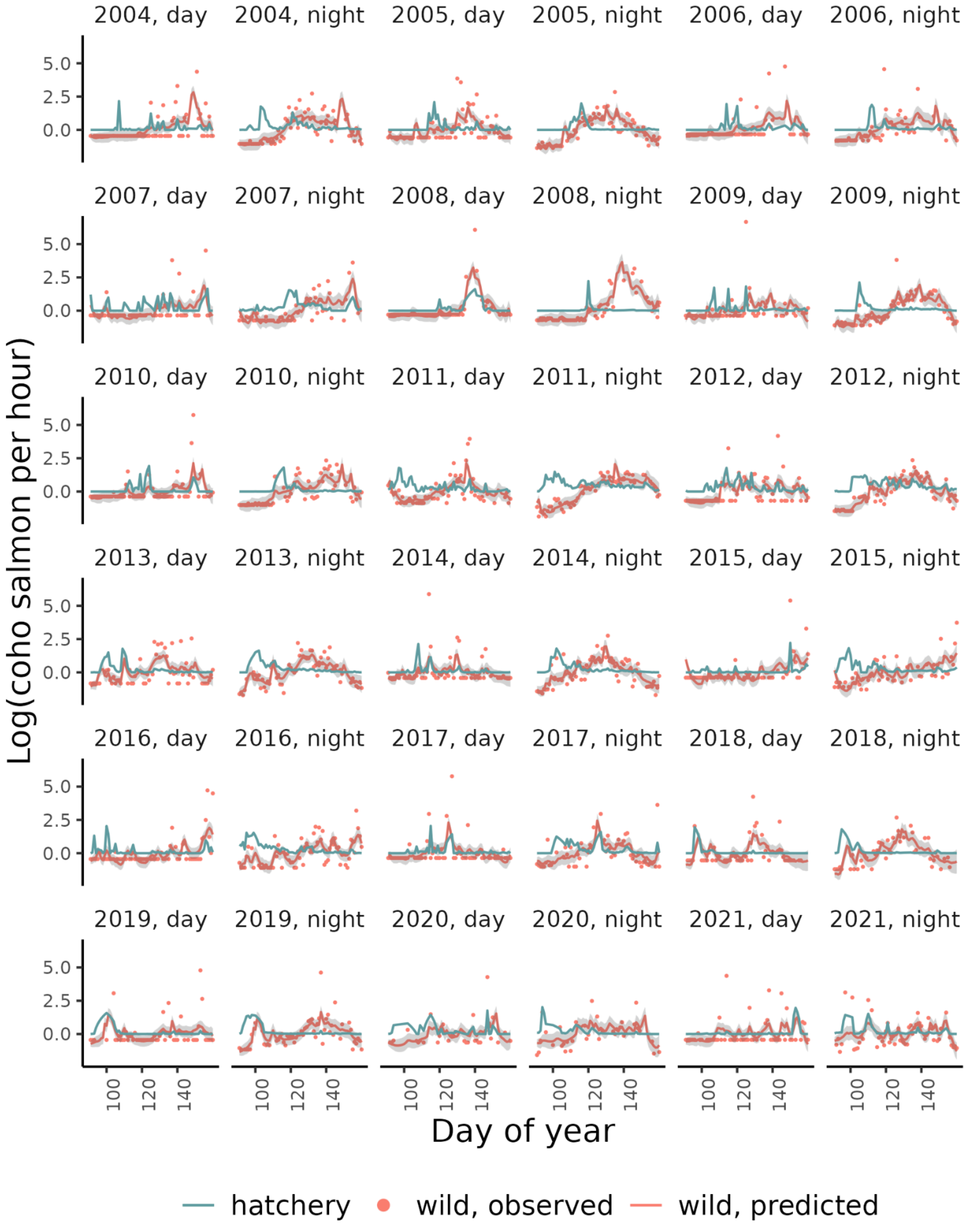
2004, day
2004, night
2005, day
2005, night
2006, day
2006, night
2007, day
2007, night
2008, day
2008, night
2009, day
2009, night
2010, day
2010, night
2011, day
2011, night
2012, day
2012, night
2013, day
2013, night
2014, day
2014, night
2015, day
2015, night
2016, day
2016, night
2017, day
2017, night
2018, day
2018, night
2019, day
2019, night
2020, day
2020, night
2021, day
2021, night
5.0
2.5
0.0
100
120
140
Log(coho salmon per hour)
Day of year
hatchery
wild, observed
wild, predicted

Figure S9 - Model estimates and observations of wild coho salmon and observations of hatchery coho salmon in the Puyallup River.

### Skagit River

The Skagit River is a 240-km river that originates in the Canadian Cascades and drains into the Puget Sound. Several dams regulate the flow of the river. The Skagit River supports all five species of Pacific Salmon as well as Steelhead and Cutthroat. WDFW operates a screw trap and a scoop trap on the mainstem of the Skagit River (48.4451, -122.3251), 27 km upstream of the river mouth. The traps catch hatchery salmon released from Countyline Ponds, Baker Lake Hatchery, and Marblemount Hatchery.

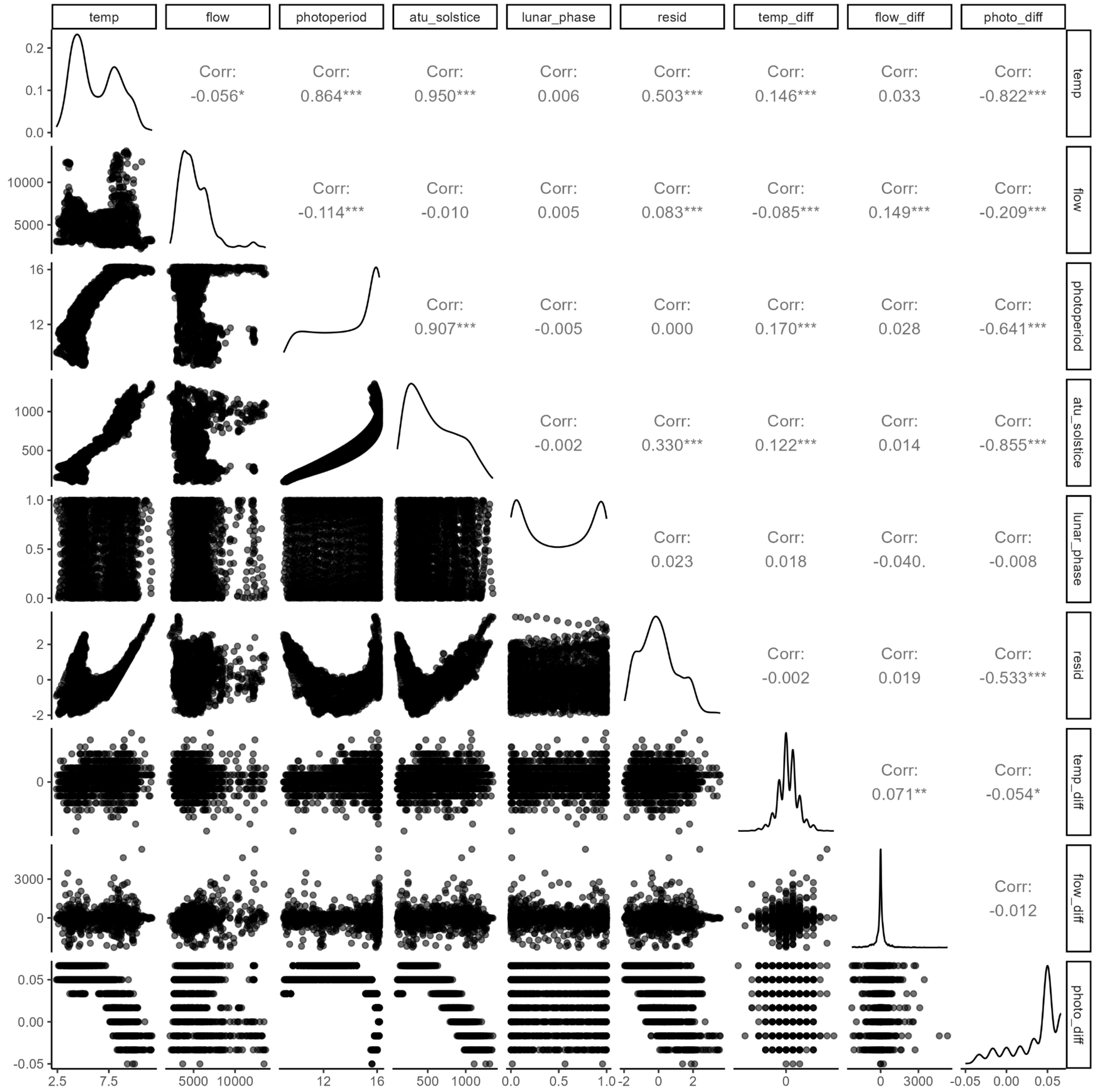

Figure S10 - Correlation between all the environmental covariates in the Skagit River.

| Chinook | | Coho | |
|---|---|---|---|
| **Covariate** | $\Delta AICc$ | **Covariate** | $\Delta AICc$ |
| Residuals | 0 | ATU | 0 |
| Photoperiod | 4.37 | Photoperiod | 6.61 |
| Temperature | 6.76 | Residuals | 7.33 |
| ATU | 8.03 | Temperature | 7.46 |
| Photo difference | 11.54 | Photo difference | 24.75 |

Table S10 - Model selection for Chinook salmon and coho salmon in the Skagit River with each of the correlated covariates.

| Chinook | | Coho | |
|---|---|---|---|
| **Variable** | **Relative Importance** | **Variable** | **Relative Importance** |
| Temperature difference | 1 | ATU | 1 |
| Residuals | 0.9 | Hatchery | 1 |
| Photoperiod | 0.8 | Flow | 0.9 |
| Hatchery | 0.4 | Lunar phase | 0.5 |
| Flow difference | 0.3 | Temperature difference | 0.4 |
| Lunar phase | 0.3 | Flow difference | 0.3 |
| Flow | 0.3 | | |

Table S11 - Relative variable importance for all variables used in the model selection process for Chinook salmon and coho salmon in the Skagit.

| Residuals | Photoperiod | Flow | Lunar phase | Temperature difference | Flow difference | Hatchery | $\Delta AICc$ | Relative Likelihood | Weights | Cumulative weights |
|---|---|---|---|---|---|---|---|---|---|---|
| 1 | 1 | 0 | 0 | 1 | 0 | 0 | 0 | 1 | 0.1 | 0.1 |
| 1 | 1 | 0 | 0 | 1 | 0 | 1 | 0.31 | 0.86 | 0.09 | 0.19 |
| 1 | 1 | 0 | 0 | 1 | 1 | 0 | 1.33 | 0.51 | 0.05 | 0.24 |
| 1 | 1 | 0 | 0 | 1 | 0 | 0 | 1.58 | 0.45 | 0.05 | 0.28 |
| 1 | 1 | 0 | 1 | 1 | 0 | 0 | 1.69 | 0.43 | 0.04 | 0.33 |
| 1 | 1 | 0 | 0 | 1 | 1 | 1 | 1.89 | 0.39 | 0.04 | 0.36 |
| 1 | 1 | 0 | 1 | 1 | 0 | 1 | 2.12 | 0.35 | 0.03 | 0.4 |
| 1 | 1 | 1 | 0 | 1 | 0 | 0 | 2.17 | 0.34 | 0.03 | 0.43 |
| 1 | 1 | 1 | 0 | 1 | 0 | 1 | 2.46 | 0.29 | 0.03 | 0.46 |
| 1 | 0 | 0 | 0 | 1 | 0 | 0 | 2.69 | 0.26 | 0.03 | 0.49 |
| 0 | 1 | 0 | 0 | 1 | 0 | 1 | 2.83 | 0.24 | 0.02 | 0.51 |
| 1 | 1 | 0 | 0 | 1 | 1 | 0 | 3.06 | 0.22 | 0.02 | 0.53 |
| 1 | 1 | 0 | 1 | 1 | 1 | 0 | 3.21 | 0.2 | 0.02 | 0.55 |
| 1 | 1 | 0 | 1 | 1 | 0 | 0 | 3.24 | 0.2 | 0.02 | 0.57 |
| 0 | 1 | 0 | 0 | 1 | 0 | 0 | 3.28 | 0.19 | 0.02 | 0.59 |
| 1 | 1 | 0 | 0 | 1 | 0 | 1 | 3.33 | 0.19 | 0.02 | 0.61 |
| 1 | 1 | 1 | 0 | 1 | 0 | 0 | 3.45 | 0.18 | 0.02 | 0.63 |

| Residuals | Photoperiod | Flow | Lunar phase | Temperature difference | Flow difference | Hatchery | $\Delta AICc$ | Relative Likelihood | Weights | Cumulative weights |
|---|---|---|---|---|---|---|---|---|---|---|
| 1 | 1 | 1 | 0 | 1 | 1 | 0 | 3.49 | 0.17 | 0.02 | 0.65 |
| 1 | 0 | 0 | 0 | 1 | 0 | 1 | 3.64 | 0.16 | 0.02 | 0.66 |
| 1 | 1 | 0 | 1 | 1 | 1 | 1 | 3.84 | 0.15 | 0.01 | 0.68 |
| 1 | 1 | 1 | 1 | 1 | 0 | 0 | 3.86 | 0.15 | 0.01 | 0.69 |
| 1 | 1 | 1 | 0 | 1 | 1 | 1 | 4.01 | 0.13 | 0.01 | 0.71 |
| 1 | 1 | 1 | 1 | 1 | 0 | 1 | 4.27 | 0.12 | 0.01 | 0.72 |
| 1 | 0 | 0 | 0 | 1 | 1 | 0 | 4.3 | 0.12 | 0.01 | 0.73 |
| 1 | 0 | 0 | 1 | 1 | 0 | 0 | 4.3 | 0.12 | 0.01 | 0.74 |
| 0 | 1 | 0 | 0 | 1 | 1 | 1 | 4.45 | 0.11 | 0.01 | 0.75 |
| 0 | 1 | 1 | 0 | 1 | 0 | 1 | 4.52 | 0.1 | 0.01 | 0.76 |
| 0 | 1 | 0 | 0 | 1 | 1 | 0 | 4.63 | 0.1 | 0.01 | 0.77 |
| 1 | 0 | 1 | 0 | 1 | 0 | 0 | 4.8 | 0.09 | 0.01 | 0.78 |
| 0 | 1 | 0 | 1 | 1 | 0 | 1 | 4.84 | 0.09 | 0.01 | 0.79 |
| 1 | 1 | 0 | 1 | 1 | 1 | 0 | 4.89 | 0.09 | 0.01 | 0.8 |
| 1 | 1 | 0 | 0 | 1 | 1 | 1 | 4.94 | 0.08 | 0.01 | 0.81 |
| 1 | 1 | 1 | 0 | 1 | 1 | 0 | 5.02 | 0.08 | 0.01 | 0.82 |
| 1 | 1 | 0 | 1 | 1 | 0 | 1 | 5.04 | 0.08 | 0.01 | 0.82 |
| 0 | 1 | 1 | 0 | 1 | 0 | 0 | 5.06 | 0.08 | 0.01 | 0.83 |

| Residuals | Photoperiod | Flow | Lunar phase | Temperature difference | Flow difference | Hatchery | $\Delta AICc$ | Relative Likelihood | Weights | Cumulative weights |
|---|---|---|---|---|---|---|---|---|---|---|
| 1 | 0 | 0 | 0 | 1 | 0 | 0 | 5.12 | 0.08 | 0.01 | 0.84 |
| 1 | 1 | 1 | 1 | 1 | 0 | 0 | 5.17 | 0.08 | 0.01 | 0.85 |
| 1 | 1 | 1 | 0 | 1 | 0 | 1 | 5.2 | 0.07 | 0.01 | 0.85 |
| 0 | 1 | 0 | 1 | 1 | 0 | 0 | 5.2 | 0.07 | 0.01 | 0.86 |
| 1 | 0 | 0 | 1 | 1 | 0 | 1 | 5.36 | 0.07 | 0.01 | 0.87 |
| 1 | 1 | 1 | 1 | 1 | 1 | 0 | 5.36 | 0.07 | 0.01 | 0.87 |
| 1 | 0 | 0 | 0 | 1 | 1 | 1 | 5.42 | 0.07 | 0.01 | 0.88 |
| 1 | 0 | 1 | 0 | 1 | 0 | 1 | 5.7 | 0.06 | 0.01 | 0.89 |
| 1 | 1 | 1 | 1 | 1 | 1 | 1 | 5.96 | 0.05 | 0.01 | 0.89 |
| 0 | 1 | 1 | 0 | 1 | 1 | 1 | 6 | 0.05 | 0 | 0.9 |
| 1 | 0 | 0 | 1 | 1 | 1 | 0 | 6.08 | 0.05 | 0 | 0.9 |
| 0 | 1 | 1 | 0 | 1 | 1 | 0 | 6.26 | 0.04 | 0 | 0.91 |
| 1 | 0 | 1 | 0 | 1 | 1 | 0 | 6.36 | 0.04 | 0 | 0.91 |
| 1 | 0 | 1 | 1 | 1 | 0 | 0 | 6.39 | 0.04 | 0 | 0.91 |
| 0 | 1 | 1 | 1 | 1 | 0 | 1 | 6.48 | 0.04 | 0 | 0.92 |
| 0 | 1 | 0 | 1 | 1 | 1 | 1 | 6.54 | 0.04 | 0 | 0.92 |
| 1 | 0 | 0 | 1 | 1 | 0 | 0 | 6.68 | 0.04 | 0 | 0.93 |
| 0 | 1 | 0 | 1 | 1 | 1 | 0 | 6.68 | 0.04 | 0 | 0.93 |

| Residuals | Photoperiod | Flow | Lunar phase | Temperature difference | Flow difference | Hatchery | $\Delta AICc$ | Relative Likelihood | Weights | Cumulative weights |
|---|---|---|---|---|---|---|---|---|---|---|
| 1 | 1 | 0 | 1 | 1 | 1 | 1 | 6.8 | 0.03 | 0 | 0.93 |
| 0 | 0 | 0 | 0 | 1 | 0 | 0 | 6.84 | 0.03 | 0 | 0.94 |
| 1 | 0 | 0 | 0 | 1 | 1 | 0 | 6.89 | 0.03 | 0 | 0.94 |
| 1 | 1 | 1 | 1 | 1 | 1 | 0 | 6.89 | 0.03 | 0 | 0.94 |
| 1 | 1 | 1 | 0 | 1 | 1 | 1 | 6.89 | 0.03 | 0 | 0.95 |
| 0 | 1 | 1 | 1 | 1 | 0 | 0 | 6.92 | 0.03 | 0 | 0.95 |
| 1 | 1 | 1 | 1 | 1 | 0 | 1 | 6.97 | 0.03 | 0 | 0.95 |
| 0 | 0 | 0 | 0 | 1 | 0 | 1 | 7.1 | 0.03 | 0 | 0.95 |
| 1 | 0 | 1 | 0 | 1 | 0 | 0 | 7.12 | 0.03 | 0 | 0.96 |
| 1 | 0 | 0 | 0 | 1 | 0 | 1 | 7.16 | 0.03 | 0 | 0.96 |
| 1 | 0 | 0 | 1 | 1 | 1 | 1 | 7.27 | 0.03 | 0 | 0.96 |
| 1 | 0 | 1 | 1 | 1 | 0 | 1 | 7.4 | 0.02 | 0 | 0.97 |
| 1 | 0 | 1 | 0 | 1 | 1 | 1 | 7.43 | 0.02 | 0 | 0.97 |
| 0 | 1 | 1 | 1 | 1 | 1 | 1 | 8.06 | 0.02 | 0 | 0.97 |
| 1 | 0 | 1 | 1 | 1 | 1 | 0 | 8.12 | 0.02 | 0 | 0.97 |
| 0 | 0 | 1 | 0 | 1 | 0 | 0 | 8.24 | 0.02 | 0 | 0.97 |
| 0 | 1 | 1 | 1 | 1 | 1 | 0 | 8.27 | 0.02 | 0 | 0.97 |
| 0 | 0 | 1 | 0 | 1 | 0 | 1 | 8.38 | 0.02 | 0 | 0.98 |

| Residuals | Photoperiod | Flow | Lunar phase | Temperature difference | Flow difference | Hatchery | $\Delta AICc$ | Relative Likelihood | Weights | Cumulative weights |
|---|---|---|---|---|---|---|---|---|---|---|
| 0 | 0 | 0 | 0 | 1 | 1 | 0 | 8.5 | 0.01 | 0 | 0.98 |
| 1 | 0 | 0 | 1 | 1 | 1 | 0 | 8.59 | 0.01 | 0 | 0.98 |
| 0 | 0 | 0 | 1 | 1 | 0 | 0 | 8.72 | 0.01 | 0 | 0.98 |
| 1 | 0 | 1 | 1 | 1 | 0 | 0 | 8.73 | 0.01 | 0 | 0.98 |
| 1 | 0 | 0 | 1 | 1 | 0 | 1 | 8.76 | 0.01 | 0 | 0.98 |
| 1 | 1 | 1 | 1 | 1 | 1 | 1 | 8.79 | 0.01 | 0 | 0.98 |
| 0 | 0 | 0 | 0 | 1 | 1 | 1 | 8.94 | 0.01 | 0 | 0.99 |
| 1 | 0 | 1 | 0 | 1 | 1 | 0 | 8.95 | 0.01 | 0 | 0.99 |
| 1 | 0 | 0 | 0 | 1 | 1 | 1 | 8.99 | 0.01 | 0 | 0.99 |
| 0 | 0 | 0 | 1 | 1 | 0 | 1 | 9.07 | 0.01 | 0 | 0.99 |
| 1 | 0 | 1 | 0 | 1 | 0 | 1 | 9.17 | 0.01 | 0 | 0.99 |
| 1 | 0 | 1 | 1 | 1 | 1 | 1 | 9.25 | 0.01 | 0 | 0.99 |
| 0 | 1 | 0 | 0 | 1 | 0 | 0 | 9.48 | 0.01 | 0 | 0.99 |
| 0 | 0 | 1 | 0 | 1 | 1 | 0 | 9.71 | 0.01 | 0 | 0.99 |
| 0 | 0 | 1 | 1 | 1 | 0 | 0 | 10.03 | 0.01 | 0 | 0.99 |
| 0 | 0 | 1 | 0 | 1 | 1 | 1 | 10.05 | 0.01 | 0 | 0.99 |
| 0 | 0 | 1 | 1 | 1 | 0 | 1 | 10.26 | 0.01 | 0 | 0.99 |
| 0 | 0 | 0 | 1 | 1 | 1 | 0 | 10.49 | 0.01 | 0 | 0.99 |

| Residuals | Photoperiod | Flow | Lunar phase | Temperature difference | Flow difference | Hatchery | $\Delta AICc$ | Relative Likelihood | Weights | Cumulative weights |
|---|---|---|---|---|---|---|---|---|---|---|
| 1 | 0 | 1 | 1 | 1 | 1 | 0 | 10.68 | 0 | 0 | 0.99 |
| 1 | 0 | 0 | 1 | 1 | 1 | 1 | 10.71 | 0 | 0 | 1 |
| 1 | 0 | 1 | 1 | 1 | 0 | 1 | 10.82 | 0 | 0 | 1 |
| 0 | 0 | 0 | 1 | 1 | 1 | 1 | 10.98 | 0 | 0 | 1 |
| 1 | 0 | 1 | 0 | 1 | 1 | 1 | 11.05 | 0 | 0 | 1 |
| 0 | 1 | 0 | 0 | 1 | 1 | 0 | 11.09 | 0 | 0 | 1 |
| 0 | 1 | 0 | 0 | 1 | 0 | 1 | 11.15 | 0 | 0 | 1 |
| 0 | 1 | 0 | 1 | 1 | 0 | 0 | 11.44 | 0 | 0 | 1 |
| 0 | 0 | 1 | 1 | 1 | 1 | 0 | 11.63 | 0 | 0 | 1 |
| 0 | 1 | 1 | 0 | 1 | 0 | 0 | 11.64 | 0 | 0 | 1 |
| 0 | 0 | 1 | 1 | 1 | 1 | 1 | 12.04 | 0 | 0 | 1 |
| 1 | 0 | 1 | 1 | 1 | 1 | 1 | 12.81 | 0 | 0 | 1 |
| 0 | 1 | 0 | 0 | 1 | 1 | 1 | 12.88 | 0 | 0 | 1 |
| 0 | 1 | 0 | 1 | 1 | 1 | 0 | 13.14 | 0 | 0 | 1 |
| 0 | 1 | 0 | 1 | 1 | 0 | 1 | 13.15 | 0 | 0 | 1 |
| 0 | 1 | 1 | 0 | 1 | 1 | 0 | 13.24 | 0 | 0 | 1 |
| 0 | 1 | 1 | 0 | 1 | 0 | 1 | 13.31 | 0 | 0 | 1 |
| 0 | 1 | 1 | 1 | 1 | 0 | 0 | 13.6 | 0 | 0 | 1 |

| Residuals | Photoperiod | Flow | Lunar phase | Temperature difference | Flow difference | Hatchery | $\Delta AICc$ | Relative Likelihood | Weights | Cumulative weights |
|---|---|---|---|---|---|---|---|---|---|---|
| 0 | 1 | 0 | 1 | 1 | 1 | 1 | 14.96 | 0 | 0 | 1 |
| 0 | 1 | 1 | 0 | 1 | 1 | 1 | 15.05 | 0 | 0 | 1 |
| 0 | 1 | 1 | 1 | 1 | 1 | 0 | 15.3 | 0 | 0 | 1 |
| 0 | 1 | 1 | 1 | 1 | 0 | 1 | 15.32 | 0 | 0 | 1 |
| 0 | 0 | 0 | 0 | 1 | 1 | 0 | 16.73 | 0 | 0 | 1 |
| 0 | 0 | 0 | 1 | 1 | 0 | 0 | 16.73 | 0 | 0 | 1 |
| 0 | 0 | 0 | 0 | 1 | 0 | 1 | 16.79 | 0 | 0 | 1 |
| 0 | 0 | 1 | 0 | 1 | 0 | 0 | 16.92 | 0 | 0 | 1 |
| 0 | 1 | 1 | 1 | 1 | 1 | 1 | 17.13 | 0 | 0 | 1 |
| 0 | 0 | 0 | 1 | 1 | 1 | 0 | 18.72 | 0 | 0 | 1 |
| 0 | 0 | 0 | 1 | 1 | 0 | 1 | 18.74 | 0 | 0 | 1 |
| 0 | 0 | 0 | 0 | 1 | 1 | 1 | 18.76 | 0 | 0 | 1 |
| 0 | 0 | 1 | 0 | 1 | 1 | 0 | 18.81 | 0 | 0 | 1 |
| 0 | 0 | 1 | 1 | 1 | 0 | 0 | 18.82 | 0 | 0 | 1 |
| 0 | 0 | 1 | 0 | 1 | 0 | 1 | 18.91 | 0 | 0 | 1 |
| 0 | 0 | 0 | 1 | 1 | 1 | 1 | 20.77 | 0 | 0 | 1 |
| 0 | 0 | 1 | 1 | 1 | 1 | 0 | 20.79 | 0 | 0 | 1 |
| 0 | 0 | 1 | 1 | 1 | 0 | 1 | 20.84 | 0 | 0 | 1 |

| Residuals | Photoperiod | Flow | Lunar phase | Temperature difference | Flow difference | Hatchery | $\Delta AICc$ | Relative Likelihood | Weights | Cumulative weights |
|---|---|---|---|---|---|---|---|---|---|---|
| 0 | 0 | 1 | 0 | 1 | 1 | 1 | 20.85 | 0 | 0 | 1 |
| 0 | 0 | 1 | 1 | 1 | 1 | 1 | 22.85 | 0 | 0 | 1 |

Table S12 - Model selection with all combinations of uncorrelated covariates for Chinook salmon in the Skagit River.

| ATU | Flow | Lunar phase | Temperature difference | Flow difference | Hatchery | $\Delta AICc$ | Relative Likelihood | Weights | Cumulative weights |
|---|---|---|---|---|---|---|---|---|---|
| 1 | 1 | 1 | 0 | 0 | 1 | 0 | 1 | 0.16 | 0.16 |
| 1 | 1 | 0 | 0 | 0 | 1 | 0.1 | 0.95 | 0.15 | 0.31 |
| 1 | 0 | 1 | 0 | 0 | 1 | 1.32 | 0.52 | 0.08 | 0.39 |
| 1 | 0 | 0 | 0 | 0 | 1 | 1.41 | 0.49 | 0.08 | 0.47 |
| 1 | 1 | 1 | 1 | 0 | 1 | 1.41 | 0.49 | 0.08 | 0.55 |
| 1 | 1 | 1 | 0 | 1 | 1 | 1.78 | 0.41 | 0.07 | 0.62 |
| 1 | 1 | 0 | 1 | 0 | 1 | 1.79 | 0.41 | 0.07 | 0.69 |
| 1 | 1 | 0 | 0 | 1 | 1 | 1.9 | 0.39 | 0.06 | 0.75 |
| 1 | 0 | 1 | 0 | 1 | 1 | 2.65 | 0.27 | 0.04 | 0.79 |
| 1 | 0 | 1 | 1 | 0 | 1 | 2.76 | 0.25 | 0.04 | 0.83 |
| 1 | 0 | 0 | 0 | 1 | 1 | 2.79 | 0.25 | 0.04 | 0.87 |
| 1 | 0 | 0 | 1 | 0 | 1 | 3.13 | 0.21 | 0.03 | 0.9 |
| 1 | 1 | 1 | 1 | 1 | 1 | 3.18 | 0.2 | 0.03 | 0.93 |

| ATU | Flow | Lunar phase | Temperature difference | Flow difference | Hatchery | $\Delta AICc$ | Relative Likelihood | Weights | Cumulative weights |
|---|---|---|---|---|---|---|---|---|---|
| 1 | 1 | 0 | 1 | 1 | 1 | 3.59 | 0.17 | 0.03 | 0.96 |
| 1 | 0 | 1 | 1 | 1 | 1 | 4.07 | 0.13 | 0.02 | 0.98 |
| 1 | 0 | 0 | 1 | 1 | 1 | 4.49 | 0.11 | 0.02 | 1 |
| 1 | 1 | 0 | 0 | 0 | 0 | 44.84 | 0 | 0 | 1 |
| 1 | 1 | 0 | 1 | 0 | 0 | 45.61 | 0 | 0 | 1 |
| 1 | 1 | 0 | 0 | 1 | 0 | 46.65 | 0 | 0 | 1 |
| 1 | 1 | 1 | 0 | 0 | 0 | 46.66 | 0 | 0 | 1 |
| 1 | 1 | 0 | 1 | 1 | 0 | 47.42 | 0 | 0 | 1 |
| 1 | 1 | 1 | 1 | 0 | 0 | 47.53 | 0 | 0 | 1 |
| 1 | 1 | 1 | 0 | 1 | 0 | 48.46 | 0 | 0 | 1 |
| 1 | 0 | 0 | 0 | 0 | 0 | 48.87 | 0 | 0 | 1 |
| 1 | 1 | 1 | 1 | 1 | 0 | 49.33 | 0 | 0 | 1 |
| 1 | 0 | 0 | 1 | 0 | 0 | 49.49 | 0 | 0 | 1 |
| 1 | 0 | 0 | 0 | 1 | 0 | 50.05 | 0 | 0 | 1 |
| 1 | 0 | 0 | 1 | 1 | 0 | 50.67 | 0 | 0 | 1 |
| 1 | 0 | 1 | 0 | 0 | 0 | 50.72 | 0 | 0 | 1 |
| 1 | 0 | 1 | 1 | 0 | 0 | 51.43 | 0 | 0 | 1 |
| 1 | 0 | 1 | 0 | 1 | 0 | 51.88 | 0 | 0 | 1 |

| ATU | Flow | Lunar phase | Temperature difference | Flow difference | Hatchery | $\Delta AICc$ | Relative Likelihood | Weights | Cumulative weights |
|---|---|---|---|---|---|---|---|---|---|
| 1 | 0 | 1 | 1 | 1 | 0 | 52.6 | 0 | 0 | 1 |
| 0 | 0 | 0 | 0 | 0 | 1 | 55.78 | 0 | 0 | 1 |
| 0 | 0 | 1 | 0 | 0 | 1 | 56.07 | 0 | 0 | 1 |
| 0 | 0 | 0 | 0 | 1 | 1 | 56.49 | 0 | 0 | 1 |
| 0 | 1 | 0 | 0 | 0 | 1 | 56.52 | 0 | 0 | 1 |
| 0 | 0 | 1 | 0 | 1 | 1 | 56.75 | 0 | 0 | 1 |
| 0 | 1 | 1 | 0 | 0 | 1 | 56.81 | 0 | 0 | 1 |
| 0 | 0 | 0 | 1 | 0 | 1 | 57.34 | 0 | 0 | 1 |
| 0 | 0 | 1 | 1 | 0 | 1 | 57.37 | 0 | 0 | 1 |
| 0 | 1 | 0 | 0 | 1 | 1 | 57.68 | 0 | 0 | 1 |
| 0 | 1 | 1 | 0 | 1 | 1 | 57.94 | 0 | 0 | 1 |
| 0 | 0 | 1 | 1 | 1 | 1 | 58 | 0 | 0 | 1 |
| 0 | 0 | 0 | 1 | 1 | 1 | 58.02 | 0 | 0 | 1 |
| 0 | 1 | 0 | 1 | 0 | 1 | 58.05 | 0 | 0 | 1 |
| 0 | 1 | 1 | 1 | 0 | 1 | 58.06 | 0 | 0 | 1 |
| 0 | 1 | 1 | 1 | 1 | 1 | 59.17 | 0 | 0 | 1 |
| 0 | 1 | 0 | 1 | 1 | 1 | 59.19 | 0 | 0 | 1 |
| 0 | 1 | 0 | 0 | 0 | 0 | 88.98 | 0 | 0 | 1 |

| ATU | Flow | Lunar phase | Temperature difference | Flow difference | Hatchery | $\Delta AICc$ | Relative Likelihood | Weights | Cumulative weights |
|---|---|---|---|---|---|---|---|---|---|
| 0 | 1 | 0 | 0 | 1 | 0 | 90.39 | 0 | 0 | 1 |
| 0 | 1 | 1 | 0 | 0 | 0 | 90.69 | 0 | 0 | 1 |
| 0 | 1 | 0 | 1 | 0 | 0 | 90.7 | 0 | 0 | 1 |
| 0 | 0 | 0 | 0 | 1 | 0 | 90.86 | 0 | 0 | 1 |
| 0 | 0 | 0 | 1 | 0 | 0 | 91.72 | 0 | 0 | 1 |
| 0 | 0 | 1 | 0 | 0 | 0 | 91.85 | 0 | 0 | 1 |
| 0 | 1 | 1 | 0 | 1 | 0 | 92.09 | 0 | 0 | 1 |
| 0 | 1 | 0 | 1 | 1 | 0 | 92.1 | 0 | 0 | 1 |
| 0 | 1 | 1 | 1 | 0 | 0 | 92.47 | 0 | 0 | 1 |
| 0 | 0 | 0 | 1 | 1 | 0 | 92.48 | 0 | 0 | 1 |
| 0 | 0 | 1 | 0 | 1 | 0 | 92.58 | 0 | 0 | 1 |
| 0 | 0 | 1 | 1 | 0 | 0 | 93.53 | 0 | 0 | 1 |
| 0 | 1 | 1 | 1 | 1 | 0 | 93.87 | 0 | 0 | 1 |
| 0 | 0 | 1 | 1 | 1 | 0 | 94.27 | 0 | 0 | 1 |

Table S13 - Model selection with all combinations of uncorrelated covariates for coho salmon in the Skagit River.

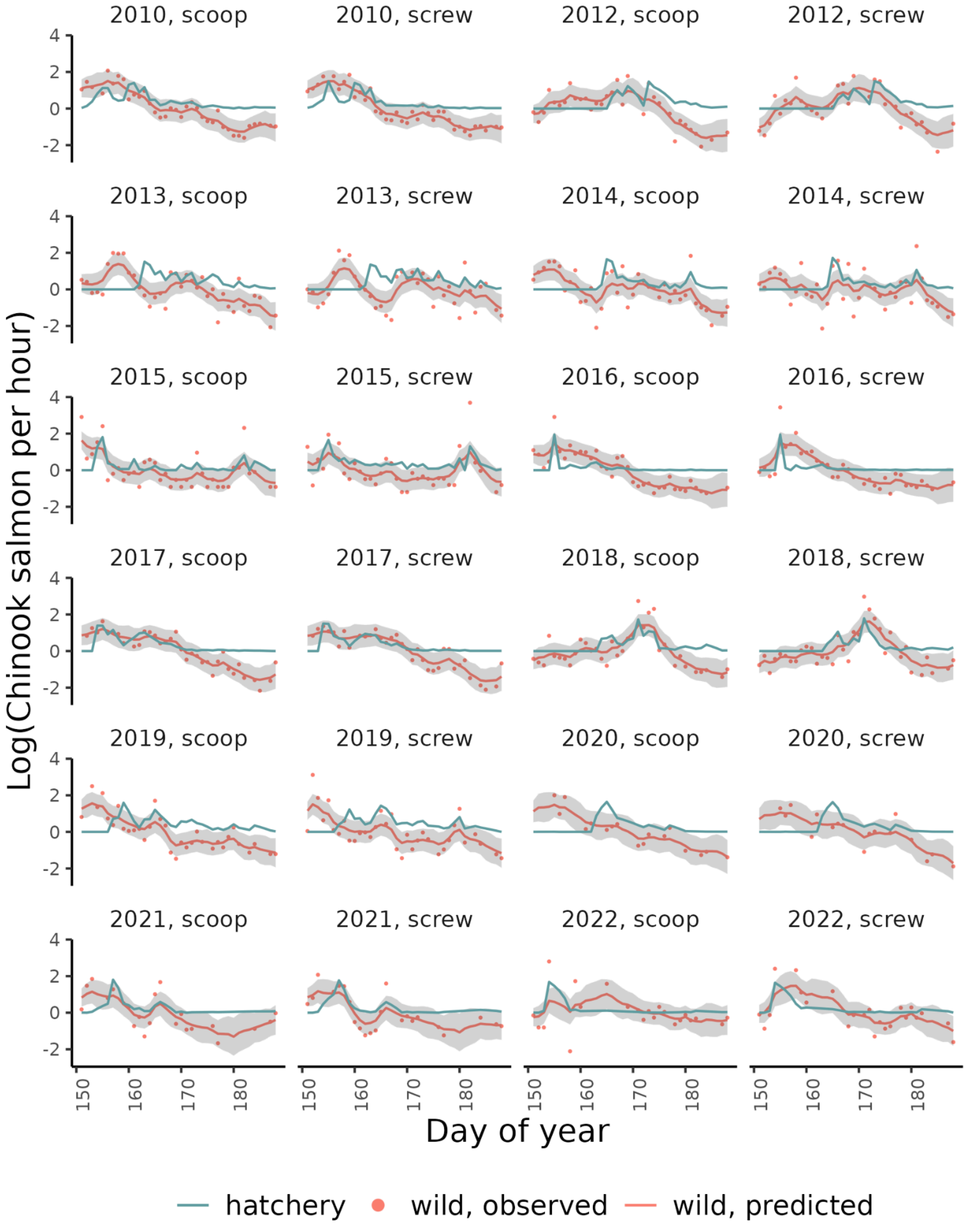


Figure S11 - Model estimates and observations of wild Chinook salmon and observations of hatchery Chinook salmon in the Skagit River.

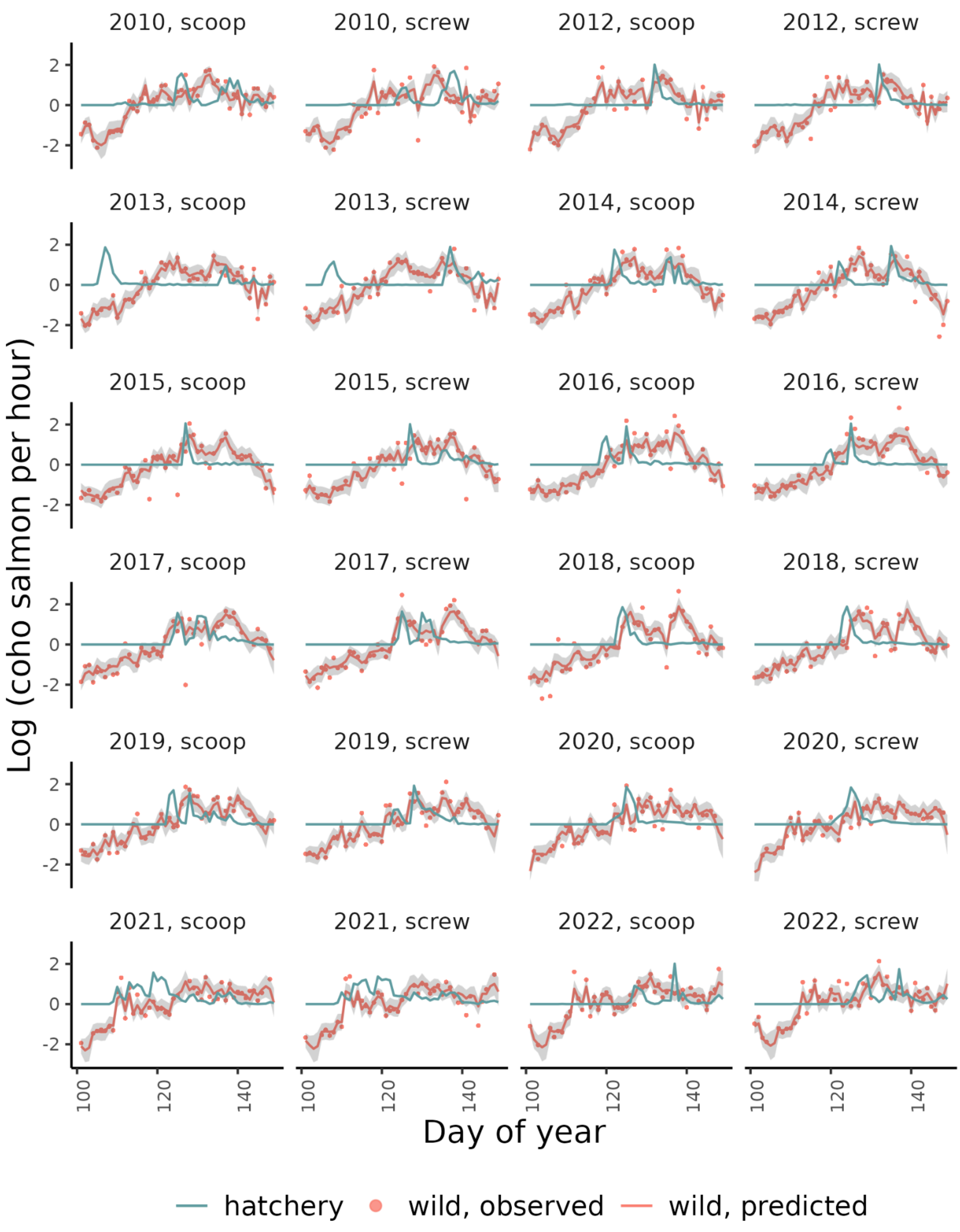

Figure S12 - Model estimates and observations of wild coho salmon and observations of hatchery coho salmon in the Skagit River.

| Species | River | Average releases | Average total released | Hatcheries |
|---|---|---|---|---|
| Chinook | Dungeness | 2 | 147731 | GRAY WOLF R ACCL PD, DUNGENESS HATCHERY, UPR DUNGENESS ACC PD, HURD CR HATCHERY |
| Chinook | Puyallup | 2 | 1279570 | VOIGHTS CR HATCHERY , GREENWATER ACCLIMATION PD, COWSKULL ACCLIM POND, PUYALLUP TRIBAL HATCHERY |
| Chinook | Skagit | 2 | 848206 | MARBLEMOUNT HATCHERY |
| Coho | Dungeness | 1 | 548141 | DUNGENESS HATCHERY |
| Coho | Puyallup | 2 | 844170 | VOIGHTS CR HATCHERY , GREENWATER ACCLIMATION PD, COWSKULL ACCLIM POND, PUYALLUP TRIBAL HATCHERY |
| Coho | Skagit | 3 | 421403 | MARBLEMOUNT HATCHERY, BAKER LK HATCHERY |

Table S14 - Average number of hatchery salmon released, hatchery location names, and average number of hatchery releases for each river-species combination as reported in RMIS.
